# The contribution of diverse and stable functional connectivity edges to brain-behavior associations

**DOI:** 10.1101/2023.11.27.568848

**Authors:** Andraž Matkovič, John D. Murray, Alan Anticevic, Grega Repovš

**Author notes:** Email addresses:* (John D. Murray), (Alan Anticevic), (Grega Repovš).

## Abstract

Resting-state functional connectivity (FC) has received considerable attention in the study of brain-behavior associations. However, the low generalizability of brain-behavior studies is a common challenge due to the limited sample-to-feature ratio. In this study, we aimed to improve the generalizability of brain-behavior associations in resting-state FC by focusing on diverse and stable edges, i.e., edges that show both high between-subject and low within-subject variability. We used resting-state data from 1003 participants with multiple fMRI sessions from the Human Connectome Project to group FC edges in terms of between-subject and within-subject variability. We found that resting-state FC variability was dominated by stable individual factors. Furthermore, diverse and stable edges were primarily part of heteromodal associative networks, and we showed that diverse stable regions are associated with a domain-general cognitive core. We used canonical correlation analysis (CCA) combined with feature selection and principal component analysis (PCA) to investigate the impact of edge selection on the strength and generalizability of brain-behavior associations. Surprisingly, selection based on edge stability did not significantly affect the results, but diverse edges were more informative than uniform edges in two of the three parcellations tested. Regardless, using all edges resulted in the highest strength and generalizability of canonical correlations. Our simulations suggest that under certain circumstances a combination of feature selection and PCA could improve the generalizability of the results, depending on the sample size and the information value of the features. The lack of improvement in generalizability with selection of stable edges may be due to unreliable estimation of within-subject edge variability or because within-subject edge variability is not related to the information value of the edges for brain-behavior associations. In other words, unstable edges may be equally informative as stable ones.

## 1. Introduction

The study of the neurobiological basis of interindividual differences in behavior and neuropsychiatric diagnoses is a particularly active area of research in neuroscience. In recent years the study of brain-behavior associations has focused on resting-state functional connectivity (RSFC). Functional connectivity (FC) is defined as a statistical dependence between time series of neurophysiological signals, and reflects causal relationships between brain regions [1]. The study of resting-state functional connectivity is of particular interest for the investigation of interindividual differences because it can be easily used with any group of people (e.g. patients, children) and the results are not affected by task difficulty, task performance, or task learning effect [2].

Recently, several studies [e.g. 3, 4, 5, 6] have warned that the generalizability of brain-behavior studies linking brain structure or function to behavior is low. The generalizability of brain-behavior studies depends on numerous factors such as sample size (or, more specifically, the number of samples per feature), effect size, type of data (e.g., functional or structural MRI; cognitive tests or questionnaires), type of statistical method (multivariate or univariate), quality and length of individual-level data [4, 3, 7, 8]. The general recommendation for improving generalizability is to increase sample size. Other recommendations include increasing the signal-to-noise ratio (SNR) and reliability of individual-level data, using feature selection or feature reduction, regularization, and comparing in-sample and out-of-sample effect sizes through cross-validation [3, 9, 4, 10, 7, 11].

The aim of this study was to explore the possibility of improving the generalizability of brain-behavior associations by exploiting information about between-subject and within-subject variability in FC. Both brain and behavior can vary between and within individuals. Traditionally, neuroscience and personality psychology have focused on the study of between-subject differences, treating within-subject differences as measurement error or noise [12, 13]. In recent years, however, studies have begun to focus on within-subject differences.

Most of the variation in FC can be explained by group and individual effects, followed by individual × task interaction [14]. Regions in associative networks (frontoparietal, cingulo-opercular, dorsal and ventral attentional networks, salience network) show greater between-subject variance and lower within-subject variance compared to unimodal/sensory regions (auditory, visual, somatomotor network) [14, 15, 16]. Mueller et al. [15] have provided indirect (meta-analytic) evidence that regions with high between-subject variance (which we termed diverse vs. uniform) and low within-subject variance (which we termed stable, i.e., regions in associative networks) correspond to regions associated with individual differences in behavioral and cognitive domains.

Because measures of personality and cognition largely reflect stable traits, we hypothesized that the brain features that are most useful for predicting behavior would also be those with the highest interindividual and lowest intraindividual variability. Two lines of research are relevant to this hypothesis. First, “fingerprinting” research aims to optimize fMRI preprocessing and analysis methods with respect to individual identification. Early research has shown that the frontoparietal regions that contribute most to identifiability also have the greatest predictive power for intelligence [17]. However, recent research has challenged these findings, showing that fingerprinting and behavioral prediction involve very different parts of the functional connectome [18]. The second line of research focuses on improving test-retest reliability. Noble et al. [19] examined the relationship between test-retest reliability and behavioral utility. They found that (1) the predictive value of edges does not correlate with test-retest reliability and (2) that including the most reliable edges in the models does not improve the behavioral prediction compared to models based on edges with low reliability. Similarly, Shirer et al. [20] have shown that optimizing for group discriminability can actually decrease test-retest reliability in some cases. Therefore, optimizing for a particular measure (discriminability or test-retest reliability) does not guarantee improved behavioral prediction [12, 20, 19, 18].

While the study by Noble et al. [19] is informative for our research question, it conflates between-subject and within-subject variability into a single dimension (i.e., test-retest reliability). The intraclass coefficient (form ICC(2,1)), often used in studies of test-retest reliability, is defined as the ratio of between-subject variance to total variance [21]. Thus, it depends not only on the between-subject variance, but also on the total variance. Therefore, edges with a high ICC are not necessarily the edges with the lowest within-subject variance. For example, holding the within-subject variance and the residual variance constant, edges with higher between-subject variance will have a higher ICC. Therefore, examining edges on two separate dimensions (between-subject variance and within-subject variance), may provide better insight into the relationship between sources of FC variability and brain-behavior associations.

Our research is divided into three parts. In the first part, we examined patterns of FC variability across participants and over different time points, from run-to-run variability to variability over six months. Similar research has been conducted previously [14, 15], but these studies used small samples of extensively sampled individuals (10 participants with 10 sessions in Gratton et al. [14]; 25 participants with 5 sessions in Mueller et al. [15]). Thus, data from these studies are more suitable for detecting within-subject than between-subject variation. We complemented this research by using data from the Human Connectome Project, which included data from 1200 subjects with four sessions on two consecutive days[22]. Because the datasets differ in the time scales over which the fMRI was recorded (e.g., on consecutive days, over weeks, or over months), we compared the results with two smaller datasets in which the data were collected over different time periods (week, month).

In the second part, we tested the hypothesis that the brain features most useful for predicting behavior are also those with the highest between-subject variability and lowest within-subject variability by using canonical correlation analysis (CCA) to estimate brain-behavior associations. Specifically, we tested whether selecting edges based on their within-subject and between-subject variability can improve the generalizability of CCA. In order to improve readability, we refer to the dimensions of between-subject variability and within-subject variability as uniform-diverse and stable-unstable, respectively. Thus, edges with low between-subject variability are also called uniform edges, and edges with high between-subject variability are called diverse edges. Similarly, the edges with low within-subject variability are called stable edges, and the edges with high within-subject variability are called unstable edges. We compared CCA models based on all edges with CCA models based on selected edges. We made two predictions. First, models based on diverse stable edges will outperform models based on all edges. Second, models based on diverse stable edges will outperform models based on diverse unstable edges.

In the third part, we used simulations to further investigate the feasibility of our feature selection procedure and to examine how factors such as the ratio of informative to non-informative features and the number of samples per feature affect the in-sample and out-of-sample effect sizes.

## 2. Method

### 2.1. Datasets

We were interested in the reproducibility of edge variability patterns across different datasets. To address the research question related to edge variability, we used four different resting-state fMRI datasets whose characteristics are summarized in Table 1. All datasets consist of at least two sessions and at least two runs per session. The HCP and YaleTRT datasets have been described in detail elsewhere [see 19, 22], so we only describe the Multimodal Imaging (MMI) Ljubljana dataset in detail here.

**Table 1:** Overview of datasets.

| Dataset name | Wu-Minn HCP (1200 Subject Release) | Wu-Minn HCP Retest | MMI Ljubljana | Yale Test-Retest |
| --- | --- | --- | --- | --- |
| Original sample size | 1096 | 45 | 48 | 12 |
| Sample size after exclusion criteria (participants with at least 1 run) | 1003 | 41 | 33 (EO), 34 (EC) | 10 |
| Age in years (SD) | 28.8 (3.7) | 30.2 (3.3) | 21.6 (2.6) | 40.2 (10.9) |
| Number of females | 596 | 31 | 26 | 6 |
| Acquisition time (1 run) | 14' 33" | 14' 33" | 18' 20" | 6' |
| TR (ms) | 720 | 720 | 1000 | 1000 |
| Number of volumes | 1200 | 1200 | 1100 | 360 |
| Recordings per session | 2 × EO | 2 × EO | 1 × EO + 1 × EC | 6 × EO |
| Sessions | 2 | 4 | 3 | 3 |
| Mean time between sessions in days (SD) | 1 (0) | 1 (0) and 139.3 (69.0)* | 31.8 (14.1) | 9.4 (5.3) |
| Reference | Van Essen et al. [22] | Van Essen et al. [22] |  | Noble et al. [19] |

*Note.* EO: eyes open, EC: eyes closed, \* 1 day difference between 1st and 2nd / 3rd and 4th session, 139.3 days difference between 2nd and 3rd session

The MMI Ljubljana dataset consists of simultaneous EEG-fMRI recordings acquired in three sessions. In each session, participants underwent two resting state runs, one with eyes open and one with eyes closed, always in that order. Participants also underwent a spatial working memory task inside the scanner and behavioral tests outside the scanner. In this paper we focus only on the resting-state fMRI data. We acquired the MRI data using the Philips Achieva 3.0T TX scanner. For each participant, we acquired T1- and T2-weighted structural images (T1 and T2: 236 slices acquired in the saggital plane, field of view = 224 × 235 mm, matrix = 320 × 336, voxel size = 0.7 × 0.7 × 0.7 mm; T1: TE = 5.8 ms, TR = 12 ms, flip angle = 8^◦^; T2: TE = 394 ms, TR = 2500 ms, flip angle = 90^◦^). Brain activity was recorded using BOLD images with T2*-weighted echo-planar imaging sequence (56 slices in the axial plane, field of view = 240 × 240 mm, voxel size = 2.5 × 2.5 × 2.5 mm, matrix = 96 × 95, TR = 1000 ms, TE = 48 ms, flip angle = 62^◦^, MultiBand SENSE factor 8). The study was approved by the institutional Ethics Committee of the Faculty of Arts, University of Ljubljana and by the National Medical Ethics Committee, Ministry of Health, Republic of Slovenia.

### 2.2. fMRI data preprocessing

All datasets were preprocessed in the same way, unless otherwise noted. First, the data were preprocessed using the HCP minimal preprocessing pipelines [23]. The HCP dataset was additionally cleaned using ICAFIX [24, 25], followed by MSMAll registration [26].

To prepare the data for functional connectivity analyses, we performed additional cleaning steps using QuNex [27]. To identify frames with excessive motion, we marked any frame that was characterized by frame displacement (FD) greater than 0.3 mm or for which the frame-to-frame signal change, calculated as the intensity normalized root mean squared difference (DVARS) across all voxels, exceeded 1.2 times the DVARS median over the entire time series, including one frame before and two frames after them. We excluded runs with less than 50 % of useful frames. In the YaleTRT dataset, this criteria resulted in 9 subjects, 3 of whom had only 1–2 useful runs. Since we were interested in variability between and within subjects, we wanted to keep as many subjects and runs as possible. Therefore, for this dataset, we adjusted the criteria, such that we included frames with FC > 0.5 mm, but we kept only runs with at least 80 % (300 total) useful frames per run. This resulted in 12 subjects with at least 8 runs.

We then used linear regression to remove nuisance signals, including 6 motion correction parameters, their squared values, signals from the ventricles, white matter, whole brain, and first derivatives of the listed signals. Previously marked frames were not included in the regression but were removed before calculating functional connectivity. Next, we parcellated the data. Because the choice of parcellation can affect the final results [e.g. 28], we used three functional parcellations to ensure the generalizability of our results: Glasser’s multimodal parcellation with 360 regions (MMP1.0) [26], Yeo’s 17-network parcellation with 224 regions [29], and Schaefer’s local-global parcellation with 400 regions [30]. Only the results of Glasser’s parcellation are presented in the main text; the other results can be found in the Supplement. Finally, functional connectivity was calculated using Pearson correlation for each run separately, and the correlation values were Fisher z-transformed for all subsequent analyses.

### 2.3. Estimation of edge variability patterns

We estimated edge variability using linear mixed models (LMM) [31, 32] for each edge separately. The advantage of using LMM is that parameter estimates are improved by partial pooling, i.e., in the case of incomplete data, estimates are partially estimated from observations with complete data [33]. Each timescale of variation (e.g., runs within days, sessions on consecutive days, sessions separated by weeks/months) was modeled as a separate factor. Note that variation over consecutive days and variation over longer time periods (weeks/months) were modeled separately. In addition to time periods, we also included between-subject variation and all possible interactions in each model. All factors were modeled as random effects. To obtain unbiased parameter estimates, models were fitted using the restricted maximum likelihood procedure (ReML) [33].

An example model for HCP data written in lme4 [31] notation:

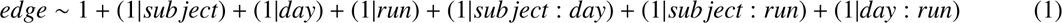

To facilitate interpretation, we averaged the edge variability results over the rows of the connectivity matrix and mapped them to the cortex. The connectivity matrices before averaging over the rows can be found in the Supplement. The HCP Retest dataset consists of participants who were also part of the HCP dataset (1200 Subjects Release), but we analyzed these two datasets separately to avoid partial pooling effects from the larger to the smaller dataset. Similarly, for the MMI Ljubljana dataset, we estimated variance components separately for eyes open and eyes closed runs. This allowed a direct comparison of the edge variability patterns between the eyes closed and eyes open conditions. If both conditions were analyzed within a single model, differences in edge variability patterns could not be detected. For example, the eyes open and eyes closed conditions could be the same in terms of fixed effects (i.e., no difference in means), but the variances (random effects) could be different.

We estimated the similarities between the edge variability patterns belonging to the different sources of variance by calculating the correlations between the patterns within each dataset.

#### 2.3.1. Similarities of edge variability patterns across studies

To estimate the reproducibility of edge variability patterns across studies, we computed the Pearson’s correlation between the variability patterns across studies for each source of variance separately.

#### 2.3.2. Similarities of edge variability patterns across sources of variability

We estimated the similarities of edge variability patterns between different sources of variance by computing the Pearson’s correlation between the variability patterns across sources of variability for each study separately.

#### 2.3.3. Stability of edge variability patterns as a function of sample size

Given the low correlations between studies for all variance sources except subjects (Figure 3), we decided to test whether the stability of edge variability patterns depends on sample size. To answer this question, we calculated the bootstrap split-half correlations at different sizes of the HCP dataset. First, we estimated the variance components for each edge on two random independent subsamples of different sizes (15, 30, 60, 125, 250, 500). Then, we correlated the edge variability patterns separately for each source of variance. We ran 500 permutations for each sample size.

**Figure 1:**
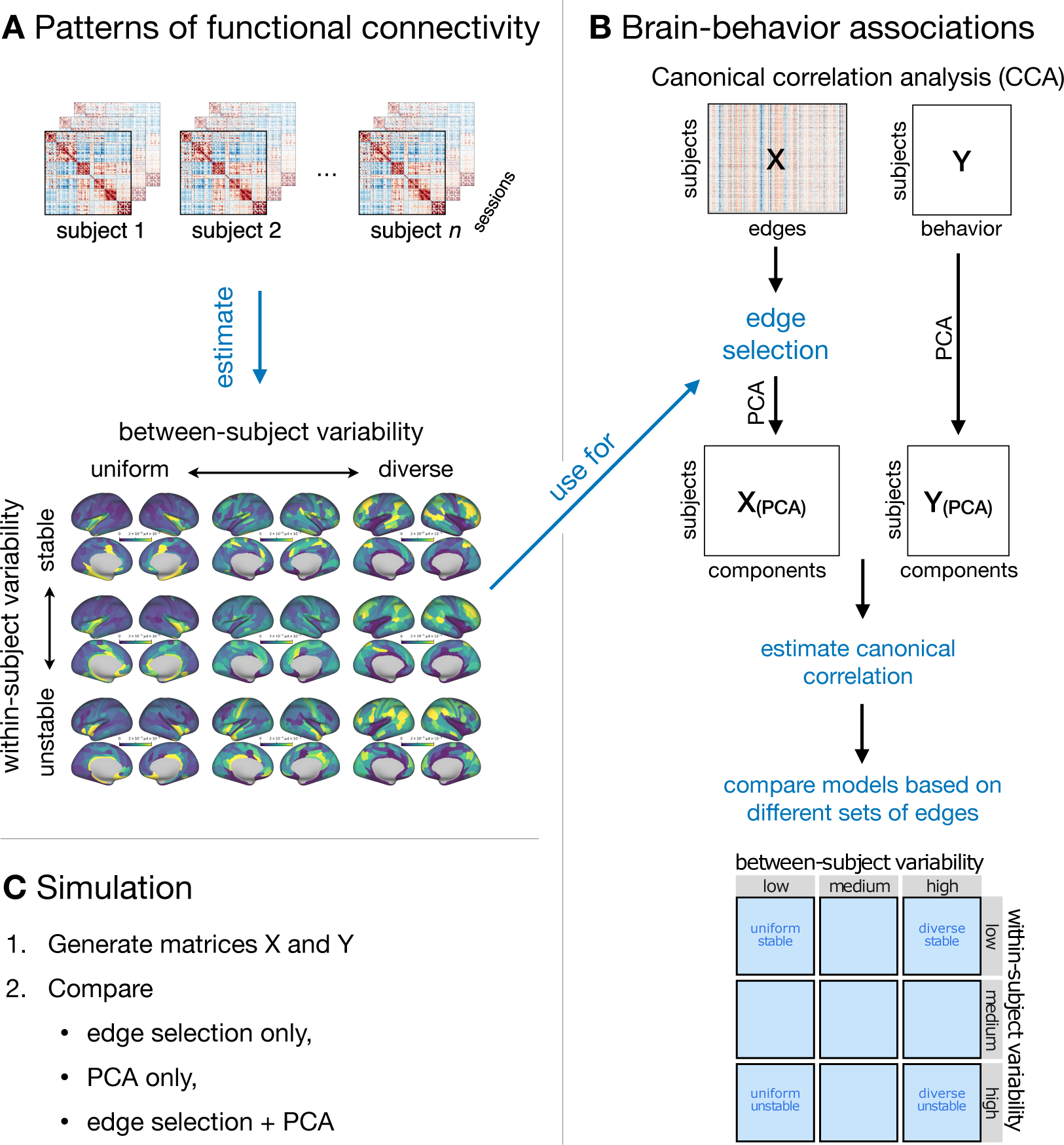
A schematic of the main steps of analysis. **A.** First, we examined patterns of within-subject and between-subject variability in functional connectivity. We defined groups of edges based on within-subject and between-subject variability. **B.** In the second part, we estimated brainbehavior associations using canonical correlation analysis (CCA). To test the hypothesis that the brain features most useful for predicting behavior are also those that are most diverse and stable, we compared models based on diverse stable edges with models based on all edges and with a model based on diverse unstable edges. **C.** In the third part, we performed a simulation in which we generated matrices X and Y with known canonical correlation. We compared feature (edge) selection, feature reduction (PCA), and a combined feature selection and feature reduction (edge selection followed by PCA). We also varied the number of features, the sample size, and the ratio of informative to non-informative features.

**Figure 2:**
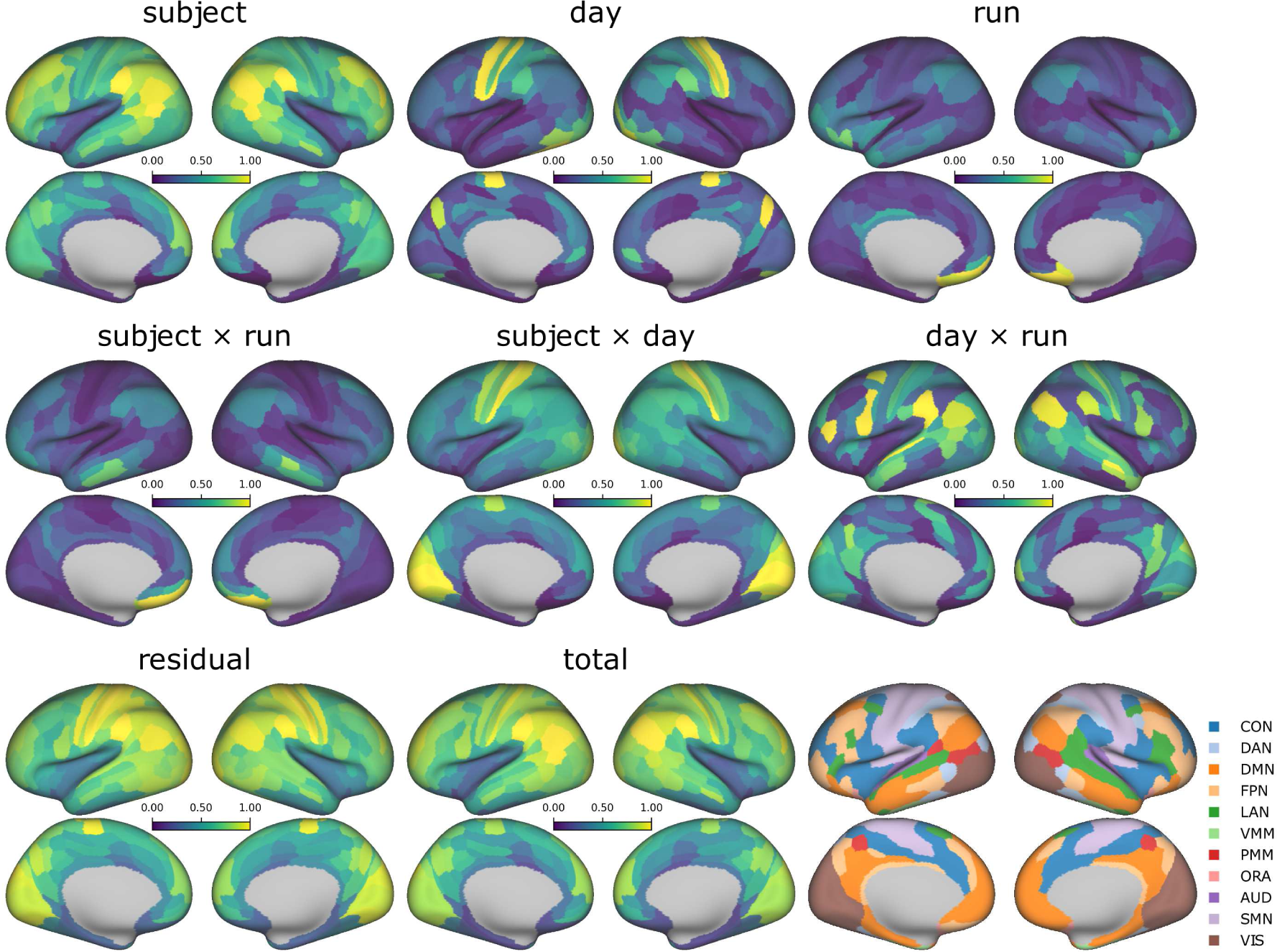
Variability of connectivity by source of variation in the HCP dataset parcellated with Glasser’s MMP1.0. Bottom right panel: Networks of Glasser’s multimodal parcellation [26] as defined by Ji et al. [37]. CON: cingulo-opercular network, DAN: dorsal attention network, DMN: default mode network, FPN: frontoparietal network, LAN: language network, VMM: ventral multimodal network, PMM: posterior multimodal network, ORA: orbito-affective network, AUD: auditory network, SMN: somatomotor network, VIS: visual network.

**Figure 3:**
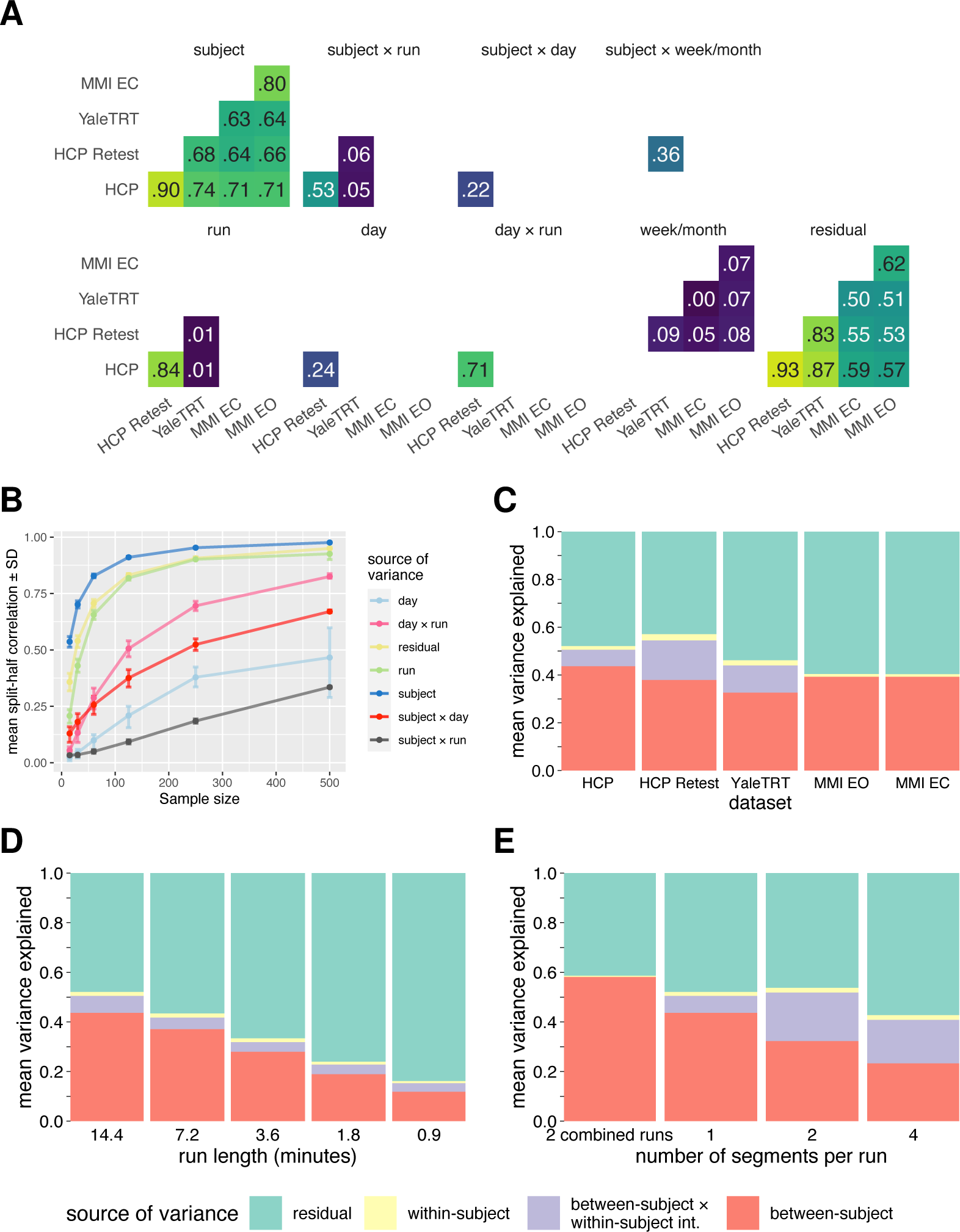
Properties of edge variability in Glasser’s multimodal parcellation. **A.** Correlations between edge variability patterns across datasets for each source of variability. **B.** Bootstrap split-half correlations between edge variability patterns as a function of sample size. The analysis was performed on the HCP dataset (1200 Subjects Release). Correlations between edge variability patterns across datasets are highest for sources of variance that are most stable at smaller sample sizes. **C.** Mean edge variance explained by each source of variability for every dataset. **D.** Mean edge variance explained as a function of run length. **E.** Mean edge variance as a function of segments per run. Note that for plots C–E, the sources of variance have been collapsed into between-subject variance, within-subject variance or interaction between these two sources. See Figure S15 for detailed plots.

#### 2.3.4. How much variance in edge variability can be explained by each source?

To estimate how much variance in edge variability can be explained by each source on average, we averaged the variances across all edges for each dataset separately (Figure 3C, Figure S15A). To facilitate comparison between datasets, the variances were normalized so that the total variance was equal to 1.

Next, to test the possibility that the variance explained by different sources might be affected by the run length or the number of runs within a session, we performed two additional analyses. First, we estimated the edge variance explained for reduced run lengths, from 14.4 minutes to 0.9 minutes (Figure 3D, Figure S15B). Second, we split the runs into 2 or 4 segments (Figure 3E, Figure S15C). In the same analysis, we also combined runs within a single day. We also tried to split the runs into 8 or more segments, but the analyses did not finish in a reasonable time.

### 2.4. Grouping of edges according to between-subject and within-subject variability

To identify the diverse stable edges (i.e., the edges with the lowest within-subject variability and the highest between-subject variability), we first calculated the total between-subject variance and the total within-subject variance for each edge. We defined between-subject variability as the sum of all subject-related variance components, excluding interactions. Within-subject variability was defined as the sum of all time-related variance components, excluding residual variance and subject × time interactions. We then divided the between-subject and within-subject variability into three groups for each dataset based on terciles. Based on this division, we defined 9 groups of edges (i.e., combinations of low, medium, or high within-subject variability and low, medium, or high between-subject variability). For visualization and analysis in the section External validation (subsection 2.5) we averaged these binary matrices across rows, resulting in a surface map of regions. These maps can be interpreted as the proportion of edges in a group that are associated with a particular region. Note that this procedure is based on ranks and therefore does not depend on the shape and location of the distribution of the edge variability.

In addition, to better understand the relationship between within-subject variability, between-subject variability, and test-retest reliability we calculated the intraclass coefficient (ICC) for each edge and plotted the ICC as a function of edge group. The ICC was estimated as the ratio of between-subject variance to total variance.

### 2.5. Similarity between edge variance maps and domain general cognitive core

When examining the surface maps of within-subject and between-subject variance, we noticed a similarity of the map of different stable regions with the map of regions of the multiple demand (MD) system [34]. This system includes regions that are co-activated during different demanding cognitive tasks, such as working memory, selective attention, and task switching [34]. Although the comparison of these maps wasn’t explicitly tied to any of the hypotheses outlined in the Introduction, their similarity implies the importance of within-subject and between-subject variability in functional connectivity for cognitive control and behavior. To quantify the similarity between the maps, we correlated the patterns of regions associated with different levels of between-subject and within-subject variability with the domain-general cognitive core defined by Assem et al. [34] (available at https://balsa.wustl.edu/study/B4nkg). In this study, the domain-general cognitive core was determined as the average of three fMRI task contrasts (2-back vs. 0-back, hard vs. easy relational reasoning, and math vs. story). MD regions were distributed across frontal, parietal, and temporal lobes.

### 2.6. Brain-behavior correlations

To test the hypothesis that the diverse stable brain features are more useful for predicting behavior than all other features, we performed a canonical correlation analysis (CCA) on the HCP dataset (1200 Subjects Release).

CCA is a multivariate statistical technique used to discover the latent structure of the shared variability between the two sets of variables. Let *X_N_*_×*P*_ and *Y_N_*_×*Q*_ be the data matrices with *N* observations and *P* or *Q* features. The goal of CCA is to solve the system of equations *U* = *XA* and *V* = *YB*, where *A_P_*_×*K*_ and *B_Q_*_×*K*_ are canonical weight matrices (*K* = *min*(*P*, *Q*)) and *U_N_*_×*K*_ and *V_N_*_×*K*_ are matrices of canonical scores. Canonical scores represent the position of each observation in the latent space, while canonical weights represent the contribution of each feature to the canonical component. The matrices of canonical scores are orthogonal, *U*^′^*U* = *V*^′^*V* = *I*. Canonical correlations are correlations between the columns of *U* and *V*.

CCA was performed (1) on all edges and (2) on edges selected on the basis of edge variability in groups with low, medium, or high within-subject and between-subject variability (as described in subsection 2.4).

We used 158 behavioral variables associated with stable traits (demographic and behavioral data) [same or similar to previous studies, e.g. 35, 10, 36, 4]. Functional connectivity and behavioral data were preprocessed in the same manner as in Smith et al. [35]. This included regression of 9 confounders (mean frame displacement, weight, height, systolic blood pressure, diastolic blood pressure, hemoglobin A1C, the cube- root of total brain volume, the cuberoot of total intracranial volume) and their demeaned and squared versions where possible. We also deconfounded age and gender.

Smith et al. [35] reduced the dimensionality of the behavioral and FC data to 100 components using PCA. Because canonical correlation and its generalizability depend on the number of samples per feature [4, 3], we included the number of principal components (PCs) as an additional variable in model comparison. We performed CCA for models containing between 10 and 100 PCs in increments of 10 PCs. Each model was evaluated using 5-fold cross-validation.

CCA and data preprocessing for CCA were performed using the GEMRR package in Python [4] (https://github.com/murraylab/gemmr).

We focused our analysis on the first canonical component. To estimate the generalizability of the models, we calculated the difference between the in-sample and out-of-sample canonical correlations. We made two comparisons, one for each prediction based on our hypothesis. First, we compared partial models (i.e., models based on selected edges) to full models (i.e., models based on all edges). Second, we compared models based on diverse stable edges with models based on diverse unstable edges. We compared the models in terms of their generalizability by calculating the difference between the in-sample and out-of-sample canonical correlations. We only directly compared models that had the same number of PCs in terms of FC and behavior.

In addition, we examined the similarity of behavioral weights between partial models based on diverse stable edges and full models, and between partial models based on diverse stable edges and partial models based on diverse unstable edges. The similarity between the weights was estimated using Pearson’s correlation. For visualization, we aggregated these correlations separately for FC and behavior.

#### 2.6.1. Control analysis

We performed a control analysis for the CCA. Since the number of edges was different in each group based on within-subject and between-subject variability, we repeated CCA on 2500 randomly selected edges in each group. The results of the control analyses are reported in the Supplement.

### 2.7. Simulation

We conducted a simulation to test the feasibility of our combined feature selection and feature reduction procedure and to examine how factors such as the ratio of informative to noninformative features and the number of samples per feature affect the in-sample and out-of-sample effect sizes. Both feature selection and feature reduction are used to reduce the feature space with the goal of improving the generalizability of a model. The difference between the procedures is that in feature selection we reduce the feature space by including only the relevant features in the model without any feature transformation, whereas in feature reduction we transform the original set of features into a lower dimensional space using techniques such as PCA.

For the simulation, we used a procedure developed by Helmer et al. [4] and implemented in the GEMMR package. Briefly, the procedure involves generating matrices *X* and *Y* with an assumed first canonical correlation. The weight matrices *A* and *B* are chosen randomly, but constrained to satisfy *A*^′^Σ*_XX_ A* = *B*^′^Σ*_YY_ B* = *I* (Σ*_XX_*and Σ*_YY_* are covariance matrices). Depending on the simulation, we created the matrix X with a sparse signal so that only a certain proportion of the features were informative. We achieved this by concatenating a matrix of informative features and a matrix of non-informative features. The informative features matrix was created using the Helmer’s procedure, while the non-informative features matrix was created by sampling from a multivariate normal distribution with uncorrelated features. Next, depending on the simulation, we performed PCA for feature reduction. Finally, we performed CCA and computed the in-sample and out-of-sample canonical correlations. The out-of-sample canonicals correlations were estimated by 5-fold cross-validation. Note that the feature selection/reduction was performed only for the matrix *X*.

We performed two simulations. In the first simulation, we generated *X* and *Y* matrices such that each contained 100 features and from 500 to 8000 observations. We set the simulated canonical correlation 0.3. The simulated canonical correlation was set at 0.3, based on a study by Helmer et al. [4] that examined 100 brain-behavior CCA analyses in 31 publications and concluded that most true canonical correlations in such studies are probably not greater than this value. We designed the *X* matrix such that 50% of the features were informative.

To evaluate the impact of feature selection and reduction, we compared three methods: feature selection, feature reduction (PCA) only, and feature selection followed by feature reduction. For feature selection, we considered three scenarios: selecting only informative features, selecting only uninformative features, or selecting an equal number of informative and uninformative features. We varied the number of selected features and principal components from 10 to 50 in increments of 10. We repeated the simulation 1000 times for each combination of manipulated variables.

In the second simulation, we generated matrices *X* and *Y* such that matrix *X* contained 1000 features, while matrix *Y* contained 100 features. The difference in the number of features between matrix *X* and matrix *Y* reflected the difference in the number of brain features (functional connectivity edges) and the number of behavioral variables. The number of observations ranged from 500 to 4000, and the canonical correlation was set to 0.3. We varied the number of informative features in *X* from 250 to 1000 in increments of 250. In the feature reduction step we selected from 100 to 500 informative features in increments of 100, followed by PCA dimensionality reduction from 10 to 50 components in increments of 10. Each simulation was repeated 100 times.

For brevity, the results of the second simulation are presented in the Supplement.

## 3. Results

### 3.1. Edge variability patterns

In this section, we focus on the results from the HCP dataset (1200 Subjects Release) parcellated using Glasser’s MMP1.0 parcellation [26]. Figures of the results on the edge variability patterns of other datasets and parcellations can be found in the Supplement.

The variability of connectivity (Figure 2) with respect to subjects was high in the associative networks, especially in the frontoparietal network (FPN), the language network, the default mode network (DMN), and the dorsal attention network (DAN). It was also high in edges within the sensory networks (visual, somatomotor), but it was low in the edges connecting the heteromodal and sensory networks (Figure S1).

Variability related to day was high in somatomotor and visual areas. Variability related to subject × day interaction resembled a similar pattern but was more widespread across the visual cortex. Variability related to run and subject × run interaction was highest in the DMN and FPN, including the edges connecting these networks.

The pattern of variability associated with the interaction between day and run was most complex, with high variability in the DMN, DAN, language, and cingulo-opercular (CON) networks, but also in the edges connecting associative and sensory cortices, particularly the connections between CON and the visual network, and between the language and the visual network.

The results were similar for all parcellations (Figure S2, Figure S4). However, in the case of Yeo’s parcellation, the clusters with high variability were generally larger, probably due to the smaller number of regions and the resulting larger average size of the regions.

#### 3.1.1. Similarities of edge variability patterns across studies

The datasets were very similar (Figure 3) only with respect to subject variability (*r* = .63–.90) and with respect to residual variability (*r* = .50–.93). Among the other sources of variability, only between the HCP (1200 Subjects Release) and HCP Retest datasets were there large similarities in terms of run, subject × run, and day × run variability. The similarities between the other datasets in terms of subject × run, week/month, and run variability were very low (*r* < .10). There was a moderate (*r* = .36) similarity in variability across subject × week/month between the Yale TRT and the HCP Retest datasets.

Results were similar for all parcellations used (Figure S10, Figure S11).

#### 3.1.2. Similarities of edge variability patterns across sources of variability

The correlations of the edge variability patterns across different variability sources were generally low, with a few exceptions (Figure S12, Figure S13, Figure S14). There was moderate to high correlation between the residual and subject variance, and between the residual and subject × day or subject × week/month variance, depending on the dataset. There were also moderate to high correlations between some terms referring to the same source of variance (e.g., subject and subject × day or subject × week/month, run and subject × run, day × run, and subject × day). The differences between parcellations were generally small, but the correlations between the sources of variance were slightly larger in the case of Yeo’s parcellation.

#### 3.1.3. Stability of edge variability patterns as a function of sample size

To explore how edge variability is affected by sample size, we performed split-half correlation analyses on subsamples of different sizes (Figure 3B). The mean bootstrap split-half correlation of edge variability patterns between pairs of subsamples increased as a function of sample size for all sources of variance. The correlations were most stable for subject variability, reaching *r* = .75 at a sample size of about 45. The patterns were most stable for run and residual variability; the correlation reached *r* = .75 at a sample size of 100. Of the other sources of variability, only the day × run variability reached *r* = .75 at *N* = 350, while the other sources did not reach *r* = .75 even at a sample size of 500. Of the remaining sources, the interaction between subject × and day was the most stable, followed by the variability between days and the variability between subject × and run.

#### 3.1.4. How much variance can be explained with each source?

For all datasets, the largest proportion of the variance across edges was explained by subject variability (32–43%, Figure 3C). This was followed by the interaction of subject with run, day, or week/month, which explained 1–10% of the variance. All other sources individually explained less than 1% of the variance.

The proportion of variance explained decreased linearly with decreasing run length for all sources of variance, from 50% for the 14.4 minute run to 15% for the 0.9 minute run (Figure 3D, Figure S15B).

The amount of variance explained was also related to the number of segments per run. When the run was split into 2 or 4 runs, the proportion of between-subject variance and total variance decreased linearly (Figure 3E). Similarly, the proportion of between-subject variance increased to 60% when runs within a single day were concatenated. On the other hand, the between-subject × within-subject interaction variance was larger when the run was split into 2 or 4 segments. However, this change was almost entirely due to subject × day or subject × run variance and not the variance between segments (Figure S15C).

### 3.2. Grouping of edges according to between-subject and within-subject variability

We classified cortical regions into nine groups according to their relative position with respect to between-subject and within-subject variability (Figure 4, Figure S16). The diverse unstable regions were distributed across the associative networks, especially DMN and CON, but also included FPN, DAN, and the language network. Diverse stable regions were located in the FPN and the posterior multimodal network. Uniform unstable regions included the FPN and the CON network. Uniform stable regions were located in the auditory network and the DMN.

**Figure 4:**
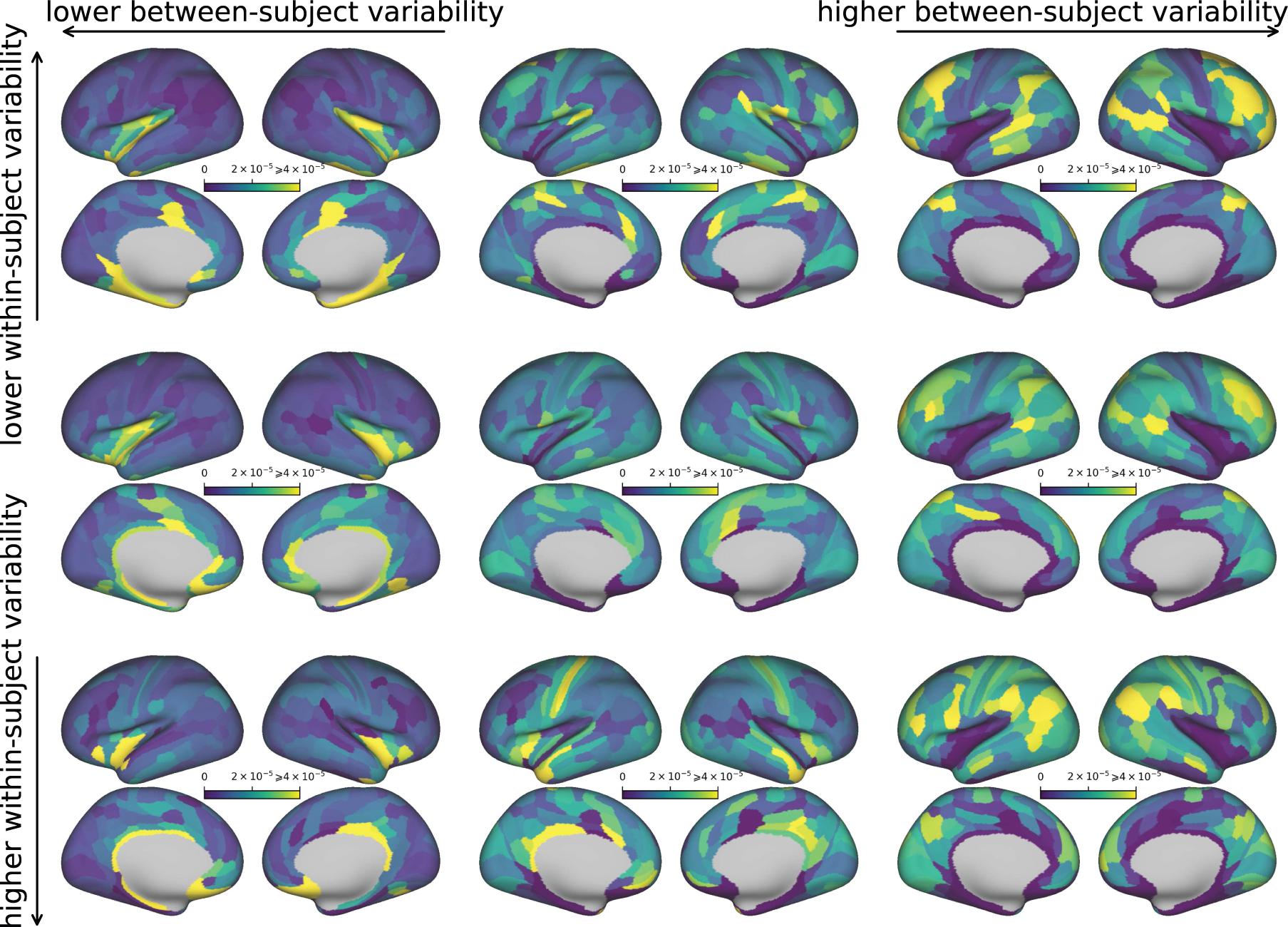
Variability of connectivity by region as a function of within-subject and between-subject variability in the HCP dataset parcellated using Glasser’s multimodal parcellation.

The analyses showed that the Glasser and Schaefer parcellations yielded similar results, while Yeo’s parcellation showed uneven distribution of edges across its nine groups. Specifically, the group of diverse stable edges had only 234 edges, and the group of uniform unstable edges had only 42 edges. Further investigation revealed that this pattern was due to a high correlation between between-subject and within-subject variance in Yeo’s parcellation (Spearman’s *ρ* = .82). On the other hand, the Glasser and Schaefer parcellations had much lower correlations between these measures (*ρ* = .47 and *ρ* = .45, respectively), resulting in a more uniform distribution of edges across groups (with a minimum of 2753 and 3753 edges in a group, respectively).

The examination of the ICC with respect to the edge grouping showed that there was a significant variability in the ICC due to the between-subject variance, while the differences due to the within-subject variance were relatively small (Figure 5, Figure S21, Figure S22). We observed the same pattern regardless of the parcellation.

**Figure 5:**
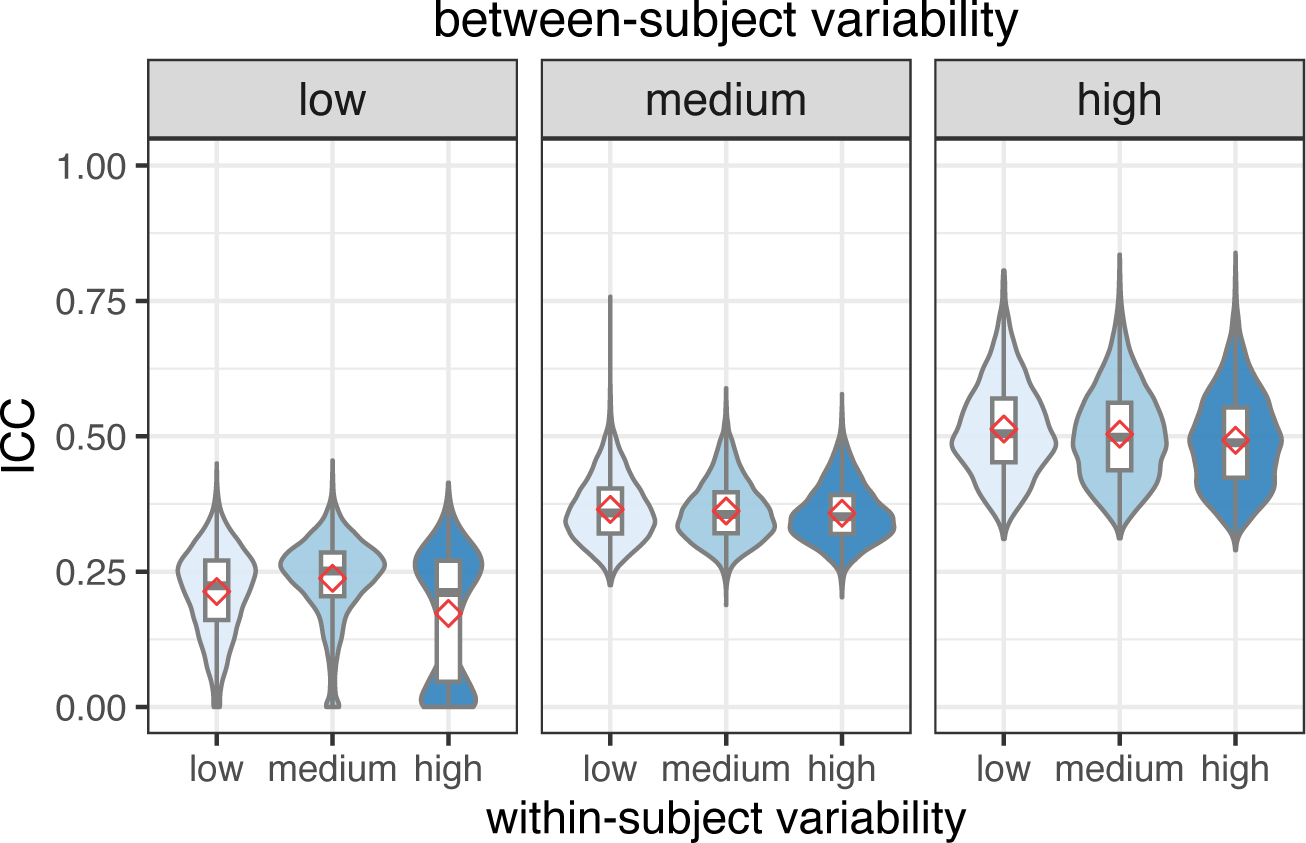
Intraclass coefficient as a function of between-subject and within-subject variability on Glasser’s parcellation.

### 3.3. Similarity between edge variance maps and domain general cognitive core

We computed the correlations between the map of domain-general cognitive core regions [34] (shown in Figure 6A) and the surface maps of regional variability in connectivity (i.e., maps from Figure 4). The correlation was largest for diverse stable edges (*r* = .45, Figure 6B) and for medium diverse stable edges (*r* = .42). Correlations for all other groups of edges were below .10 and were negative for uniform edges (*r* = −.19 for uniform stable edges; Figure S23) The results were consistent across parcellations, but the correlations were generally lower for Yeo’s parcellation (Figure S24, Figure S25).

**Figure 6:**
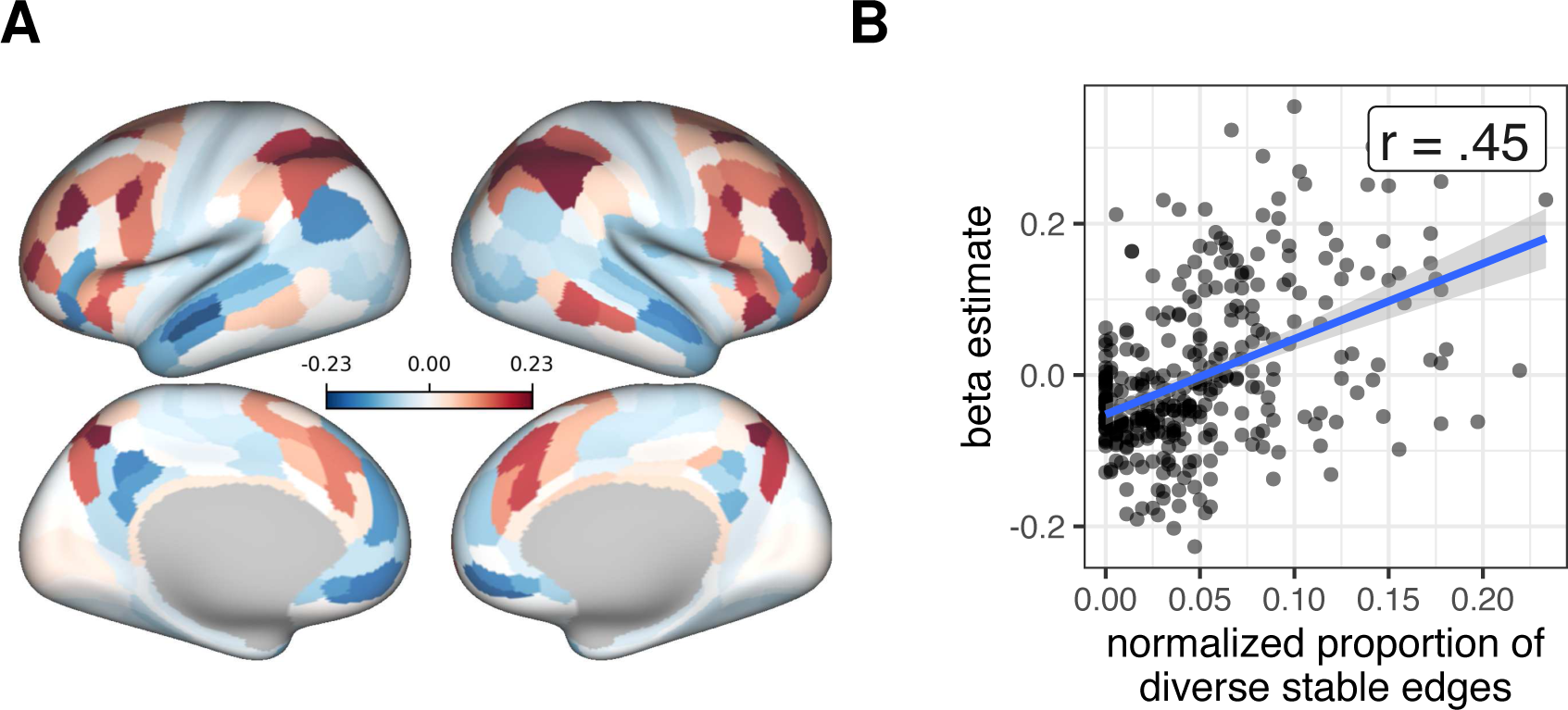
Relationship between domain-general cognitive core regions and regional variability in functional connectivity. **A.** Domain-general cognitive core regions as defined by Assem et al. [34]. **B.** Correlation between the domain-general cognitive core regions and the proportion of diverse stable edges per region (the top right map in the Figure 4). Each point represents a region. Results for other groups of regions are available in the Supplement.

### 3.4. Brain-behavior correlations

In the second part, we examined canonical correlations as a function of edge group and the number of principal components retained in the model. The in-sample canonical correlations increased as a function of the number of retained principal components related to either behavioral or functional connectivity (Figure 7A,D). The maximum canonical correlation reached about .70. We observed the same pattern regardless of the group of edges included in the model.

**Figure 7:**
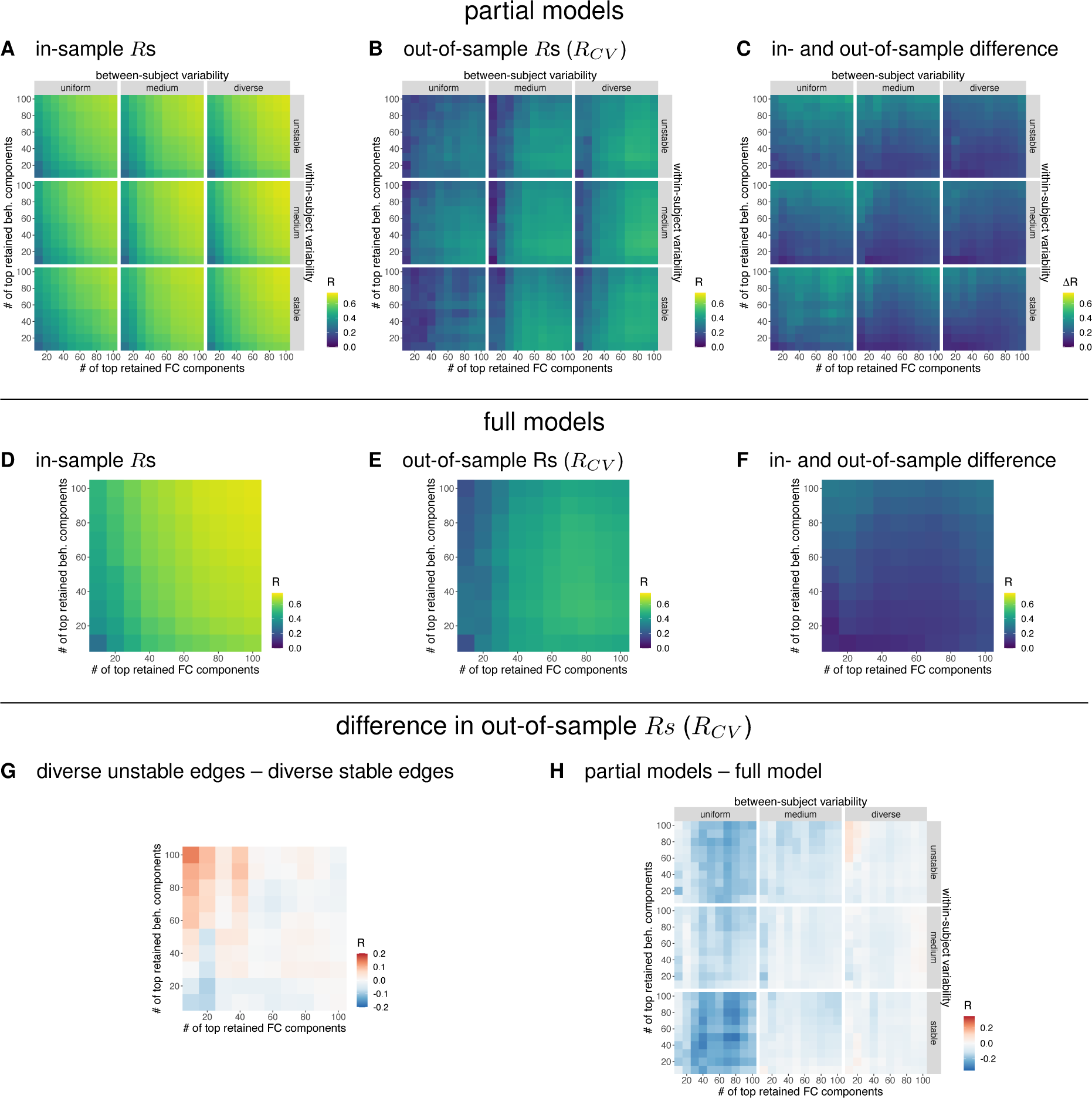
Results of CCA on Glasser’s parcellation. **A.** In-sample canonical correlations for the partial models based on groups of edges according to their between-subject and within-subject variability. **B.** Out-of-sample canonical correlations for the partial models. **C.** Difference between in-sample and out-of-sample canonical correlations for the partial models. **D.** In-sample canonical correlations for the full models. **E.** Out-of-sample canonical correlations for the full models. **F.** Difference in out-of-sample canonical correlations between full and partial models. Red color indicates higher canonical correlations for partial models. **G.** Difference between in-sample and out-of-sample canonical correlations for full models. **H.** Difference in out-of-sample canonical correlations between models based on diverse unstable edges and models based on diverse stable edges. Blue color indicates higher canonical correlations for models based on diverse stable edges.

The out-of-sample canonical correlations were lower, reaching .50 (Figure 7B,E). On average, out-of-sample canonical correlations were higher for models with a larger (70–100) number of functional connectivity components and for a medium (20–50) number of behavioral components. Out-of-sample canonical correlations were also higher for models based on diverse edges compared to models based on uniform edges.

We computed the difference between in-sample and out-of-sample canonical correlations. The differences were largest for models based on uniform edges and smallest for models based on diverse edges (Figure 7C). Among models based on diverse edges, the difference between in-sample and out-of-sample canonical correlations was smallest for models based on stable edges. For full models (i.e. models based on all edges), the difference between in-sample and out-of-sample canonical correlations was smallest when the number of functional connectivity PCs was small (< 30) and the number of behavioral PCs was medium-to-high (50–100) (Figure 7G).

Finally, we were interested in whether information about between-subject and within-subject edge variability could be used to inform models of brain-behavior correlations and to improve prediction and generalizability. We compared out-of-sample canonical correlations on full and partial models (Figure 7F). Full models outperformed partial models in most cases, especially when compared to models based on uniform edges. However, when we compared full models with partial models based on diverse stable edges, the latter had better generalizability when the number of functional connectivity PCs was small (< 30) and the number of behavioral PCs was large (50–100) (Figure 7F, top right subplot). The differences in out-of-sample canonical correlations estimates were up to .10.

We observed a similar pattern when comparing models based on diverse stable edges with models based on diverse unstable edges (Figure 7H). Models based on diverse stable edges had higher out-of-sample canonical correlations when the number of behavioral PCs was high (> 60) and the number of functional connectivity PCs was low (< 40). There were differences between the parcellations. On Schaefer’s parcellation (Figure S26), the out-of-sample canonical correlations were less dependent on the number of PCs and there were no practically important differences between partial models based on different groups of edges. In contrast, for Yeo’s parcellation (Figure S26), out-of-sample canonical correlations were highest for the partial models based on diverse unstable edges. Note, however, that in Yeo’s parcellation, the distribution of edges across groups was very uneven. We identified only 234 diverse stable edges and 42 uniform unstable edges (out of a total of 24976 edges).

#### 3.4.1. Comparison of canonical weights between the models

Next, we compared the models in terms of behavioral weight similarity. Partial models based on diverse stable edges and full models were highly similar when the number of behavioral PCs was above 30 (Figure 8A). In contrast, there was no clear relationship between the number of FC components and the similarity of the weights (Figure 8B). When comparing partial models based on uniform stable and diverse stable edges, the results were similar, except that the similarity of weights was reduced when the number of behavioral PCs was above 80 (Figure S28A). For Schaefer’s and Yeo’s parcellation, the patterns of results were generally similar, the similarity of weights plateaued at 20 behavioral PCs, but for the connectivity PCs we again observed no clear pattern (Figure S29, Figure S30).

**Figure 8:**
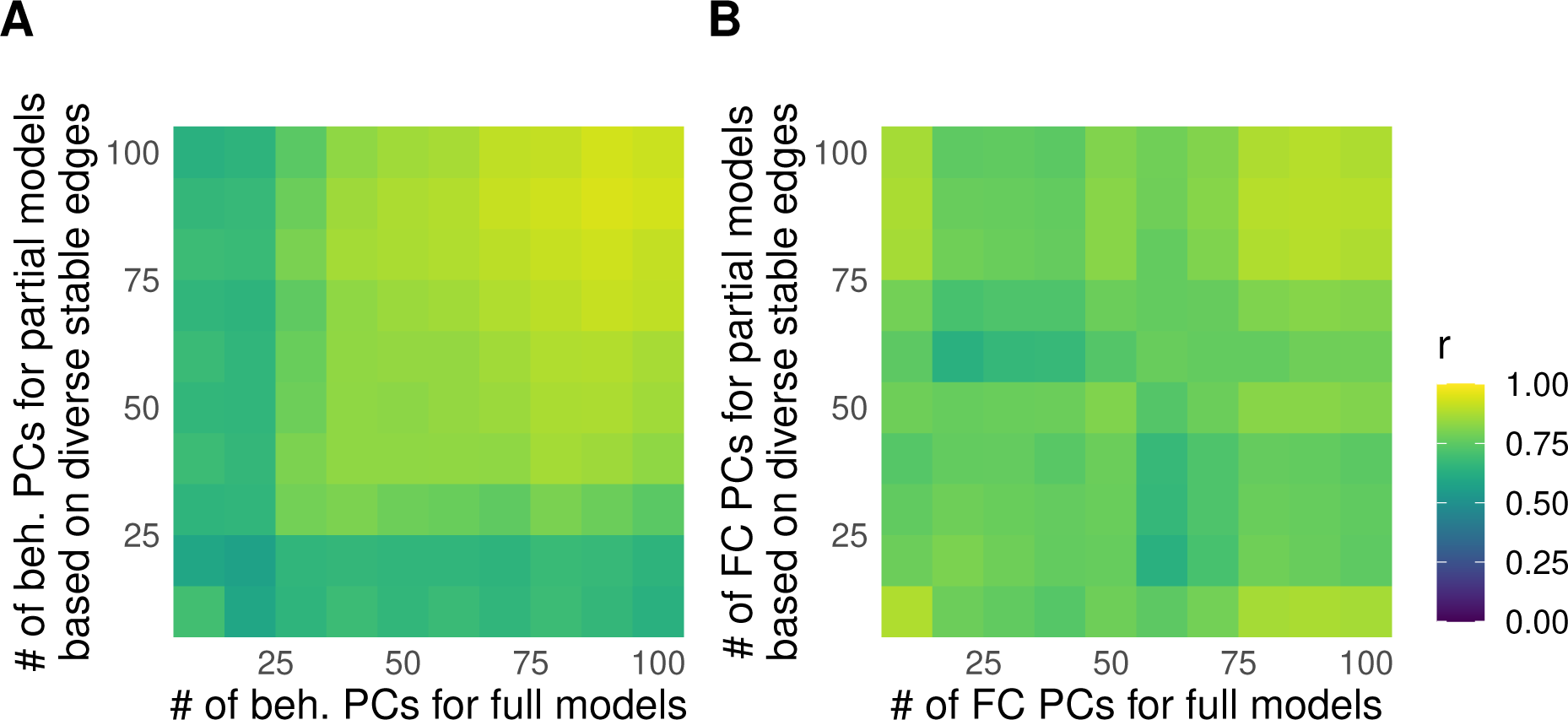
Similarity of CCA weights between full models and partial models based on diverse stable edges as a function of the number of principal components (PCs) included in the model. **A.** Similarity as a function of behavioral PCs. **B.** Similarity as a function of functional connectivity PCs. Results refer to Glasser’s parcellation.

### 3.5. Simulation

In the simulation, we examined the relationship between different feature selection or feature reduction procedures and different data properties: sample size, number of features, and proportion of informative features.

In general, in-sample canonical correlations increased as a function of the proportion of features retained, and approached true canonical correlations with increasing sample size (Figure 9A). The out-of-sample canonical correlation also increased with sample size, but only when at least some of the selected features were informative (Figure 9B).

**Figure 9:**
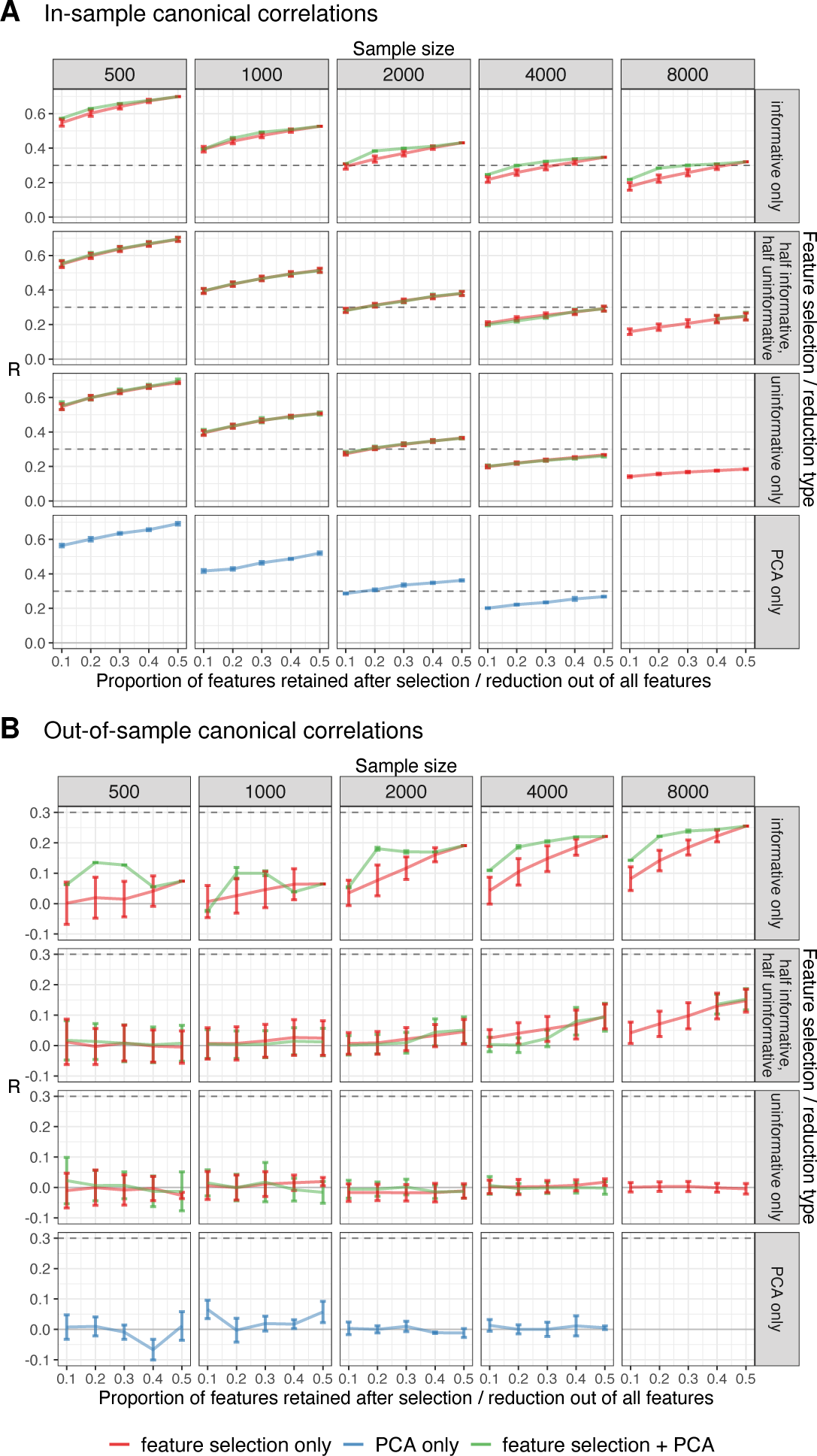
Results of the first CCA simulation. Briefly, we generated two matrices, X and Y (corresponding to functional connectivity and behavior), such that 50% of the features in X were informative. We varied the sample size, the proportion of features retained, and the type of feature selection/reduction. We used three types of feature selection/reduction: feature selection only, feature reduction (PCA only), and feature selection followed by feature reduction (feature selection + PCA). We used three schemes of feature selection, either selecting only informative features, selecting half informative and half uninformative features, and selecting only uninformative features. **A.** The in-sample canonical correlation increased with the proportion of features retained and approached the true canonical correlation with increasing sample size. **B.** Out-of-sample canonical correlations were highest when feature selection was followed by PCA, but only when the proportion of retained features was low and only informative features were selected. Some data are missing because the simulation could not be completed in time. The dashed line represents the true canonical correlation of the informative features.

In the first simulation, we compared three variants of feature selection/reduction: feature selection only, feature reduction (PCA) only, and feature selection followed by feature reduction. Compared to PCA, feature selection only was better in terms of generalizability (Figure 9B). When informative features were selected, the out-of-sample canonical correlations approached their true values as the proportion of retained features and the sample size increased. On the other hand, when only uninformative features were selected, the out-of-sample canonical correlations remained close to zero. When PCA was used for feature reduction, the out-of-sample canonical correlations remained close to zero regardless of the sample size and the number of retained PCs. When feature selection was followed by PCA, the out-of-sample canonical correlations improved compared to feature selection alone, but this improvement was only observed under certain conditions: smaller sample size, small proportion of retained features (i.e., number of PCs), and when the selected features were informative.

In the second simulation (Figure S33), we again combined feature selection and PCA, but with a larger number of features. We also varied the proportion of informative features. Feature selection and reduction improved the generalizability of CCA only when the proportion of informative features was 50%, but not when it was higher (note that we have no data for the case where the proportion of informative features was set to 25% because the simulation did not finish in time). The number of retained PCs had a negligible effect on the out-of-sample canonical correlations. The out-of-sample canonical correlations increased with the proportion of features selected before PCA.

In summary, sample size and the proportion of features retained had the greatest effect on the generalizability of CCA. Feature selection (without PCA) had a more positive effect on generalizability than PCA alone. Feature selection combined with PCA outperformed feature selection alone or PCA alone only in certain cases (small samples, small proportion of retained features, selection of informative features). Models with informative features had larger out-of-sample canonical correlations than models with uninformative features. On the other hand, the information value of the features had little effect on the in-sample canonical correlations.

## 4. Discussion

In this study, we examined the relationship between variability in functional connectivity and its relation to brain-behavior associations. FC variability across subjects was highest in associative (heteromodal) networks, whereas variability across time points was higher in unimodal networks. Regions with stable and diverse connectivity were distributed across the associative networks, particularly the frontoparietal network. Further, we showed that when comparing patterns of FC variability across subjects and across time points, edge variability patterns across subjects are stable across datasets and can be well estimated with relatively small samples. In contrast, patterns of FC variability across time points were less stable and required larger sample sizes for accurate estimation. Next, we tested the hypothesis that stable and diverse edges (i.e., edges with the lowest within-subject variability and the highest between-subject variability) would be more informative about brain-behavior associations than either edges with low between-subject variability or edges with high within-subject variability . Whereas our results indeed showed that in some parcellations diverse edges can be more informative for brain-behavior associations, in contrast to our expectations, selecting edges based on within-subject variability did not improve the generalizability of CCA. Finally, we used simulations to explore the question whether edge selection can at all improve CCA. The results indicated that appropriate selection of edges before using PCA as a dimensionality reduction step in CCA may outperform using PCA alone.

### 4.1. Associative regions exhibit variability across subjects, while unimodal regions exhibit variability across time

In the first part of the study, we examined patterns of FC variability across subjects and across different time points.

Between-subject variability in functional connectivity was highest in associative networks (fronto-parietal, language, default mode, dorsal attention network). These observed patterns of variability are consistent with previous findings [14, 15, 38] and have been linked to evolutionary cortical expansion, variations in cortical folding, and indirectly to behavioral performance [15]. They have also been associated with stable subject factors such as personality [35] and genotype [39].

Subject × time interactions were mainly influenced by variation in unimodal networks (sensorimotor, visual network). This is in contrast to Gratton et al. [14], who examined subject × session variability in a smaller dataset (Midnight Scan Club, MSC), but found no clear patterns of differences [see Figure 6 in 14]. Our results suggest that connectivity in unimodal regions, unlike connectivity in associative networks, is specific to each subject and may be more influenced by transient factors such as time in the scanner (drowsiness) [40], mood [41], amount of sleep [42], menstrual cycle phase [43], caffeine intake [44], and fasting [38].

### 4.2. The largest proportion of variance in functional connectivity can be attributed to between-subject effects

The largest amount of variability in FC (30–40% on average) was explained by between-subject effects, followed by the between subject × within-subject interaction, which explained 5–15% of the total variance. Within-subject variance was negligible. These results confirm previous findings that FC is dominated by stable group and individual factors [14, 45, 46]. In a comparable study by Gratton et al. [14], individual × session variability accounted for a very small fraction (< 5%) of total variability, while individual × task variability accounted for nearly 20% of total variability. The difference between our study and that of Gratton et al. [14] may be due to differences in the terms included in the model (time points and task). Namely, while our analysis focused solely on resting-state data, Gratton et al. [14] incorporated task-related data.

The relative variance explained by each term may also be affected by study-specific factors such as time scale (temporal distance between sessions), run length, or number of runs. By estimating variances on truncated recordings, we showed that the amount of variance explained decreases with decreasing run length. The amount of variance explained also depended on the number of segments per run – the between-subject variance decreased, but the between-subject × within-subject interaction increased when the run was divided into 2 or 4 segments. Similarly, in the YaleTRT and HCP Retest datasets, less variance was explained by between-subject variability than in the HCP (1200 Subjects Release) dataset. However, in the YaleTRT dataset, more variability was explained by the subject × day interaction compared to all other datasets. The YaleTRT dataset consists of six sessions per subject compared to 2-4 in the other datasets. In the HCP Retest dataset, the subject × day interaction explained less variance than in the HCP (1200 subjects release) dataset. The difference was likely due to more data per subject and additional terms in the model for the HCP Retest dataset (i.e., subject × day, subject × month, subject × week × day). While the exact amount of variance associated with each term depends on the particular dataset, the relative differences between terms within datasets were comparable across datasets.

### 4.3. The most stable patterns of functional connectivity are patterns of variability across subjects

Next, we showed that the pattern of between-subject FC variance is reproducible across different datasets and can be accurately estimated with sample sizes of around 50 subjects (at least when subject-level data are of similar quality as in the HCP dataset). In contrast, patterns of variability over different time points were generally less similar across studies. In the HCP dataset, these patterns also required several hundred subjects to reach stable estimates (except for variability over runs), suggesting that differences in edge variability patterns between studies are at least partly due to differences in sample size. Note that these results may be specific to the HCP dataset – with more data per subject, fewer subjects may be required to obtain stable estimates of variability over different time points.

The similarity of FC variability patterns between different time points was low. This suggests that variability over different time points is associated with different regions and may be related to different external factors. For example, variability over runs within the same day might be related to factors that change rapidly over a single session, such as fatigue [47], whereas variability over sessions recorded over weeks might be related to effects that differ between sessions, such as time of day [48, 40], affective or physiological states, recent physical activity, intake of food, liquids and psychoactive substances (nicotine, caffeine) [38], familiarity with the MR environment, etc. The low correlation between the variance patterns may also be due, at least in part, to the low stability of the variance estimates.

### 4.4. Intraclass coefficient primarily reflects between-subject variability

Studies of test-retest reliability often focus exclusively on the intraclass coefficient (ICC). We have shown that there is a large variation in the ICC as a function of the between-subject variance, while the within-subject variance has a small effect on the ICC. This is because the within-subject variance is a small proportion of the total variance compared to the between-subject variance. In other words, while the ICC combines both sources of variation into a single dimension, it predominantly reflects the between-subject variability. The within-subject variability cannot be separated from the between-subject variability when using ICC alone.

### 4.5. Diverse and stable edges are located in associative networks

We grouped the edges according to their within-subject and between-subject variability. We were particularly interested in diverse stable regions (i.e., regions with high interindividual variability and low intraindividual variability), because we expected these regions to be most informative about brain-behavior correlations. Diverse stable regions were predominantly located in heteromodal associative networks: DMN, CON, FPN, and the language network. In contrast, diverse unstable edges included both associative and unimodal networks (somatomotor, visual), whereas uniform edges (edges with low between-subject variability) included mostly auditory/limbic networks and connections between somatomotor regions and regions in heteromodal cortices.

We showed that diverse stable regions overlap with regions implicated in the domain-general cognitive core [34]. Other groups of regions showed much less overlap with the domain-general core. This is consistent with our hypothesis that these diverse stable regions are most predictive of interindividual differences in behavior. These results confirm previous findings in smaller samples [15].

These results were partially parcellation specific. While the results were similar between the Glasser and Schaefer parcellations, there were relatively few diverse stable edges when Yeo’s parcellation was used. This was due to the relatively high correlation between patterns of between-subject and within-subject variability compared to the other two parcellations. Yeo’s parcellation has fewer regions, so the parcels are on average larger and possibly more heterogeneous in terms of within-subject and between-subject variability in FC. Our results highlight the need to repeat the analysis on different parcellations, as suggested by recent reports [e.g. 28, 49, 50].

### 4.6. Brain-behavior correlations

In the second part, we tested the hypothesis that diverse and stable brain features are most useful for predicting behavior. We performed CCA on selected edges (so-called *partial* models) and on all edges (so-called *full* models).

#### 4.6.1. Data-driven approach should be used to determine the optimal number of principal components before CCA

First, we compared in-sample and out-of-sample canonical correlations. In-sample canonical correlations increased as a function of the number of features included in the models, while the relationship for out-of-sample correlations was more complex. Out-of-sample canonical correlations plateaued at about 60 principal components for FC and about 10–20 components for behavior. These results are consistent with previous reports showing that the shared variance between behavior and FC is contained in a relatively small number of PCs for behavior and a larger number of PCs for FC [4, 3]. The differences between in-sample and out-of-sample canonical correlations ranged from 0.1 to 0.4. Thus, we observed large overfitting, consistent with other similar studies [3, 4, 10]. Our results show that the optimal number of PCs for behavior or FC depends not only on the sample-to-feature ratio, but also on the information content of the features. More components would explain more variance, but including more features in the model also contributes to overfitting.

In previous studies, the number of PCs before CCA was often defined in an arbitrary way. For example, the same number of PCs was chosen for both brain traits or behaviors [35, 51, 52], or the number of PCs was chosen to explain the same proportion of variance for each modality [36]. Alternatively, the number of “significant” PCs can be determined by permutation testing for each modality separately [53, 37]. The problem with these approaches is that they do not guarantee joint variance maximization. Our and previous studies suggest that a data-driven approach should be used to select the optimal number of features when using PCA for feature reduction for CCA [10, 4, 54].

#### 4.6.2. Diverse edges may be more informative compared to uniform edges

We then compared models based on different sets of edges. Our results show that brain-behavior CCA models based on diverse and stable edges are (on average) as good as, but not better than, models based on all edges or models based on diverse unstable edges. Models based on diverse stable edges were better than full models or models based on diverse unstable edges in terms of out-of-sample canonical correlations only in very specific cases, namely for models based on a high number of behavioral PCs and a low number of connectivity PCs. Moreover, these results were specific to Glasser’s parcellation. For Schaefer’s parcellation, there were only small differences between models based on different edges. For both Glasser’s and Yeo’s parcellations, models based on different edges outperformed models based on uniform edges.

Interestingly, while we identified only 234 diverse and stable edges using Yeo’s parcellation (i.e., 0.9% out of 24976 total edges), the out-of-sample canonical correlations for this set of edges reached 0.25–0.30 (compared to 0.40 for the best model using diverse unstable edges). This suggests that even a few edges can contain a lot of behavioral information.

Including different numbers of features could also affect the content of the CCA behavioral weights. To better understand the differences and similarities between the CCA models, we correlated the behavioral weights. For behavioral PCs, the weights were very similar across models when the number of PCs was above 30 for Glasser’s parcellation and above 20 for Schaefer and Yeo’s parcellation. As before, this suggests that most of the relevant behavioral variance is captured at relatively low numbers of PCs [4]. In contrast, there was no clear pattern of weight similarity between the number of connectivity PCs for different models.

In summary, on average, full models showed equal or better out-of-sample canonical correlations than partial models. As this could be due to the fact that full models contain all the information already contained in partial models, we also compared models based on diverse stable and diverse unstable edges. Our results suggest that the information content of diverse stable edges is similar to that of diverse unstable edges. On the other hand, diverse edges were more informative than uniform edges. This effect was pronounced for the Glasser and Yeo parcellations, but not for the Schaefer parcellation. Thus, while we did not confirm our hypothesis that models based on diverse stable edges outperform full models or models based on diverse unstable edges, we did show that diverse edges have higher behavioral utility than uniform edges in some parcellations.

### 4.7. PCA alone does not necessarily select the most informative features

The inability to confirm our hypothesis does not necessarily mean that our combined feature selection and feature reduction procedure does not work in general. For example, the efficiency of our procedure may be related sample-to-feature ratio. If the sample-to-feature ratio is too low, generalizability may not be improved by any feature selection procedure, but only by increasing the sample size. On the other hand, for cases with a high sample-to-feature ratio, feature selection may not be necessary and will show comparable generalizability to models without feature selection. The feasibility of our feature selection procedure may also be affected by the ratio of informative to non-informative features. If the ratio of informative features is high, overly aggressive feature selection may exclude features that are actually informative. To address these issues, we performed a simulation in which we manipulated the sample size, the number of features, and the ratio of informative to non-informative features, as well as different combinations of feature reduction (PCA) and feature selection procedures. We reproduced the well-known finding about the overfitting of CCA at small sample sizes or small sample-to-feature ratios [4, 10]. Next, we examined the different combinations of feature selection and feature reduction procedures.

Feature selection procedure alone (without PCA) improved the generalizability of CCA when at least some of the selected features were informative. On the other hand, when PCA alone (without feature selection) was used for feature reduction, the out-of-sample canonical correlations were close to zero. The goal of PCA is to reduce the dimensionality of a dataset by exploiting correlations between variables. PCA transforms the data into a new coordinate system, and dimensionality can be reduced by keeping only the dimensions with the highest variance. However, as our simulation shows, this does not necessarily ensure that the features with the highest predictive power are retained. Importantly, using PCA, we can reduce dimensionality by removing informative components (i.e., the components that are correlated between sets) if they are not the components with the highest variance.

Next, we tested the feasibility of combining feature selection and feature reduction. This procedure improved generalizability compared to feature selection alone when the sample size was “small” (< 2000), when we retained a small proportion of features, and when the selected features were informative. Thus, when PCA is preceded by appropriate feature selection, PCA cannot retain uninformative features because they have already been removed from the data. Our procedure improved generalizability only at smaller sample sizes – as sample sizes increase, feature selection becomes less important because the sample-to-feature ratio increases and there is less room to improve generalizability. Similarly, our procedure worked best when a small proportion of features were retained. As more features are included in the model, the sample-to-feature ratio decreases and, consequently, the generalizability of the model decreases.

Our second simulation showed that feature selection and reduction improved the generalizability of CCA only when the proportion of informative features was 0.50, but not when it was higher. Thus, when a high proportion of features is informative, selecting too few features would remove the relevant information from the model.

In their simulations, Helmer et al. [4] found that the average of the in-sample and out-of-sample canonical correlations was a much better estimate of the true association strength than either the in-sample or out-of-sample correlations. Our simulations show that this is rarely the case. Even when the features are largely uninformative and the true canonical correlation is around zero, the in-sample canonical correlation can remain high even for relatively large sample sizes (e.g. up to .30 for sample size 4000 in our simulations, Figure 9). Thus, while out-of-sample canonical correlations may underestimate the true strength of the association, they still provide a more reliable measure of canonical correlations than either the in-sample canonical correlation or the average of both.

In summary, we have shown that feature reduction does not necessarily extract the most informative features, but also that the model can be improved by appropriate feature selection prior to feature reduction. However, this combination is only useful in certain circumstances. At low sample-to-feature ratios, neither feature selection nor feature reduction may improve generalizability, whereas at high sample-to-feature ratios, these procedures may not be needed.

### 4.8. Limitations and future directions

Our work has several limitations. Although we have shown by simulation that a combined feature selection and feature reduction procedure works in principle, there are several reasons why it did not work in practice. First, the patterns of variability associated with within-subject variability required large sample sizes to obtain stable estimates. Therefore, the estimated patterns of within-subject variability may not have been accurate enough, and consequently, the selection of edges based on within-subject variability may not have been accurate either. On the other hand, we have shown that the pattern of diverse stable regions correlates with the pattern of domain-general cognitive core regions, which indirectly shows that our grouping of edges based on within-subject and between-subject variability was valid. Regardless, the stability of within-subject variability patterns could be improved with more sessions per subject.

Second, we calculated within-subject variability at the group level. However, the patterns of within-subject variability may differ between subjects, and different edges may have been informative for different people. This is also indicated by the proportion of variance explained: the between-subject × within-subject interaction explained more variance than the within-subject factors alone. It would be possible to select different edges for different subjects, e.g. by first selecting a set of edges with the highest between-subject variability, and then selecting a predefined number of edges with the lowest within-subject variability for each subject separately. However, this would introduce additional complexity into the procedure, and the subsequent PCA and CCA results may not be interpretable. In addition, reliable estimates of within-subject variability at the subject level would likely require more recordings per subject.

We used PCA for dimensionality reduction prior to CCA to improve the sample-to-feature ratio. As explained earlier, this procedure does not necessarily maximize the correlation between sets. Alternatively, we could have used regularized or sparse CCA, which reduces overfitting by shrinking the weights (or forcing some weights to zero in the case of sparse CCA) [55, 56]. Regularization reduces the variance of the parameters and improves the prediction accuracy at the cost of introducing bias into the parameter estimates. The amount of shrinkage is controlled by hyperparameters, which are typically optimized by cross-validation to maximize the out-of-sample canonical correlation. Regularized CCA has already been compared to combined PCA-CCA in the context of brain-behavior correlations, and the results between the two methods were very similar [10]. This is to be expected, since the ridge penalty tends to shrink the coefficients of the low variance components more than those of the high variance components [57]. The main difference is that the ridge penalty gradually shrinks the regression coefficients of the principal components, whereas PCA truncates them after some components.

### 4.9. Conclusions

In conclusion, we showed that FC variability across subjects is highest in associative networks, whereas FC variability across time points is highest in unimodal networks. Furthermore, the variability of FC across subjects could be estimated more reliably with fewer data points than the variability across time points. FC variability across subjects was also more stable across different datasets. Diverse and stable edges were primarily located in heteromodal associative networks, and we showed that diverse stable regions are associated with a domain-general cognitive core. However, contrary to our hypothesis, diverse stable edges did not consistently outperform other edges in predicting brain-behavior associations.

Furthermore, our results suggest that resting-state functional networks are largely influenced by stable individual factors, followed by between-subject × within-subject interaction. Different time points were associated with different edges. Further studies are needed to investigate whether patterns of variability over different time points are associated with different internal (homeostatic) and external factors.

In addition, we showed that the optimal number of principal components for behavior and FC in CCA depends not only on the sample-to-feature ratio, but also on the information content of the features, suggesting that a datadriven approach should be used to optimize the number of features in CCA. Although models based on diverse stable edges performed similarly to models based on all edges, edges with higher between-subject variability were more informative than edges with lower between-subject variability in two of the three parcellations tested.

Finally, our simulations showed that combining feature selection and feature reduction can improve the generalizability of CCA under certain circumstances. The results open a new avenue for further research that could focus on other feature selection methods to improve the assessment of brain-behavior associations. More generally, these results provide valuable insights into the relationship between FC variability, brain-behavior associations, and the optimization of CCA models.

## 5. Data and code availability

The HCP raw data are available at https://www.humanconnectome.org/. The Yale Test-Retest dataset is available at http://fcon_1000.projects.nitrc.org/indi/retro/yale_trt.html. The map of general cognitive core regions [34] is available at https://balsa.wustl.edu/study/B4nkg. For CCA and simulations we used the GEMMR package: https://github.com/murraylab/gemmr. Preprocessed and parcellated data and relevant code are available in the Open Science Framework repository https://dx.doi.org/10.17605/OSF.IO/EPG6K.

## 6. Author contributions

**Andraž Matkovič:** Conceptualization, Methodology, Formal analysis, Investigation, Writing - Original Draft, Writing - Review & Editing, Visualization. **Alan Anticevic:** Conceptualization, Writing - Review & Editing. **John D. Murray**: Conceptualization, Writing - Review & Editing. **Grega Repovš:** Conceptualization, Software, Writing - Original Draft, Writing - Review & Editing, Supervision, Project administration, Funding acquisition.

## 9. Acknowledgments

Data were provided [in part] by the Human Connectome Project, WU-Minn Consortium (Principal Investigators: David Van Essen and Kamil Ugurbil; 1U54MH091657) funded by the 16 NIH Institutes and Centers that support the NIH Blueprint for Neuroscience Research; and by the McDonnell Center for Systems Neuroscience at Washington University.

## 10. Supplement

### 10.1. Variability of functional connectivity

**Figure S1:**
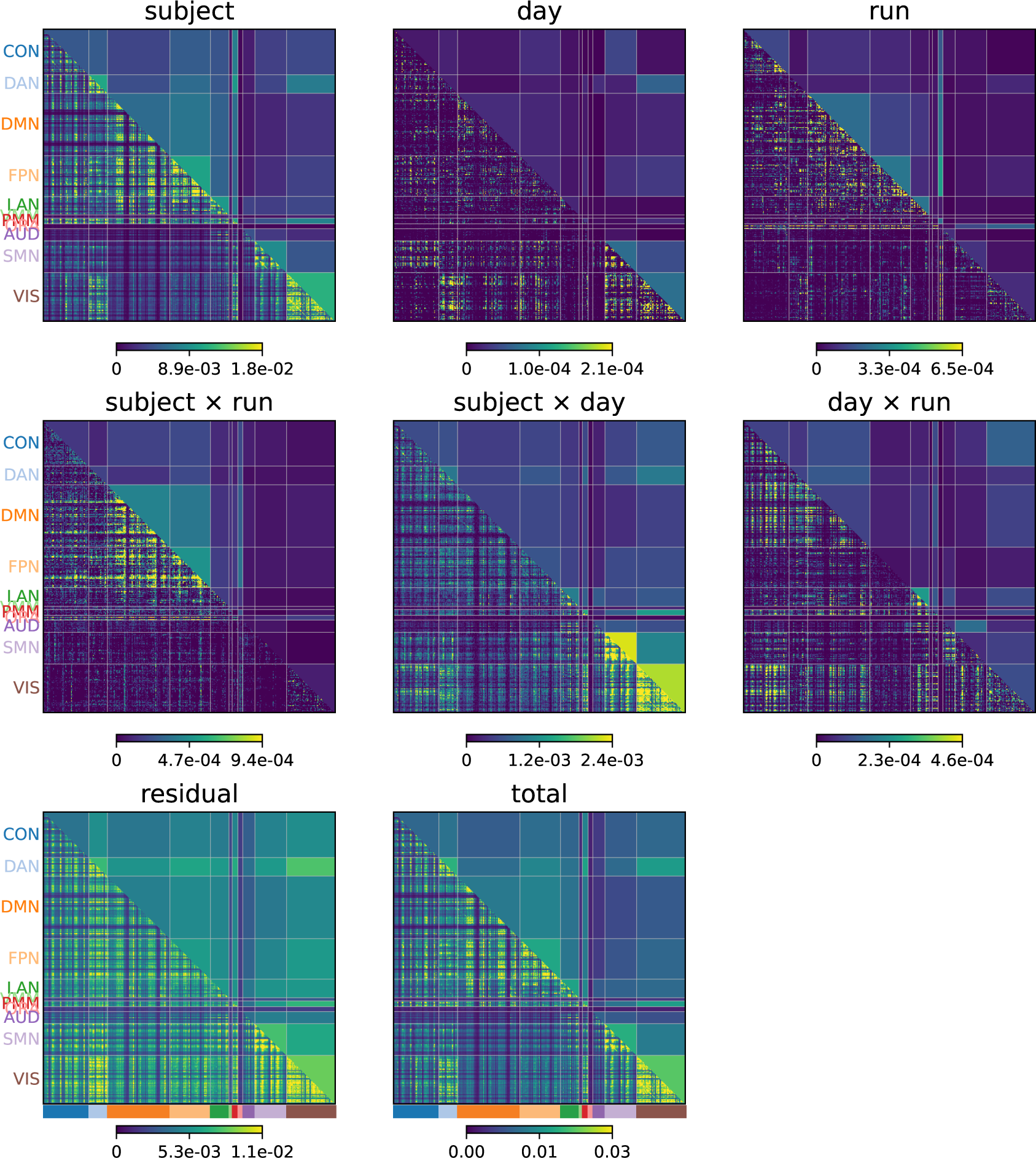
Edge connectivity on the HCP dataset parcellated with Glasser’s parcellation by source of variation. Same as in Figure 2 but not averaged by rows.

**Figure S2:**
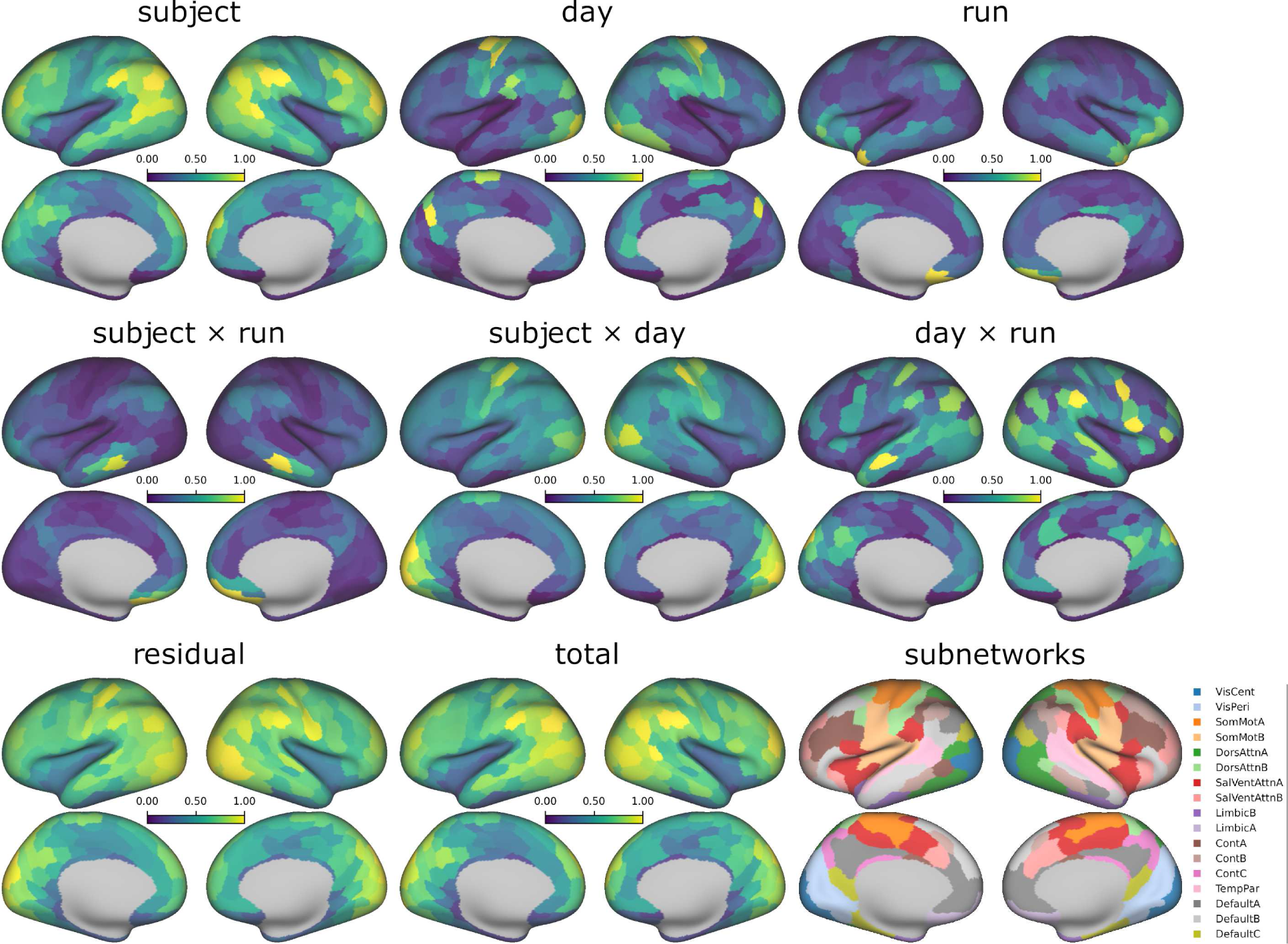
Variability of connectivity on the HCP dataset parcellated with Schaefer’s Local-Global parcellation by source of variation. Right bottom inset: Networks of Schaefer’s Local Global parcellation [30]. VisCent: visual central network, VisPeri: visual peripheral network, SomMot: somatomotor network, DorsAttn: dorsal attention network, SalVentAttn: salience / ventral attention network, Cont: control network, TempPar: temporal parietal network.

**Figure S3:**
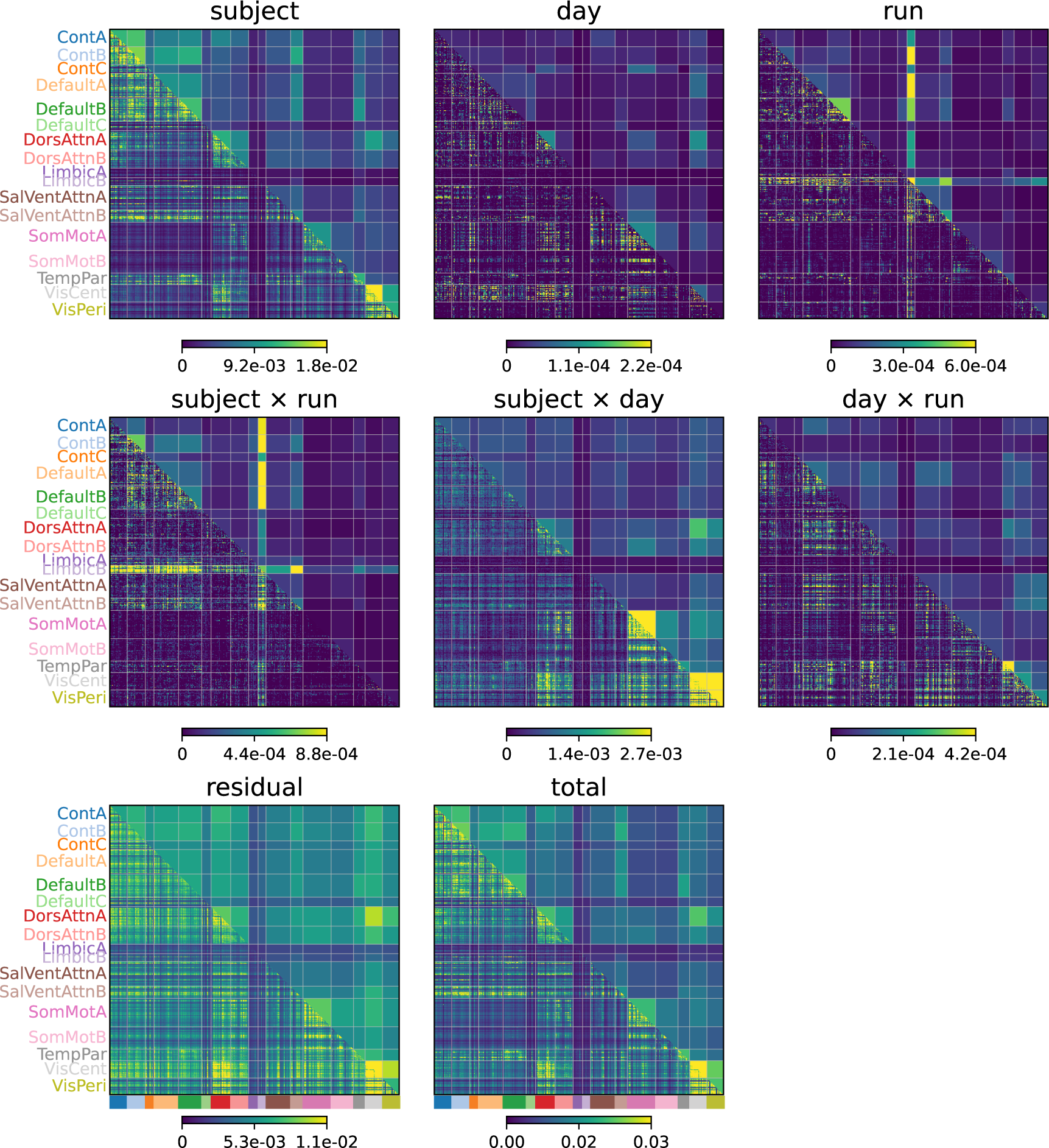
Edge connectivity on HCP dataset parcellated with Schaefer’s parcellation by source of variation. Same as in Figure S2 but not averaged by rows.

**Figure S4:**
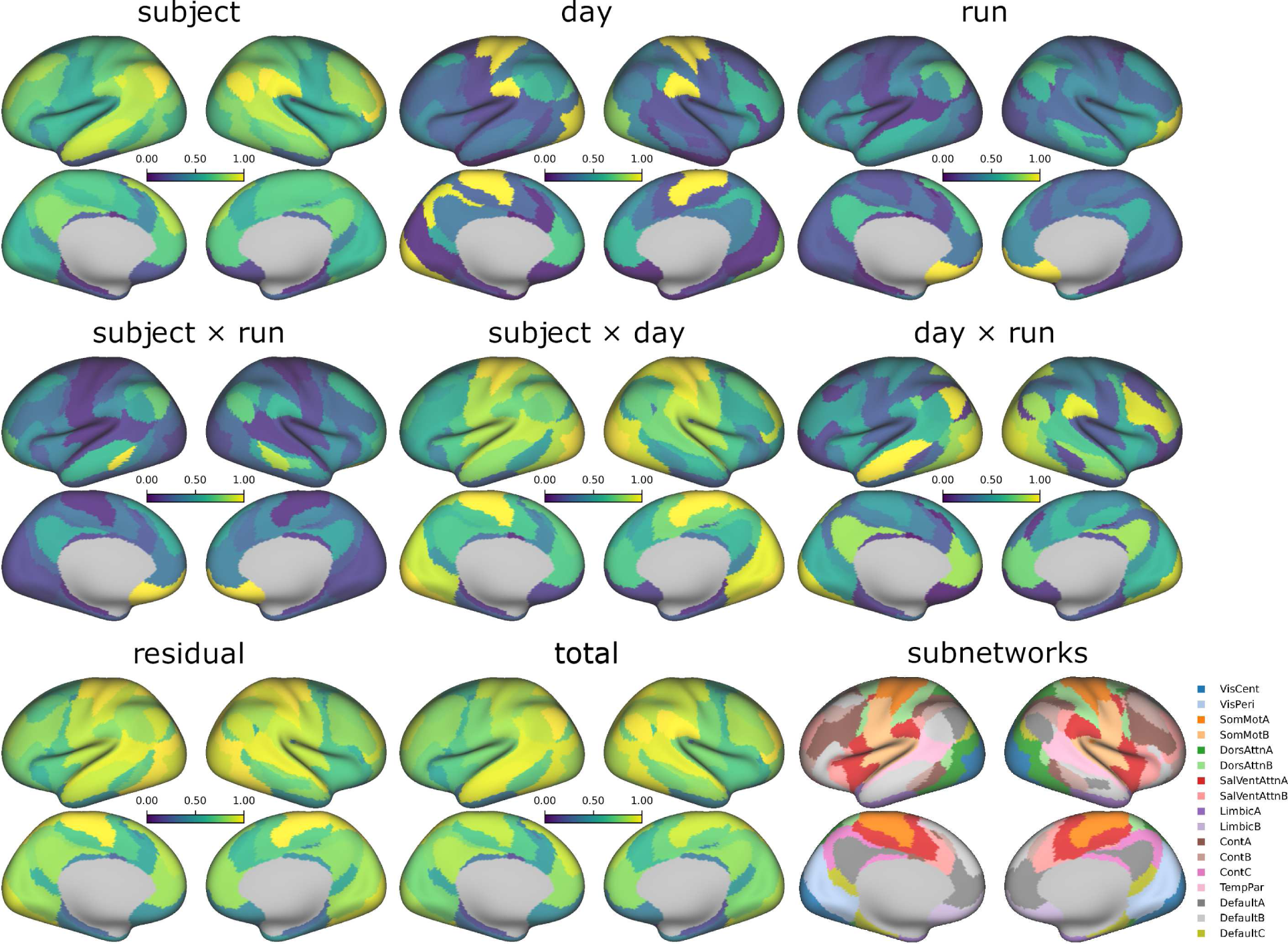
Variability of connectivity on the HCP dataset parcellated with Yeo’s 17-network parcellation by source of variation. Right bottom inset: Networks of Yeo’s 17-network parcellation [29]. VisCent: visual central network, VisPeri: visual peripheral network, SomMot: somatomotor network, DorsAttn: dorsal attention network, SalVentAttn: salience / ventral attention network, Cont: control network, TempPar: temporal parietal network.

**Figure S5:**
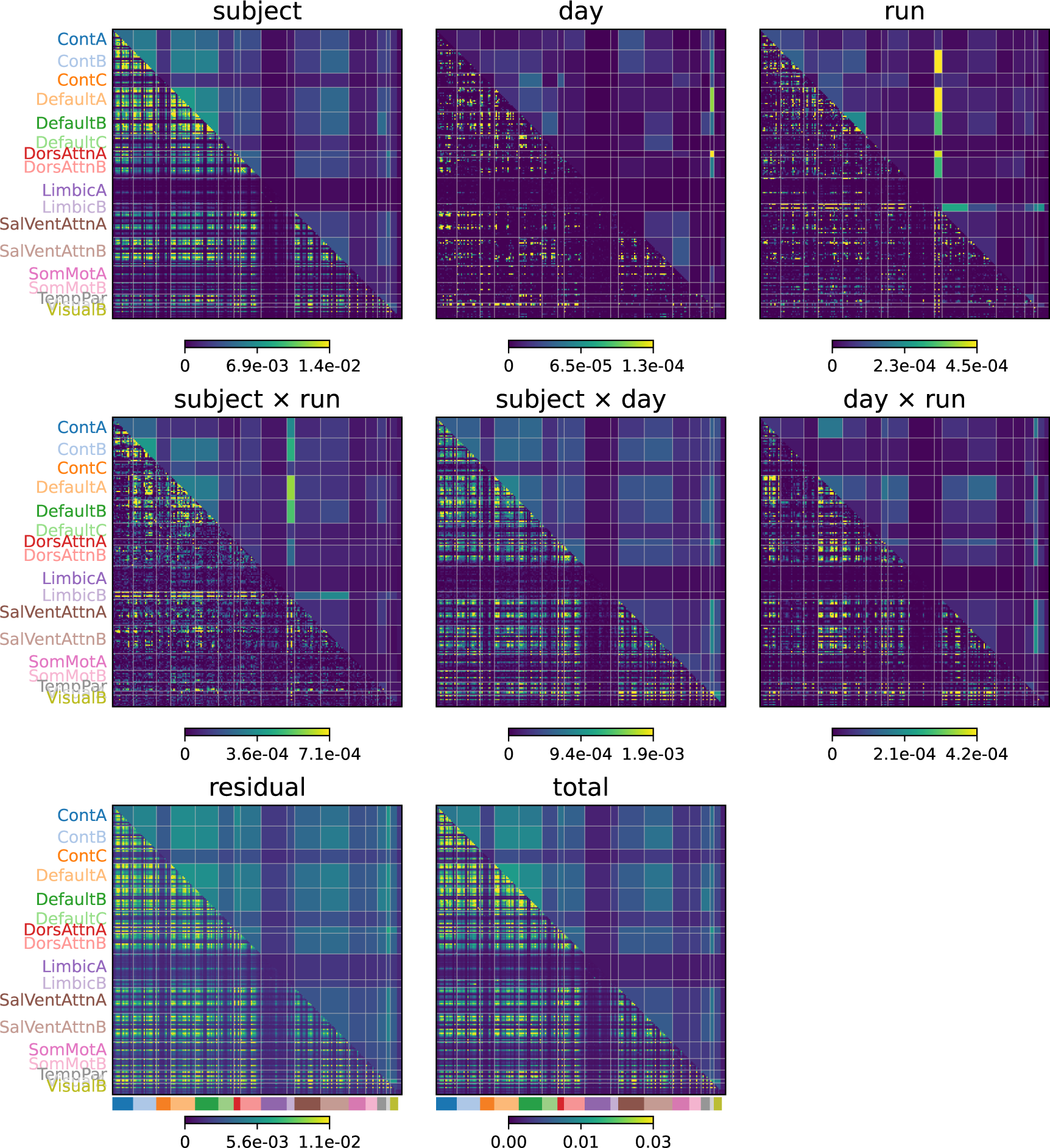
Edge connectivity on the HCP dataset parcellated with Yeo’s parcellation by source of variation. Same as in Figure S4 but not averaged by rows.

**Figure S6:**
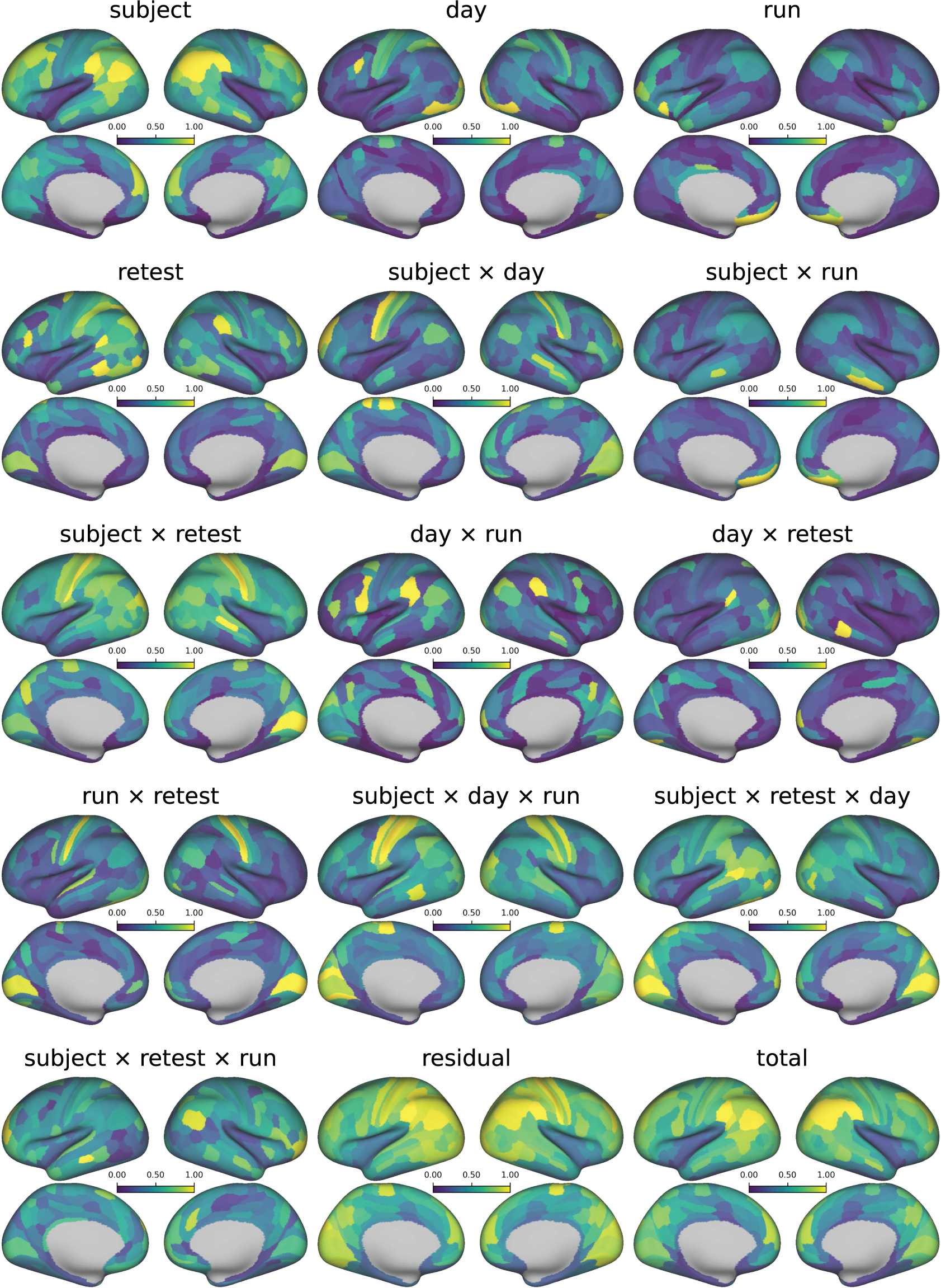
Variability of connectivity on the HCP Retest dataset parcellated with Glasser’s multimodal parcellation by source of variation.

**Figure S7:**
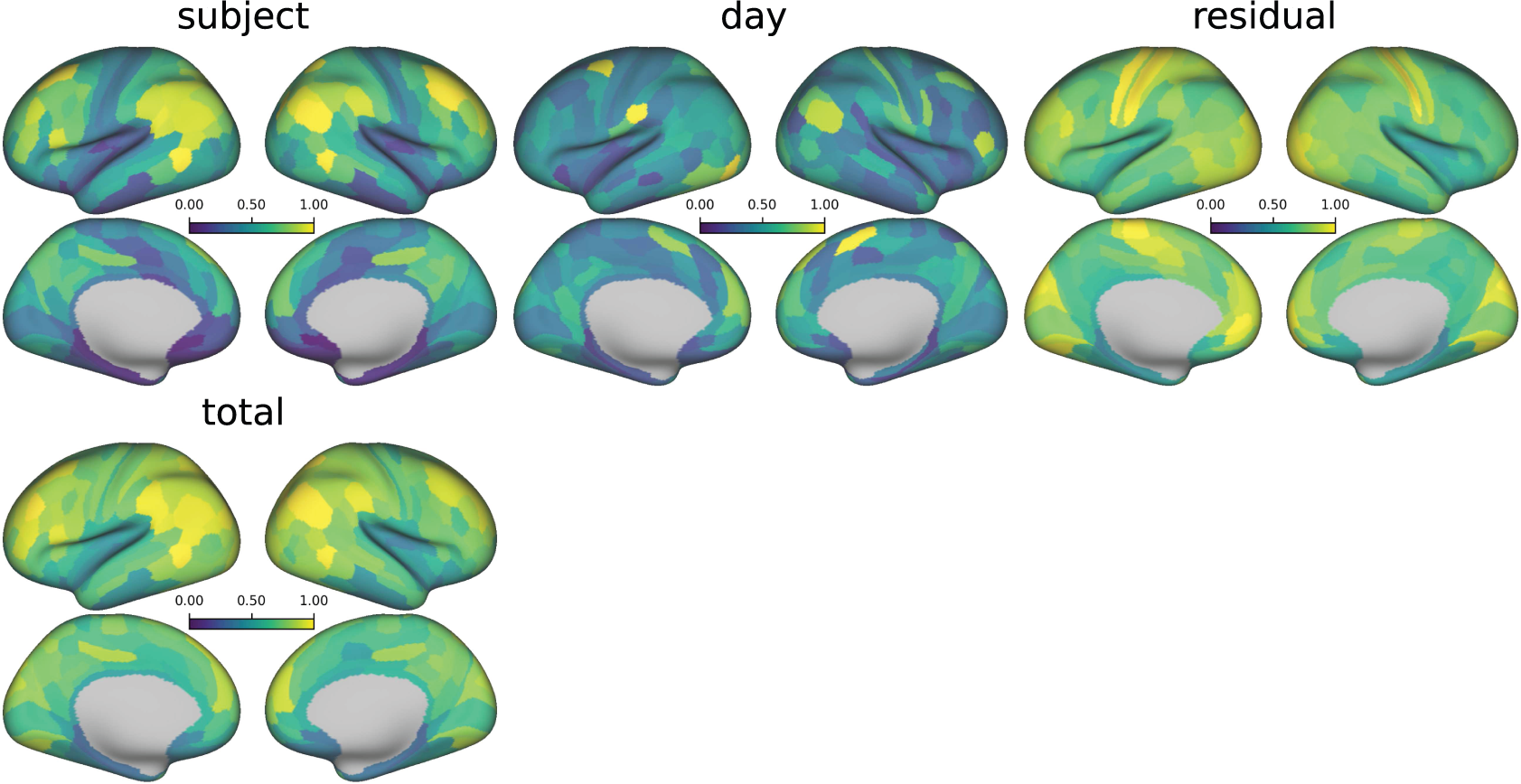
Variability of connectivity on the MMI Ljubljana dataset (eyes open condition) parcellated with Glasser’s multimodal parcellation by source of variation.

**Figure S8:**
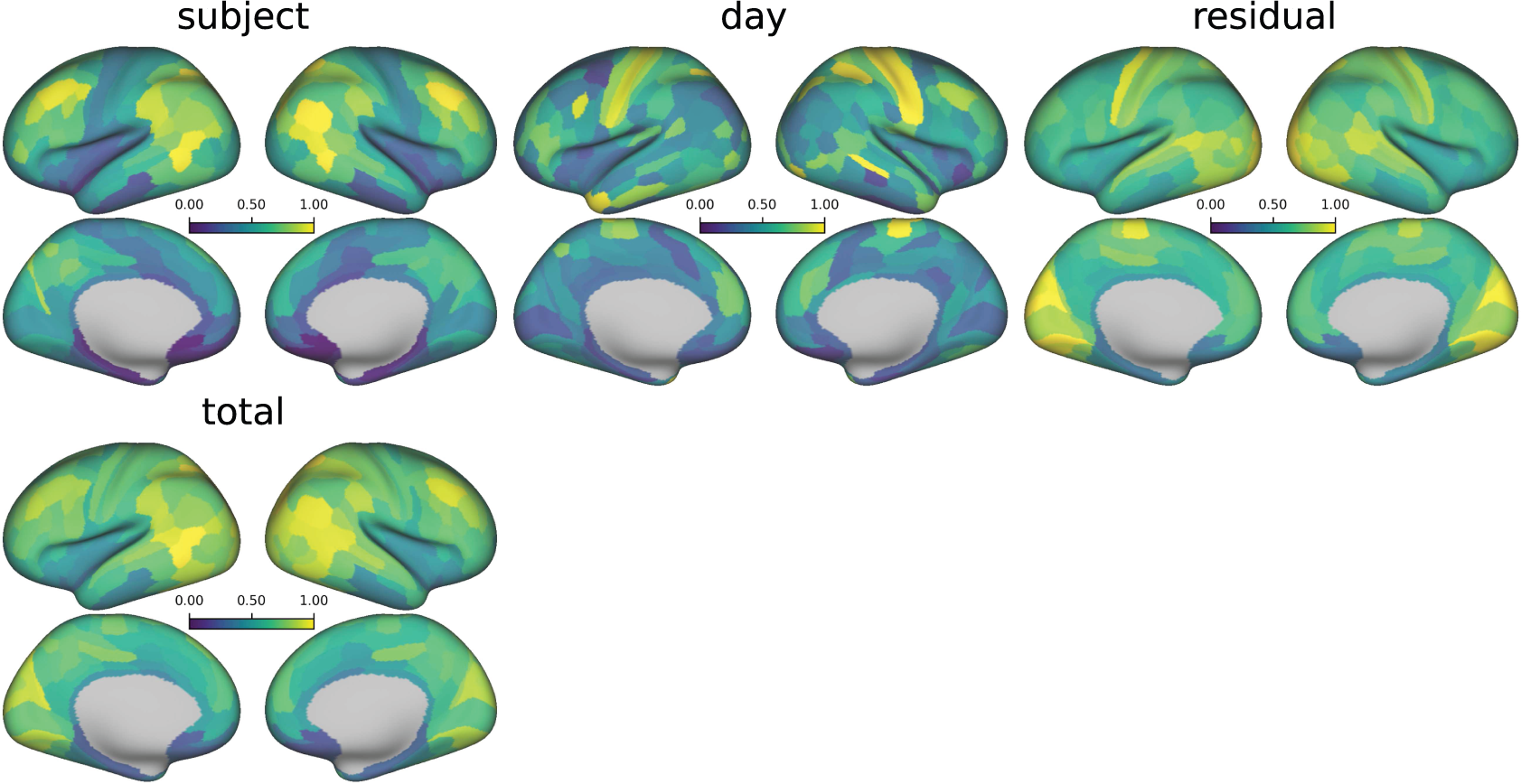
Variability of connectivity on the MMI Ljubljana dataset (eyes closed condition) parcellated with Glasser’s multimodal parcellation by source of variation.

**Figure S9:**
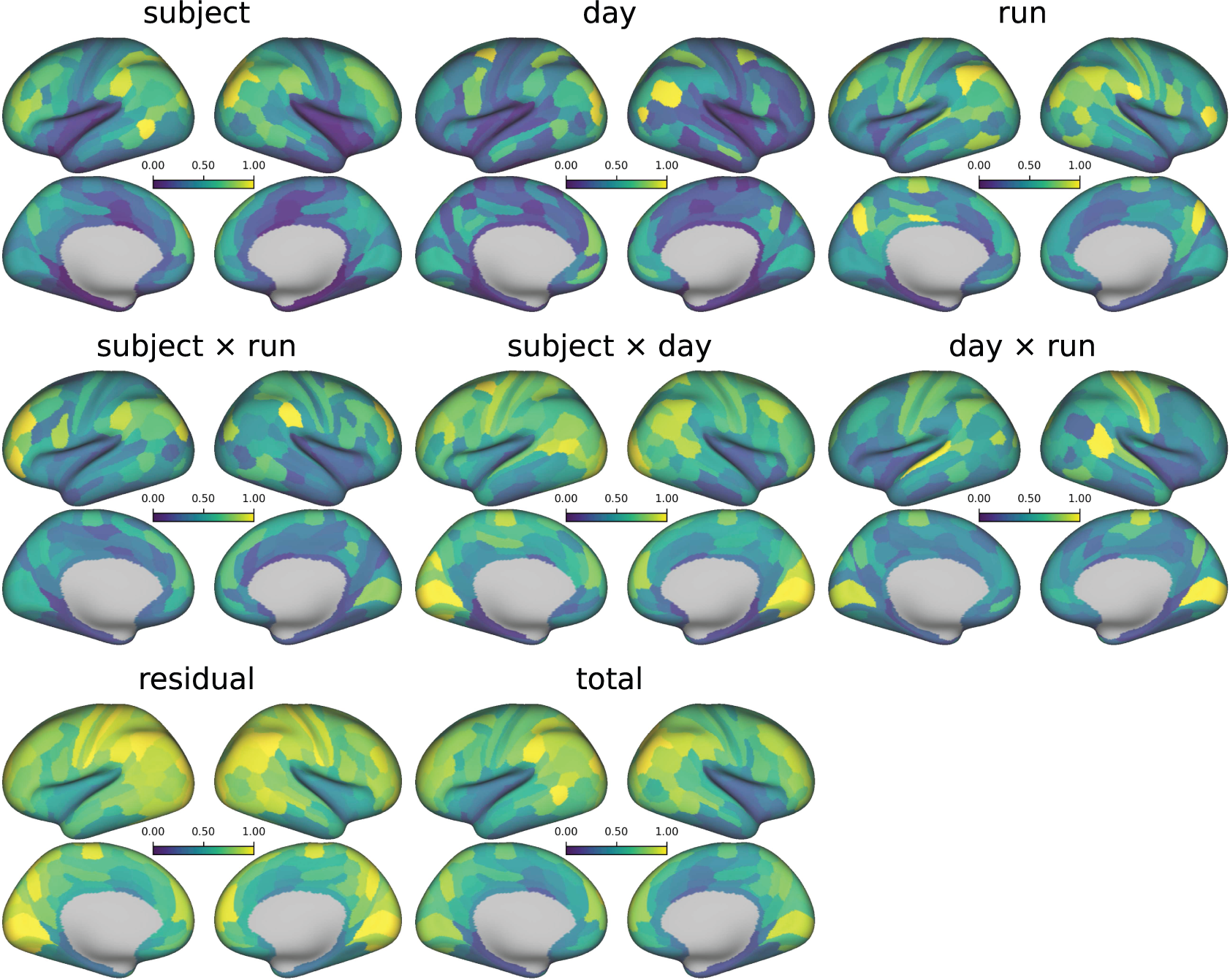
Variability of connectivity on the YaleTRT dataset parcellated with Glasser’s multimodal parcellation by source of variation.

### 10.2. Similarity of FC variability patterns across datasets

**Figure S10:**
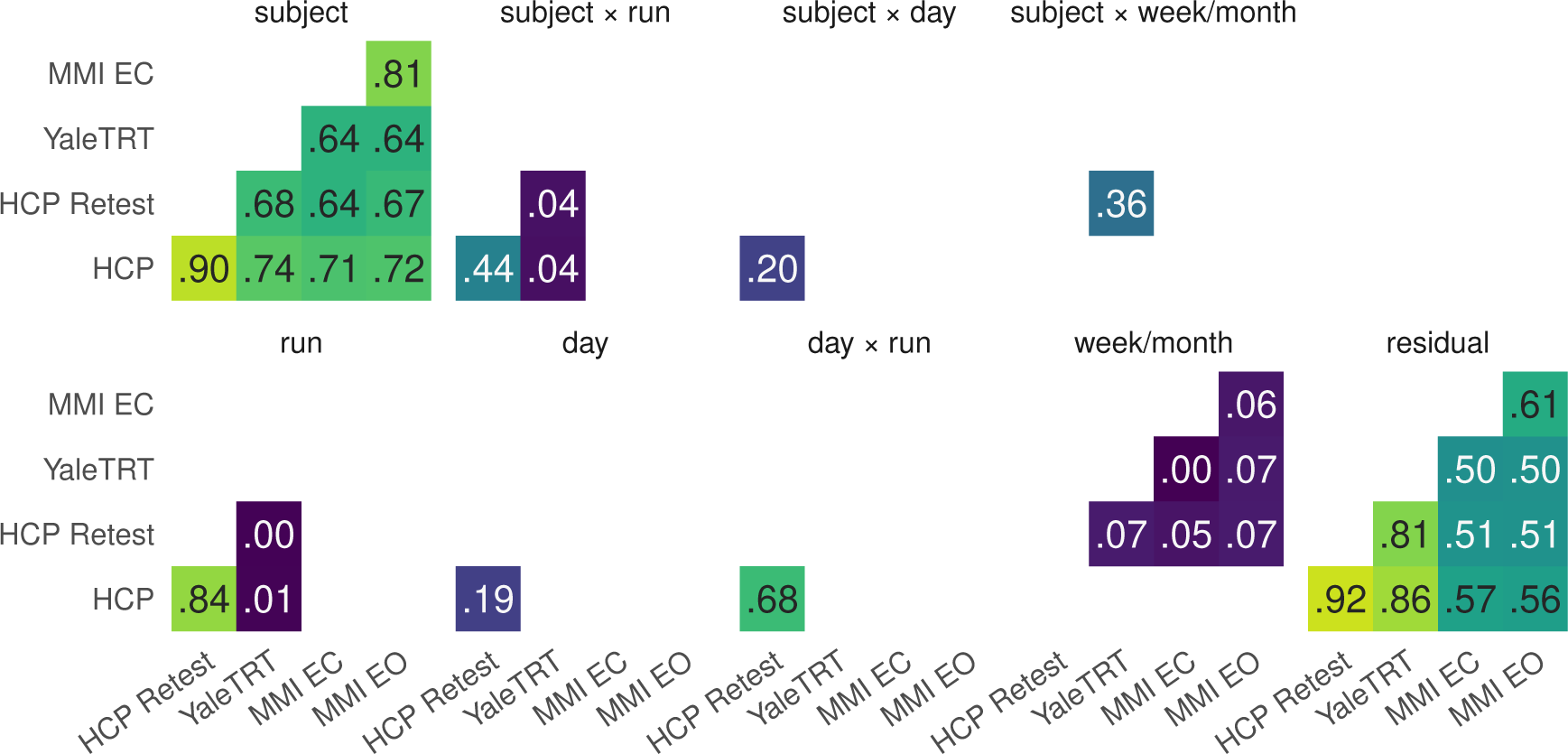
Correlations between edge variability patterns across datasets for each source of variability on Schaefer’s Local Global parcellation.

**Figure S11:**
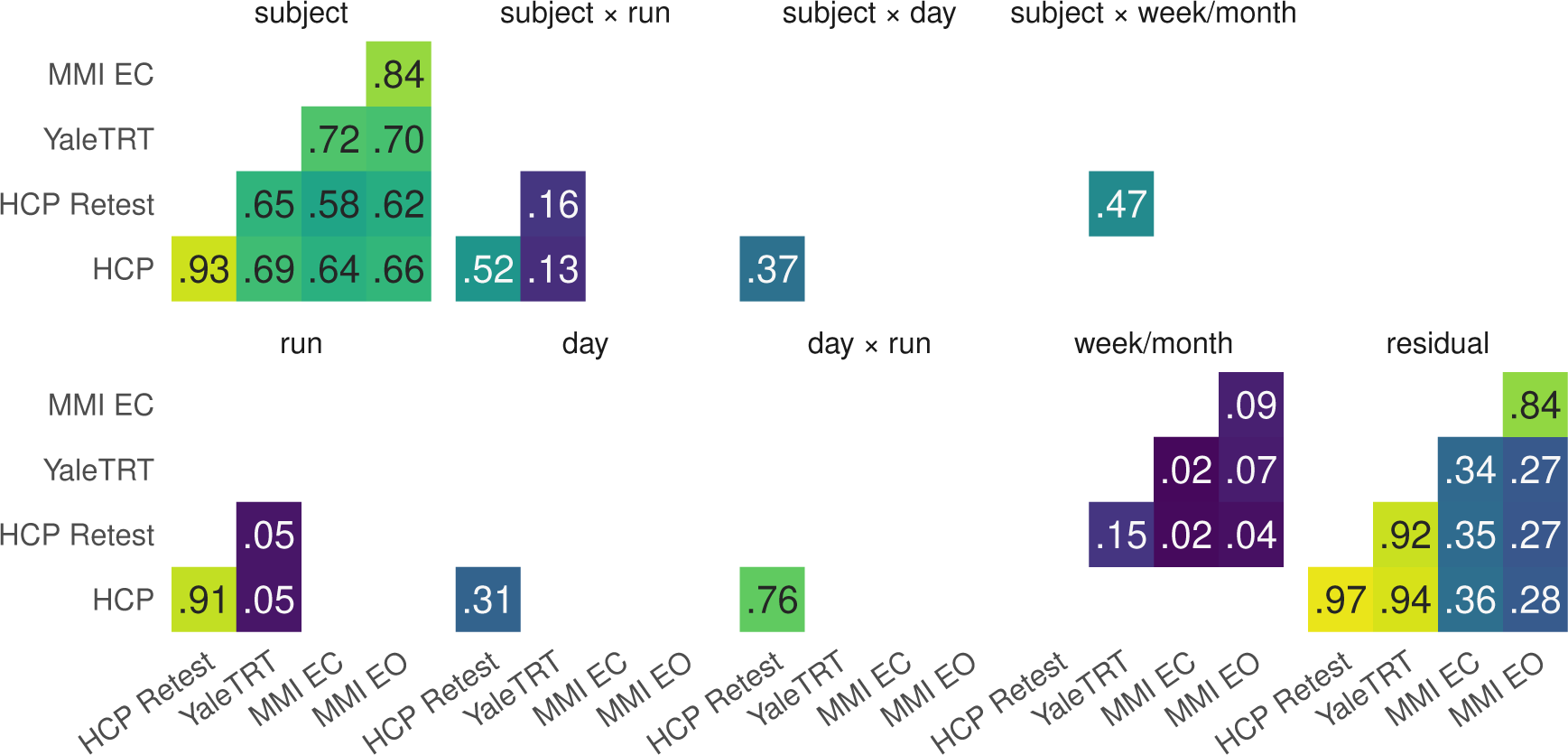
Correlations between edge variability patterns across datasets for each source of variability source on Yeo’s 17 network parcellation.

### 10.3. Similarity of FC variability patterns across sources of variability

**Figure S12:**
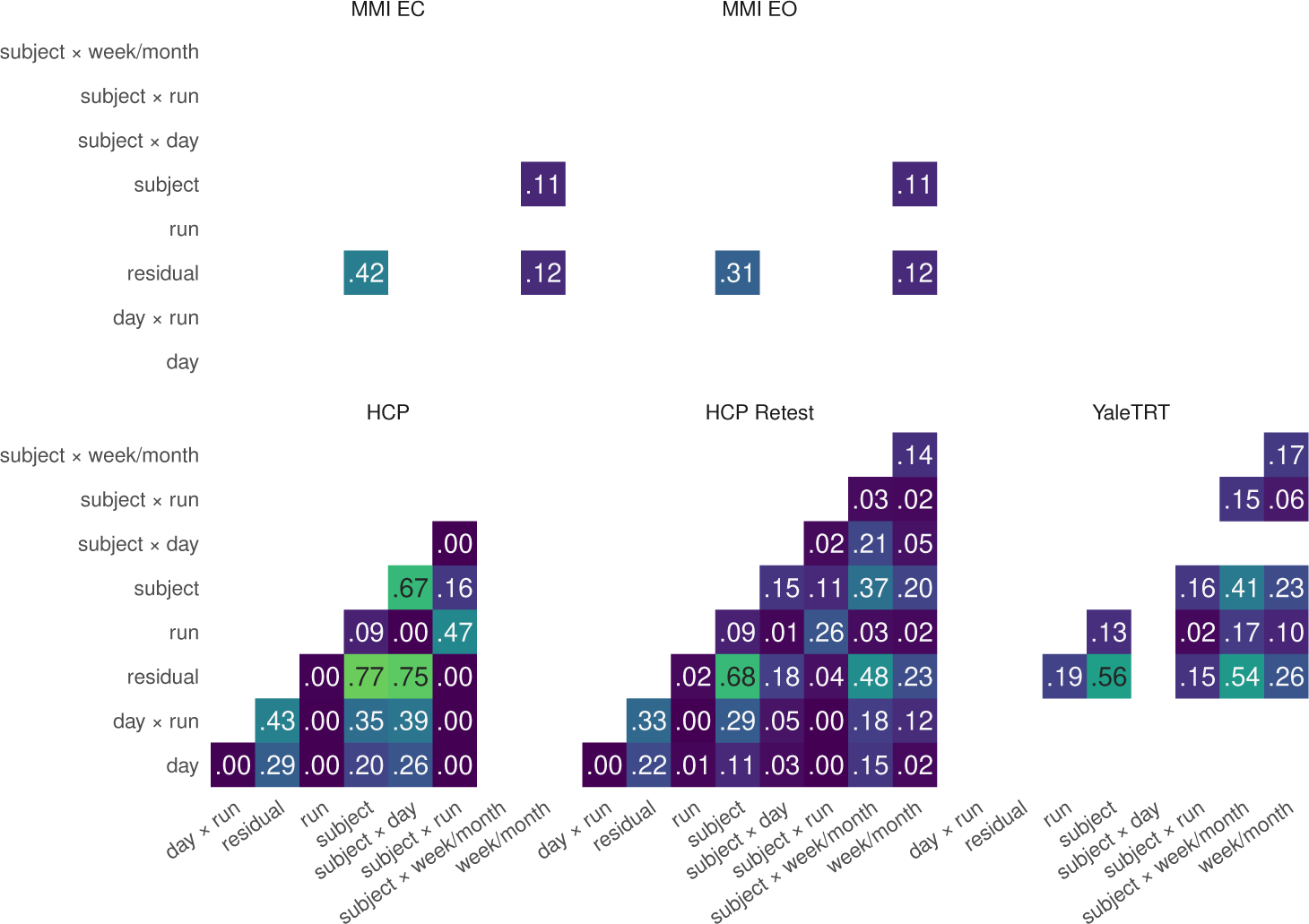
Correlations between edge variability patterns across sources of variability for each dataset on Glasser’s multimodal parcellation.

**Figure S13:**
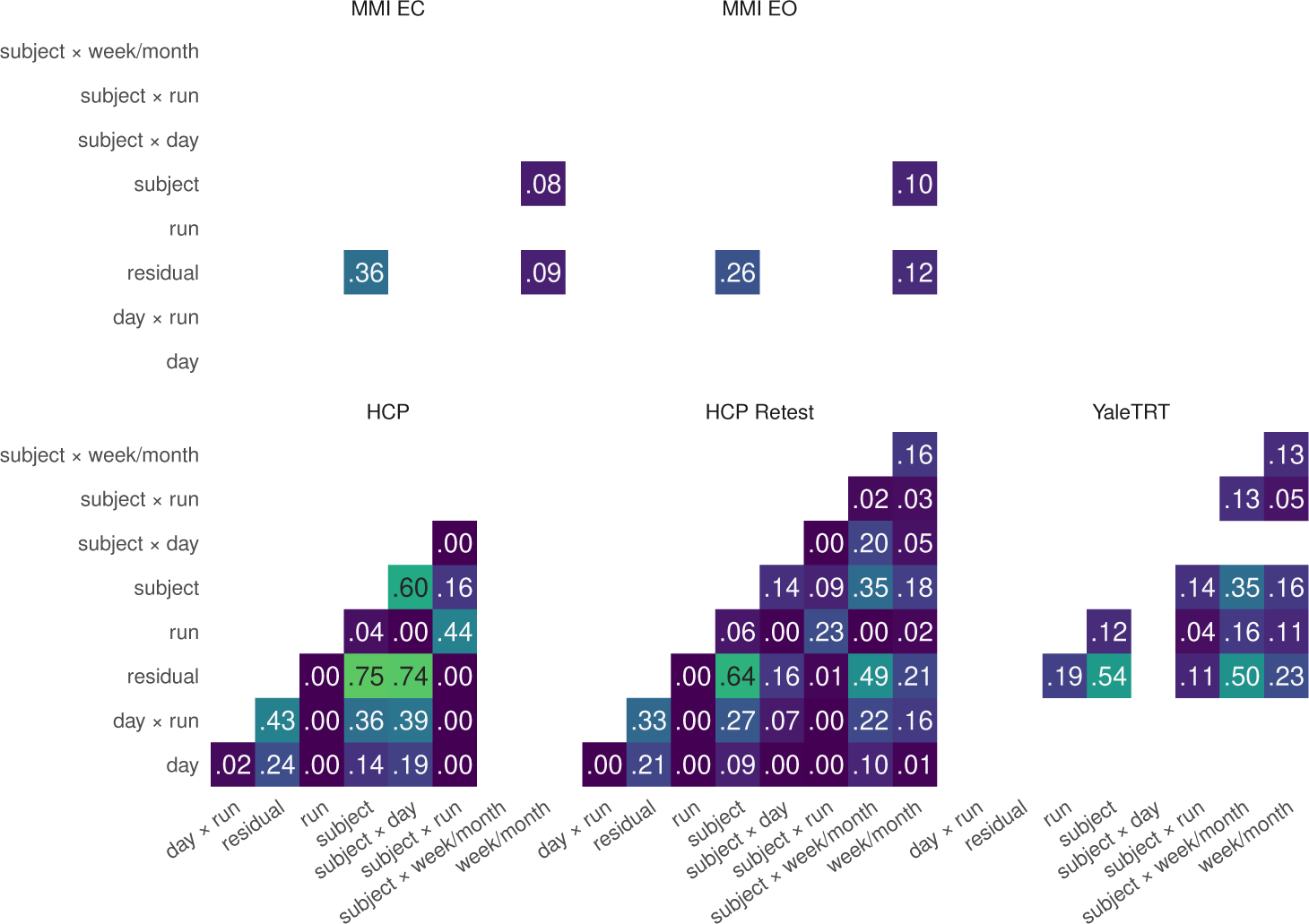
Correlations between edge variability patterns across sources of variability for each dataset on Schaefer’s Local Global parcellation.

**Figure S14:**
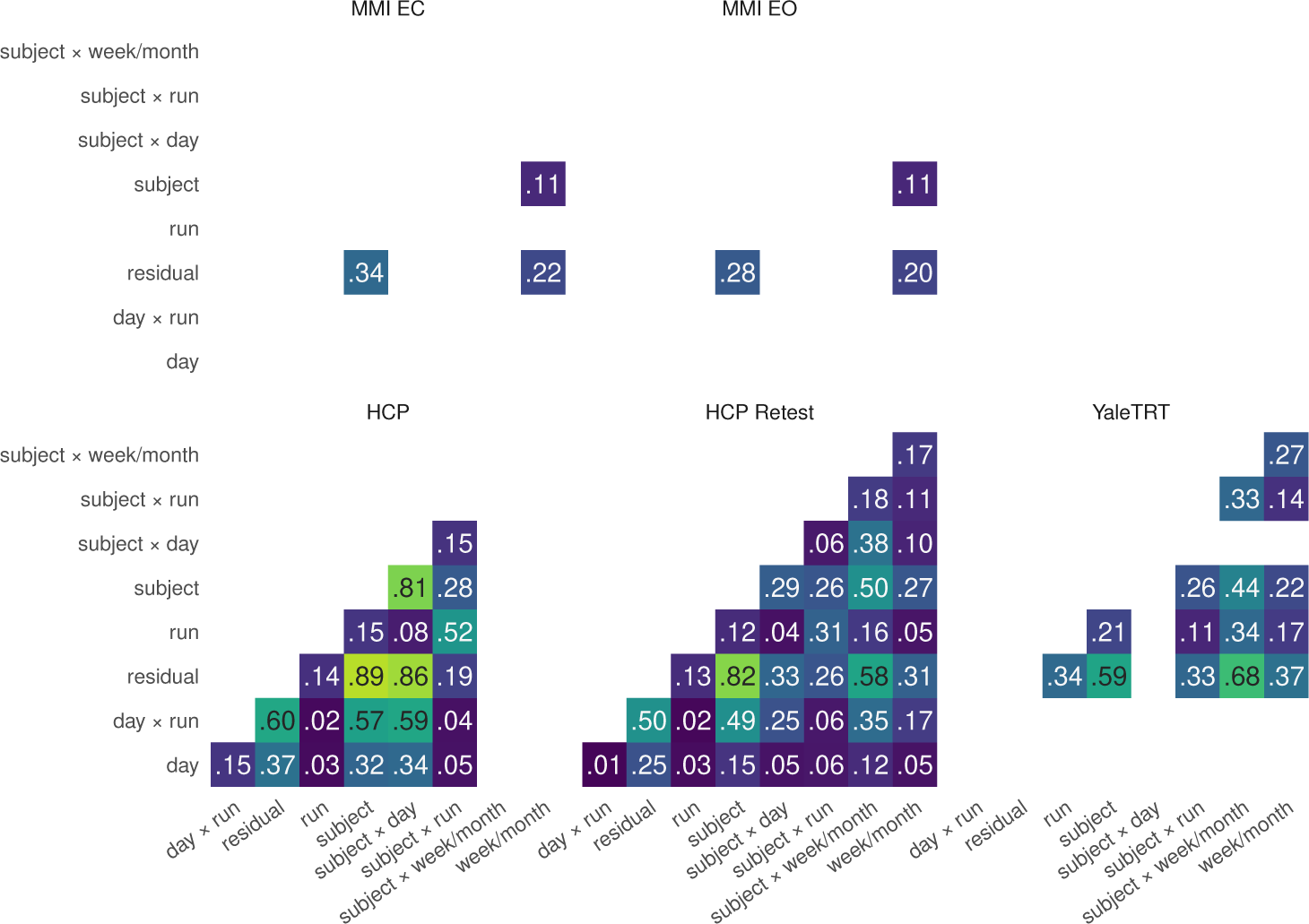
Correlations between edge variability patterns across sources of variability for each dataset on Yeo’s 17-networks parcellation.

**Figure S15:**
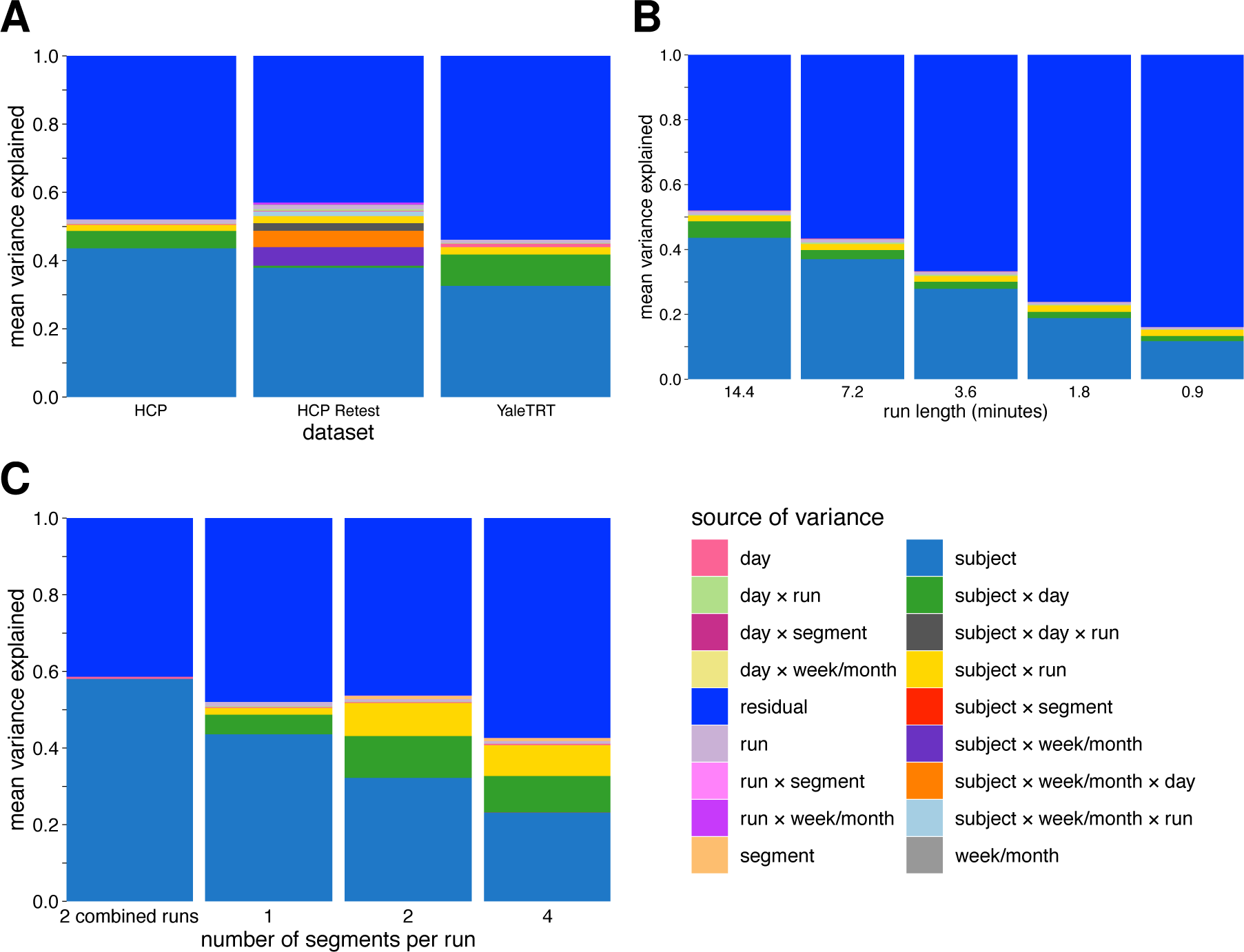
Mean edge variance as a function of dataset, run length and number of segments per run. **A.** Mean edge variance explained by each source of variance for each dataset. **B.** Mean edge variance explained as a function of run length. **C.** Mean edge variance as a function of segments per run.

### 10.4. Grouping of edges according to between-subject and within-subject variability

**Figure S16:**
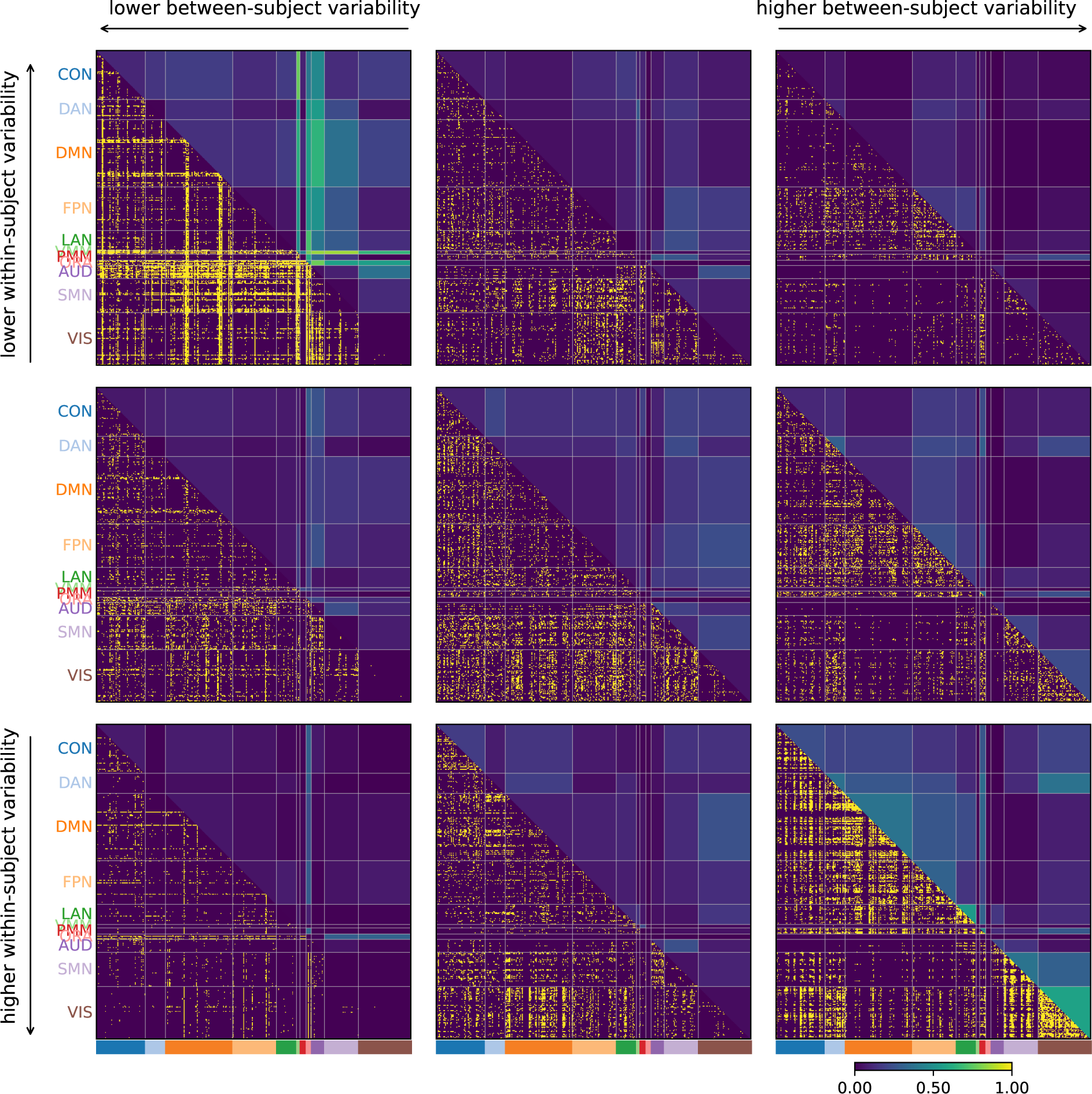
Variability of connectivity by edge as a function of within-subject and between-subject variability in the HCP dataset parcellated with Schaefer’s Local Global parcellation.

**Figure S17:**
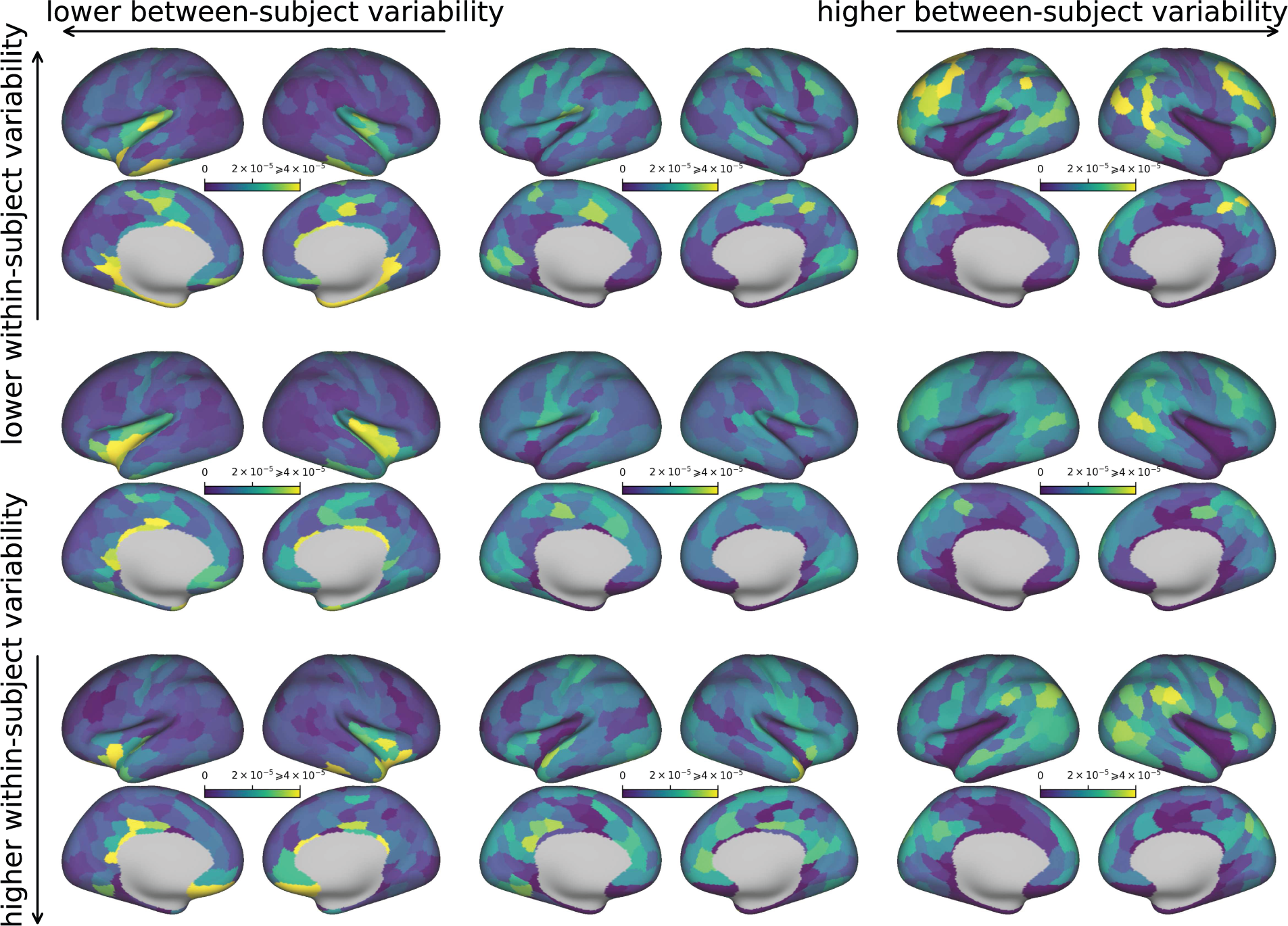
Variability of connectivity by region as a function of within-subject and between-subject variability in the HCP dataset parcellated with Schaefer’s Local Global parcellation. Same as in Figure S18 but averaged across rows of matrices.

**Figure S18:**
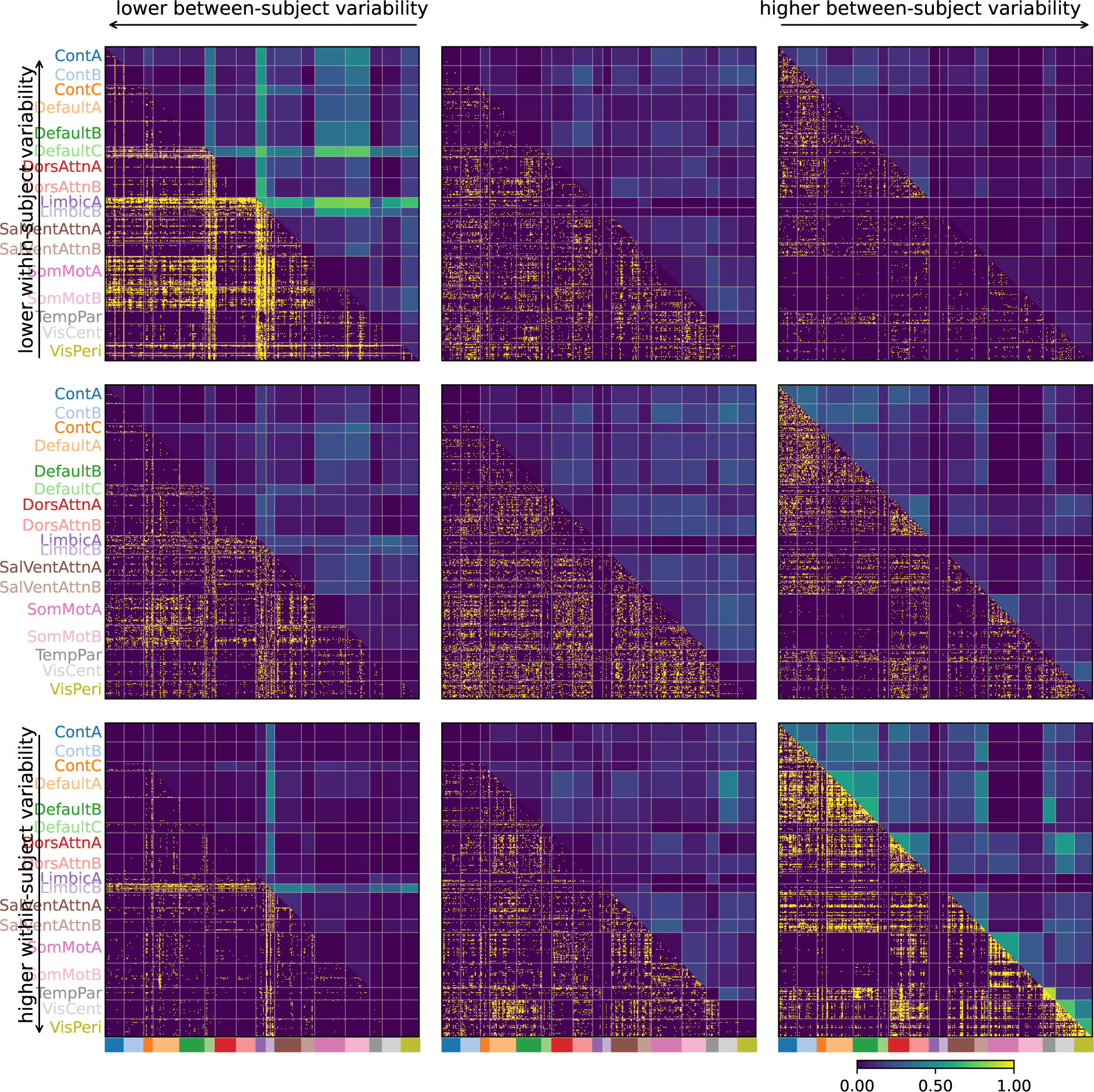
Variability of connectivity by edge as a function of within-subject and between-subject variability in HCP dataset parcellated with Schaefer’s Local Global parcellation.

**Figure S19:**
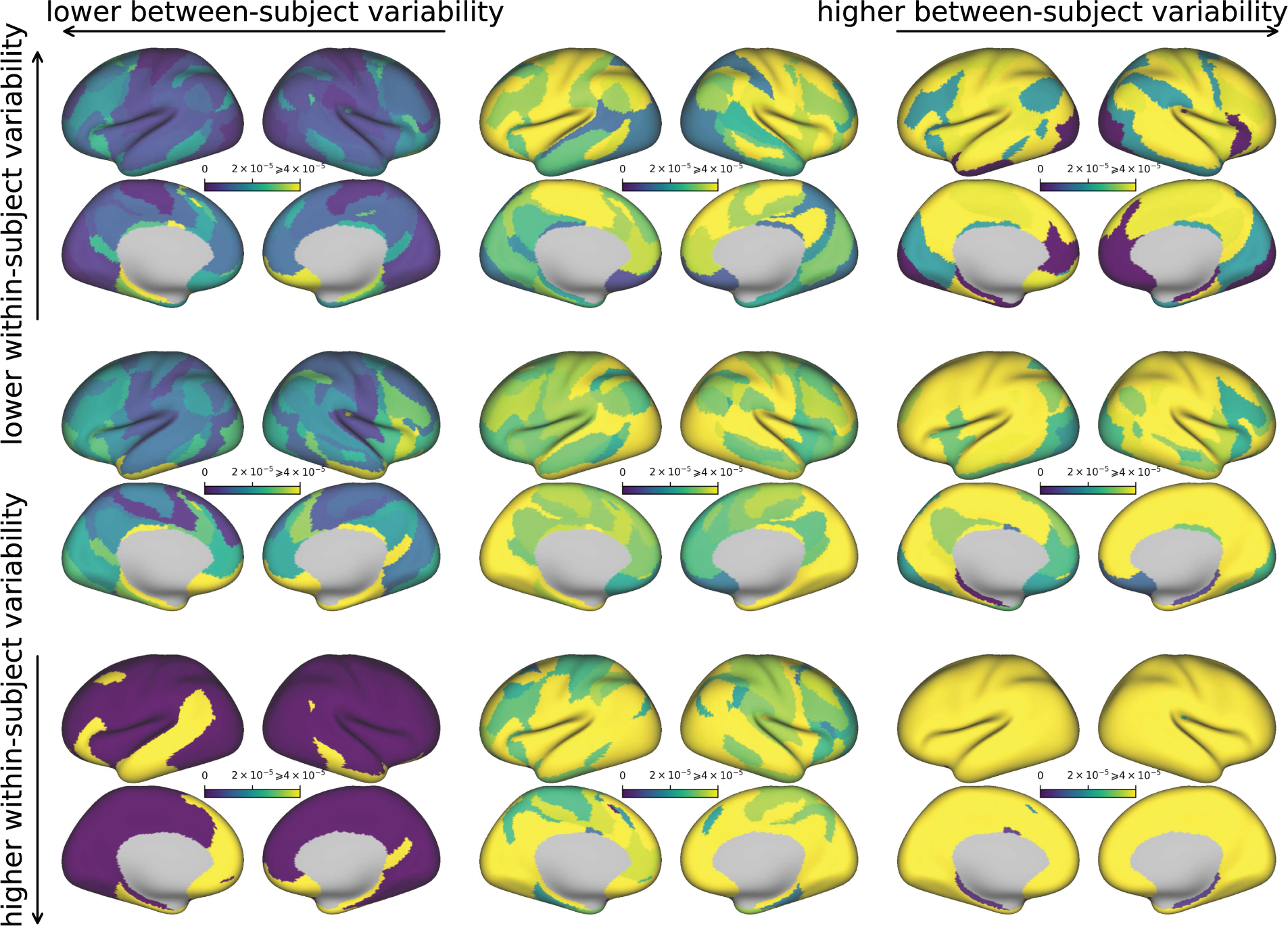
Variability of connectivity by region as a function of within-subject and between-subject variability in HCP dataset parcellated with Yeo’s 17 network parcellation.

**Figure S20:**
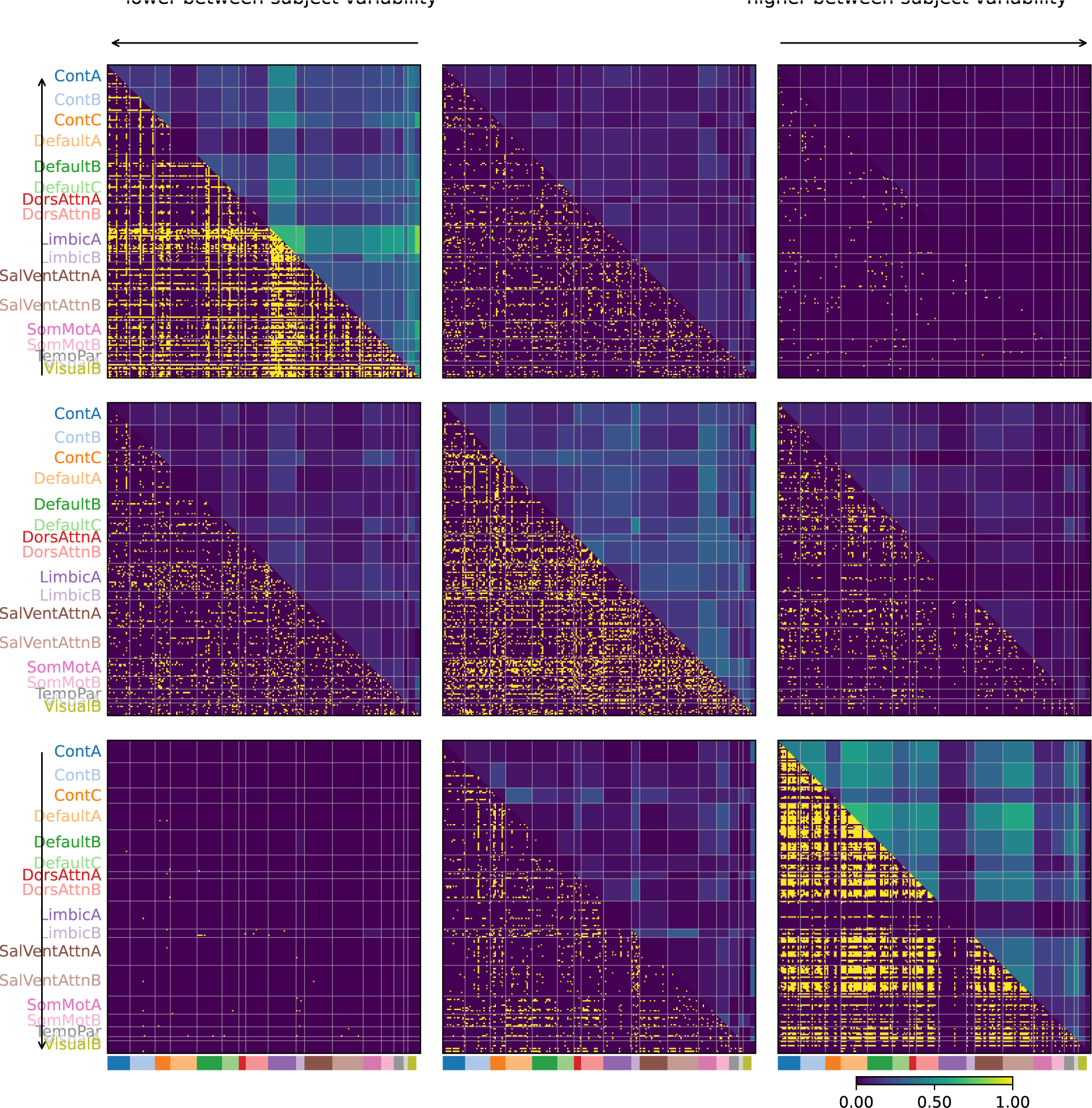
Variability of connectivity by edge as a function of within-subject and between-subject variability in HCP dataset parcellated with Schaefer’s Local Global parcellation.

### 10.5. Intraclass coefficient with respect to to between-subject and within-subject variance

**Figure S21:**
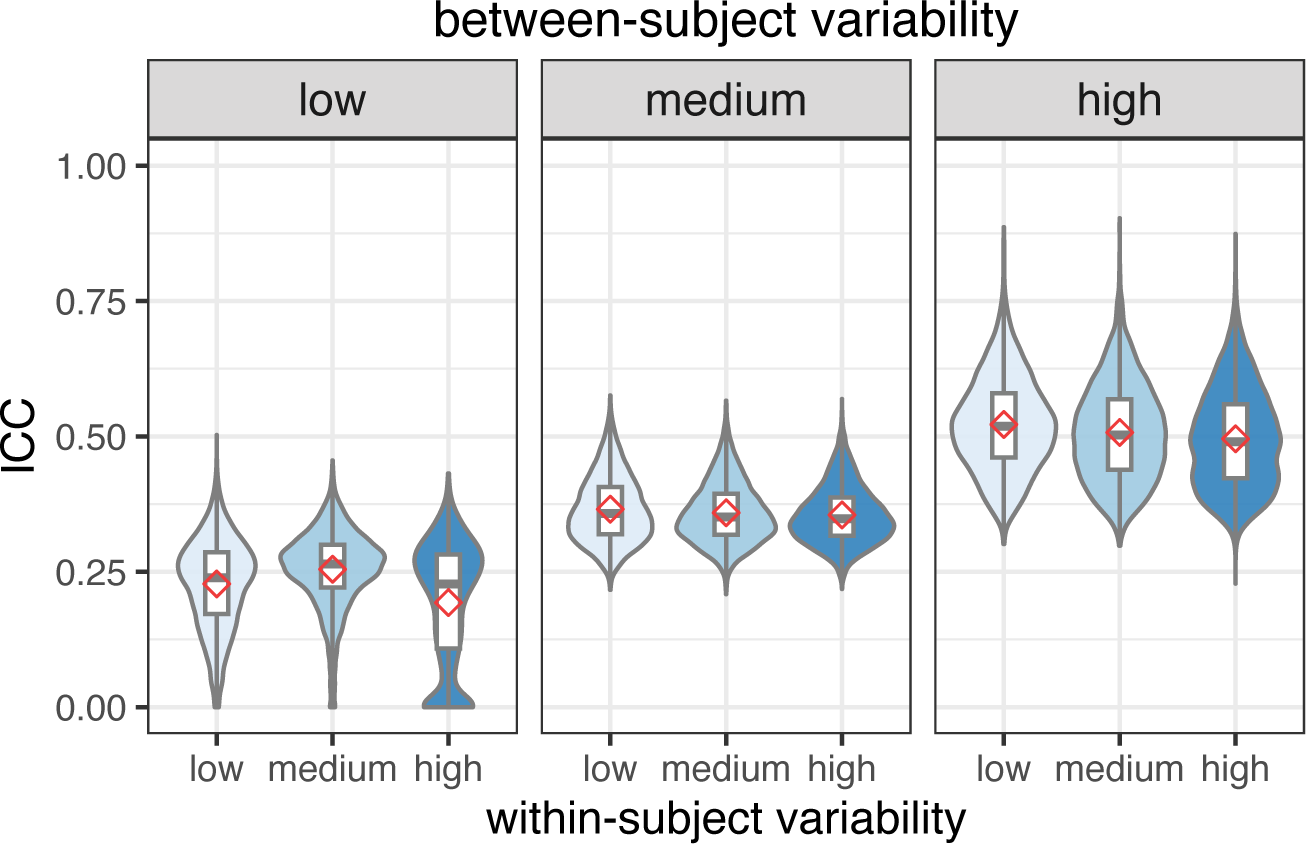
Intraclass coefficient as a function of between-subject and within-subject variability on Schaefer’s parcellation.

**Figure S22:**
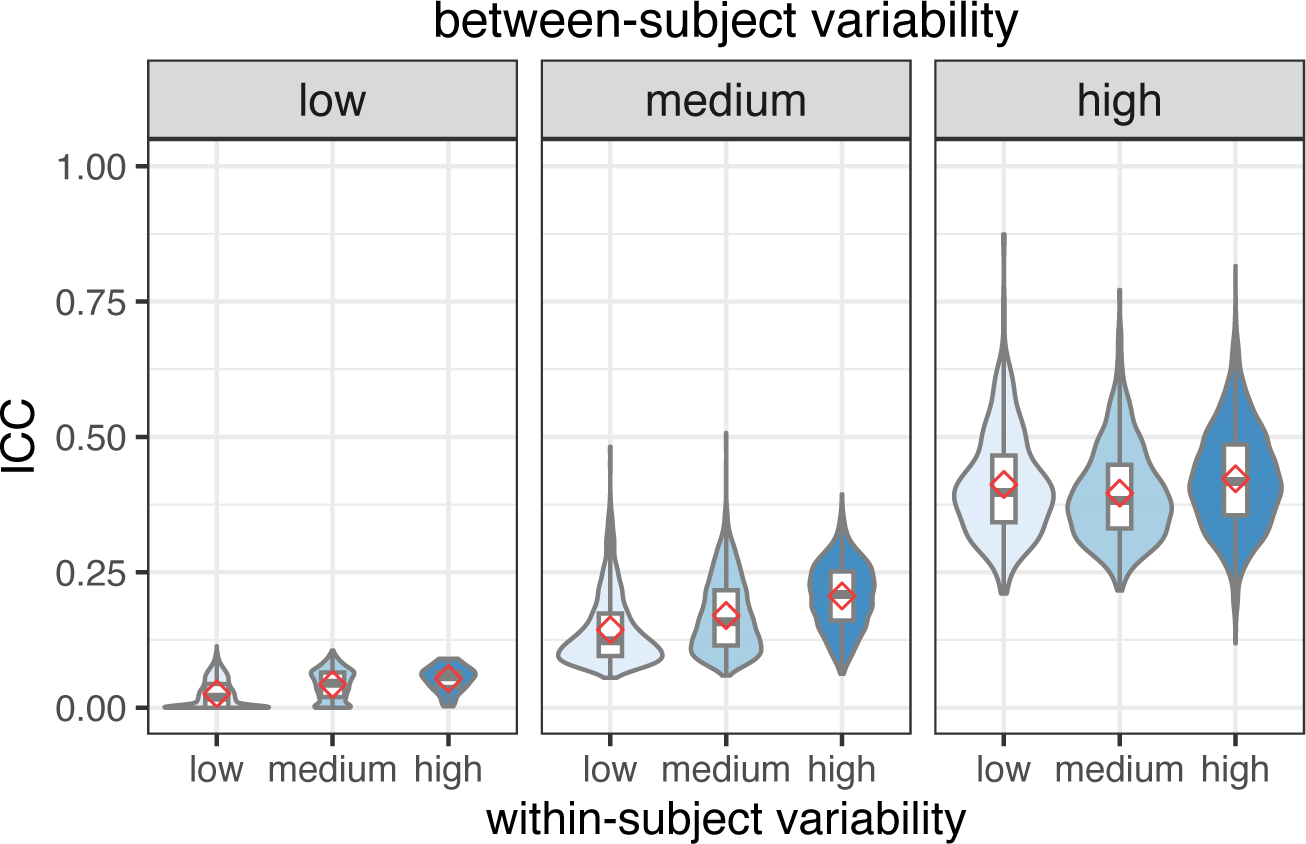
Intraclass coefficient as a function of between-subject and within-subject variability on Yeo’s parcellation.

### 10.6. External validation

**Figure S23:**
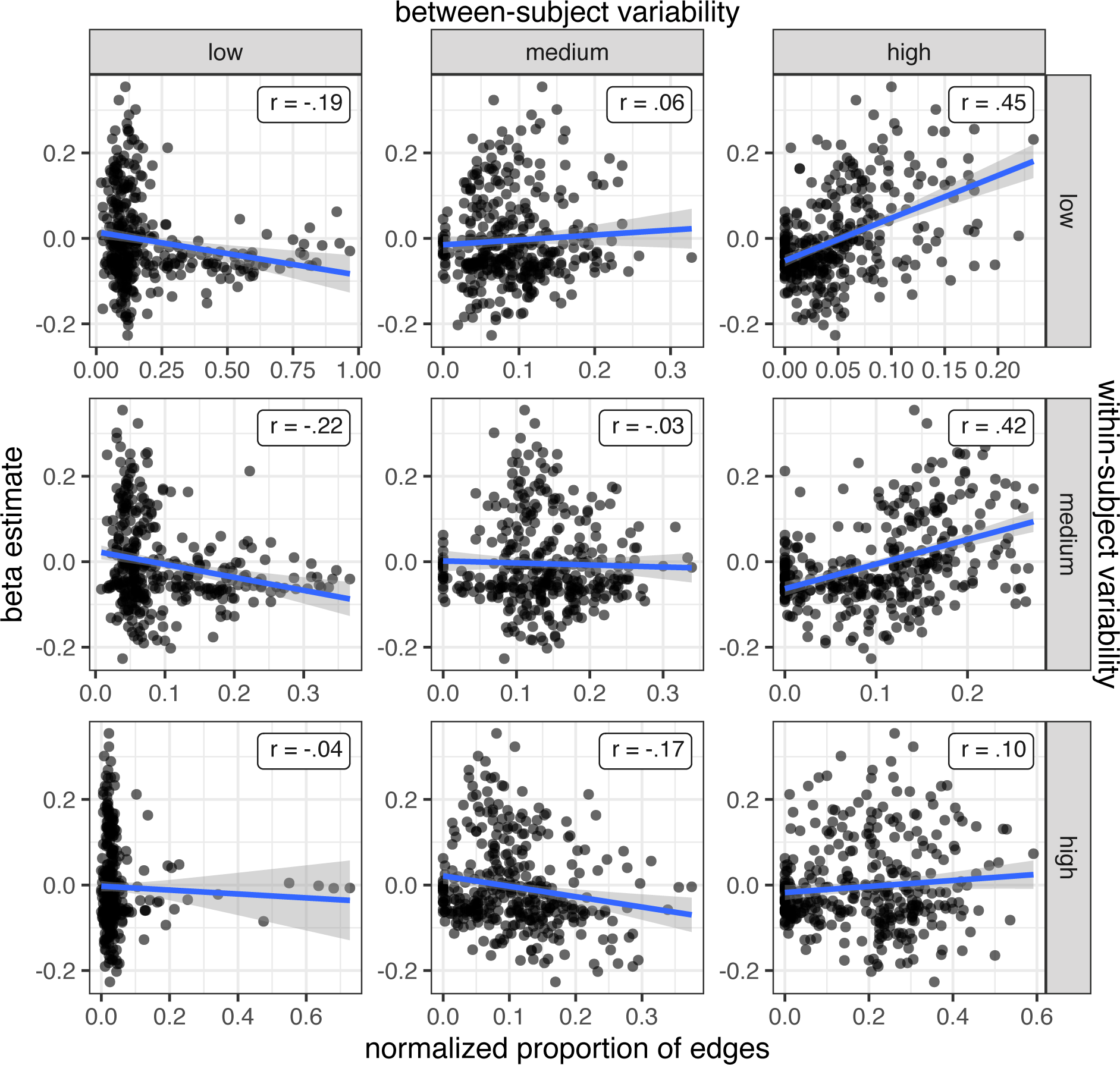
Correlations between the map of domain-general cognitive core regions and the proportion of edges in each region by groups of edges. Parcellated using Glasser’s parcellation.

**Figure S24:**
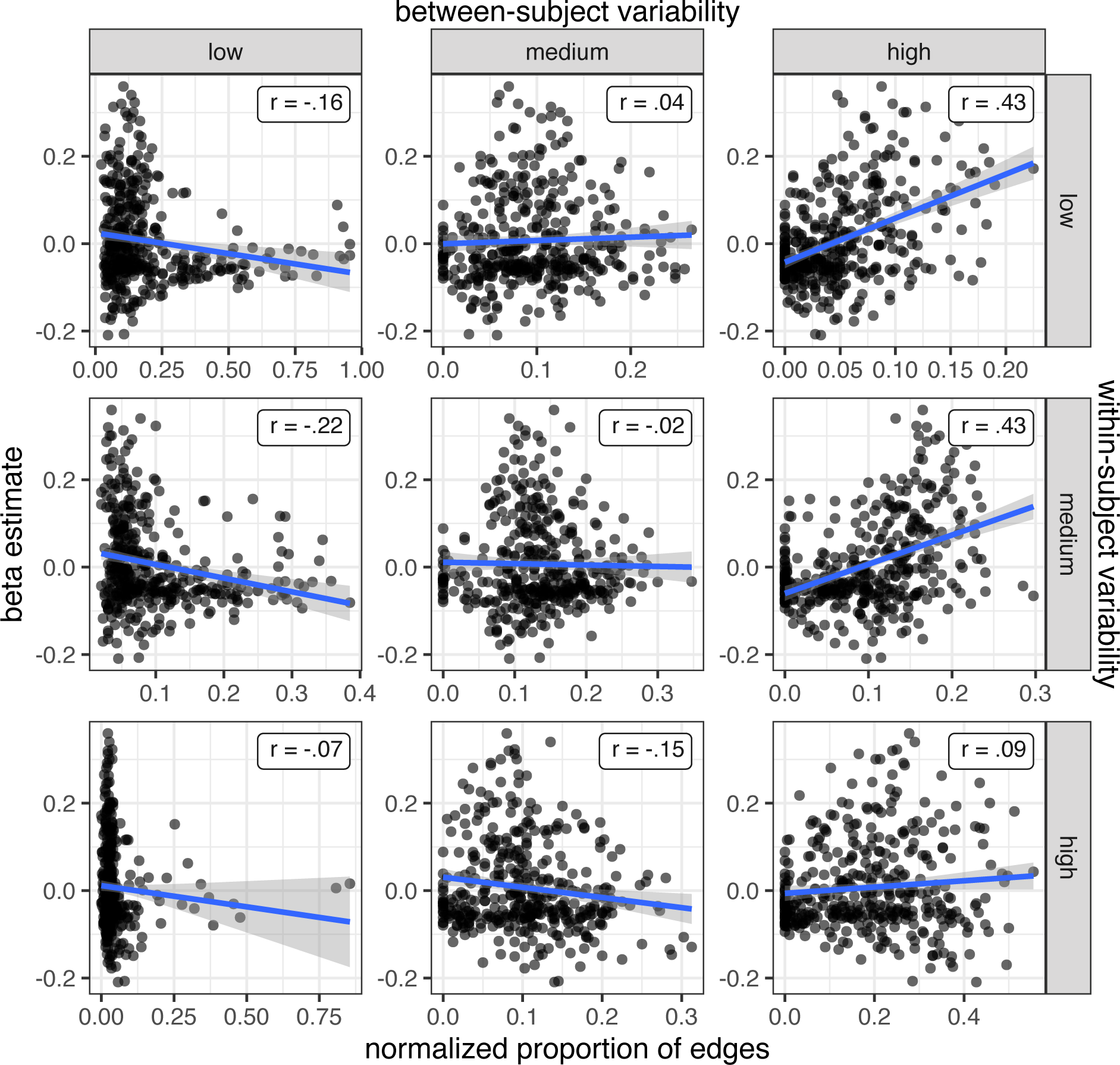
Correlations between the map of domain-general cognitive core regions and the proportion of edges in each region by groups of edges. Parcellated using Schaefer’s parcellation.

**Figure S25:**
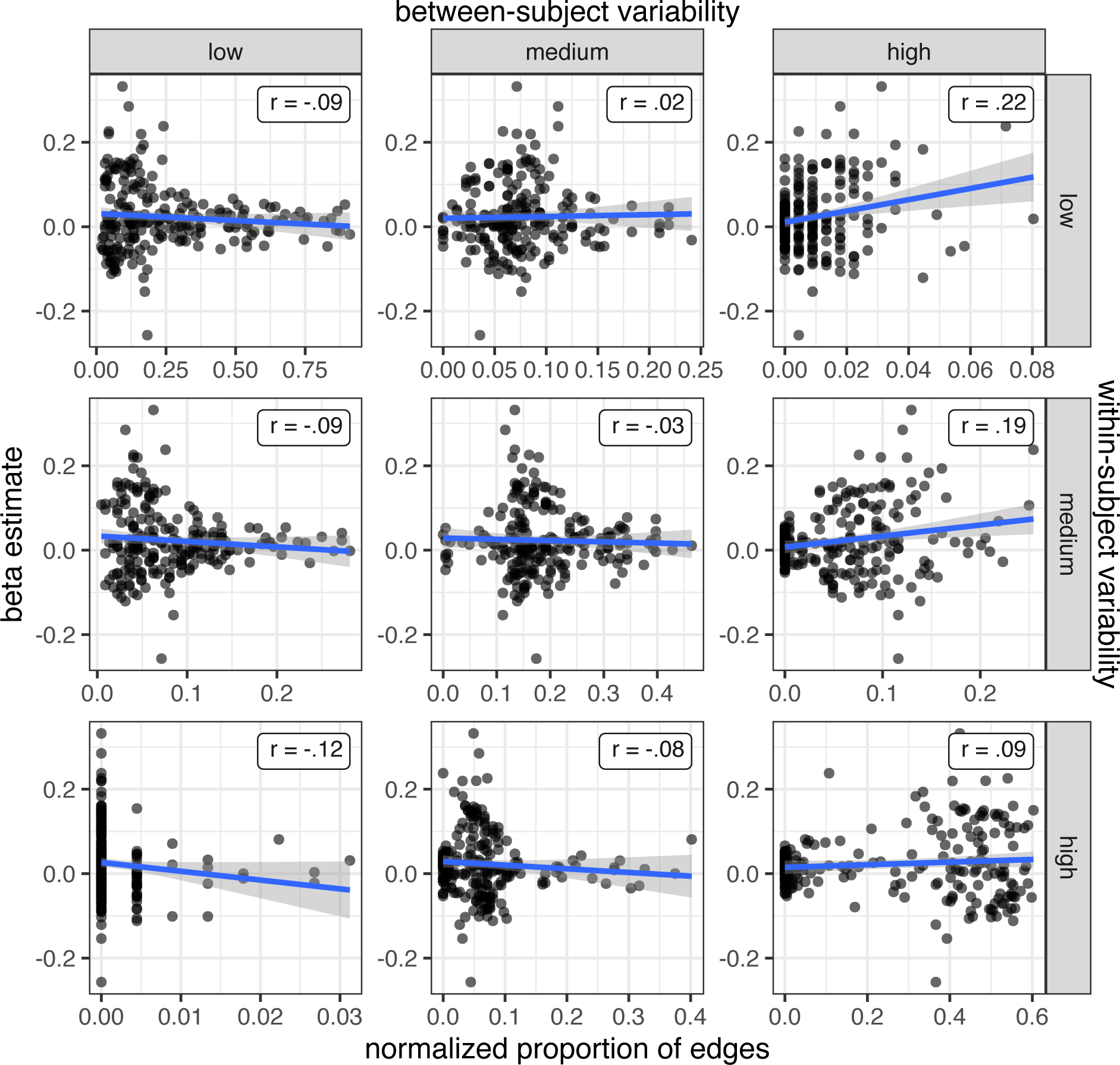
Correlations between the map of domain-general cognitive core regions and the proportion of edges in each region by groups of edges. Parcellated using Yeo’s parcellation.

### 10.7. Brain-behavior correlations

**Figure S26:**
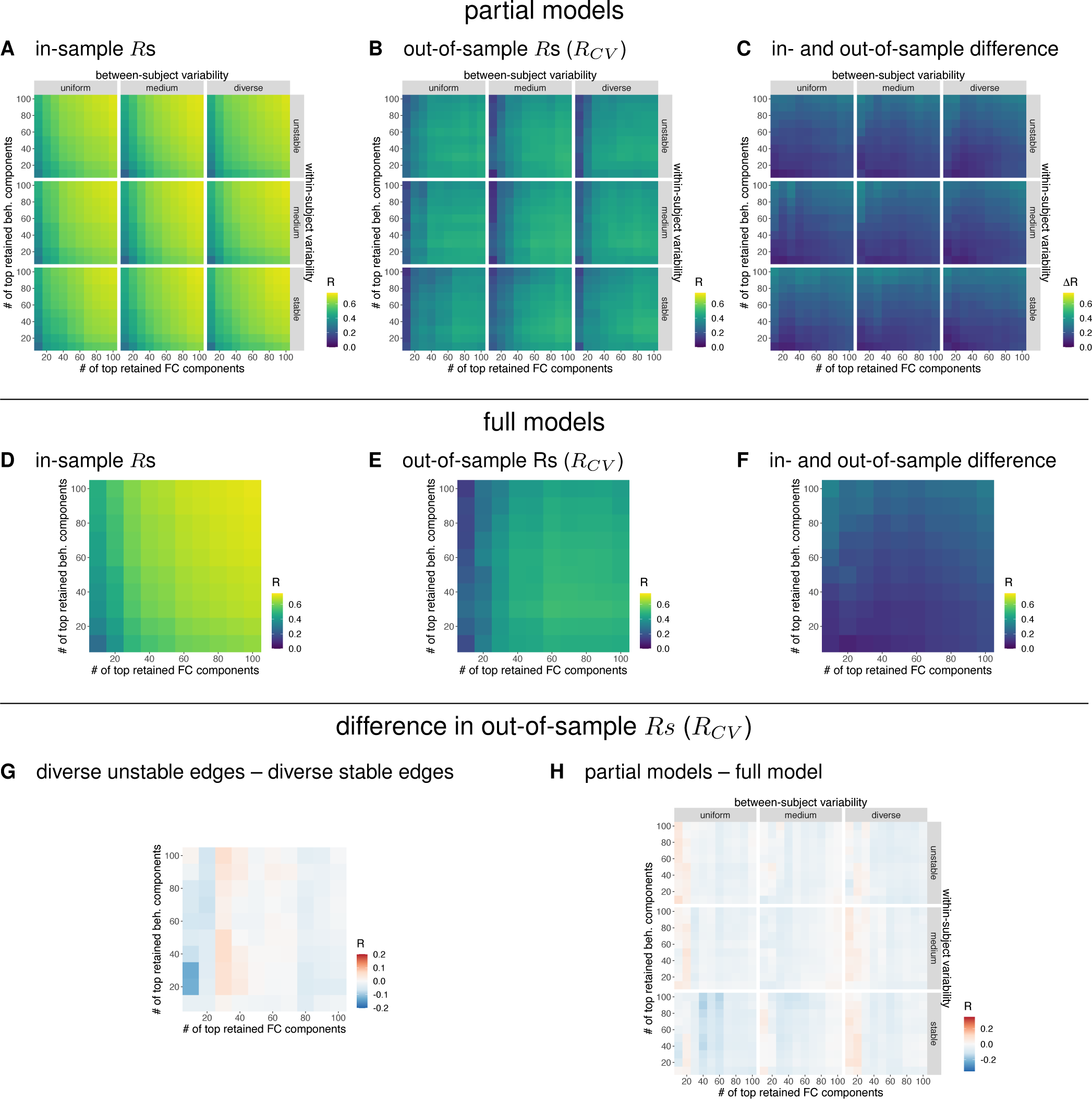
Results of CCA on Schaefer’s parcellation. **A.** In-sample canonical correlations for the partial models based on groups of edges according to their between-subject and within-subject variability. **B.** Out-of-sample canonical correlations for the partial models. **C.** Difference between in-sample and out-of-sample canonical correlations for the partial models. **D.** In-sample canonical correlations for the full models. **E.** Out-of-sample canonical correlations for the full models. **F.** Difference in out-of-sample canonical correlations between full and partial models. Red color indicates higher canonical correlations for partial models. **G.** Difference between in-sample and out-of-sample canonical correlations for full models. **H.** Difference in out-of-sample canonical correlations between models based on diverse unstable edges and models based on diverse stable edges. Blue color indicates higher canonical correlations for models based on diverse stable edges.

**Figure S27:**
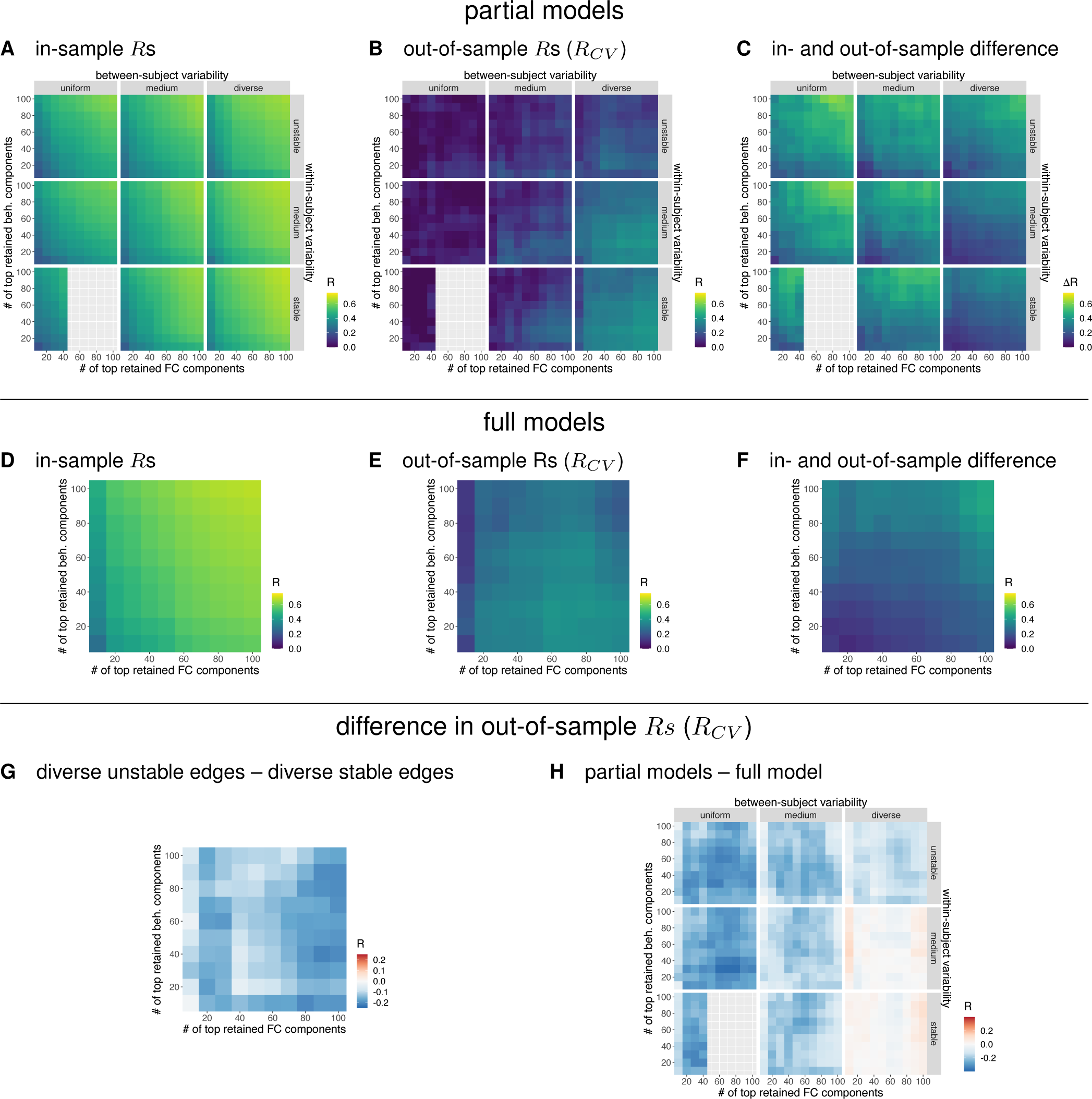
Results of CCA on Yeo’s parcellation. **A.** In-sample canonical correlations for the partial models based on groups of edges according to their between-subject and within-subject variability. **B.** Out-of-sample canonical correlations for the partial models. **C.** Difference between in-sample and out-of-sample canonical correlations for the partial models. **D.** In-sample canonical correlations for the full models. **E.** Out-of-sample canonical correlations for the full models. **F.** Difference in out-of-sample canonical correlations between full and partial models. Red color indicates higher canonical correlations for partial models. **G.** Difference between in-sample and out-of-sample canonical correlations for full models. **H.** Difference in out-of-sample canonical correlations between models based on diverse unstable edges and models based on diverse stable edges. Blue color indicates higher canonical correlations for models based on diverse stable edges.

**Figure S28:**
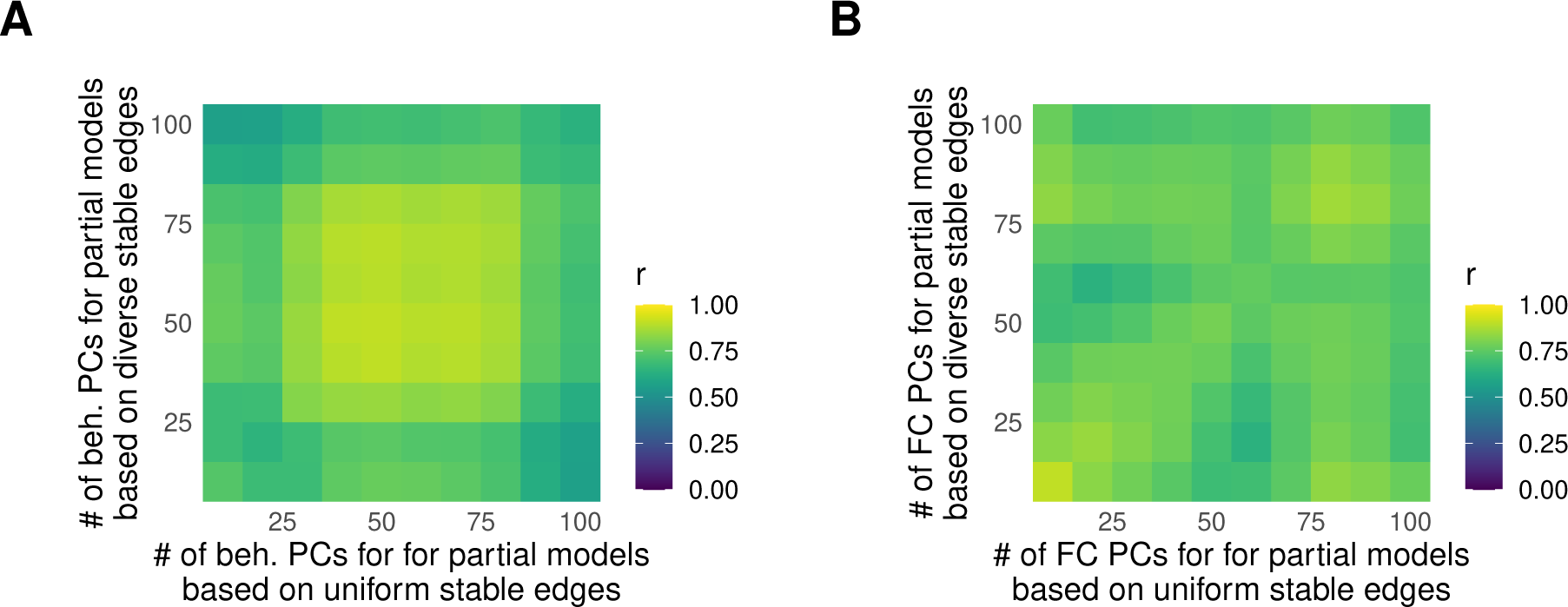
Similarity of CCA weights between partial models based on diverse and uniform stable edges as a function of the number of principal components (PCs) included in the model. **A.** Similarity as a function of behavioral PCs. **B.** Similarity as a function of functional connectivity PCs. Results refer to Glasser’s parcellation.

**Figure S29:**
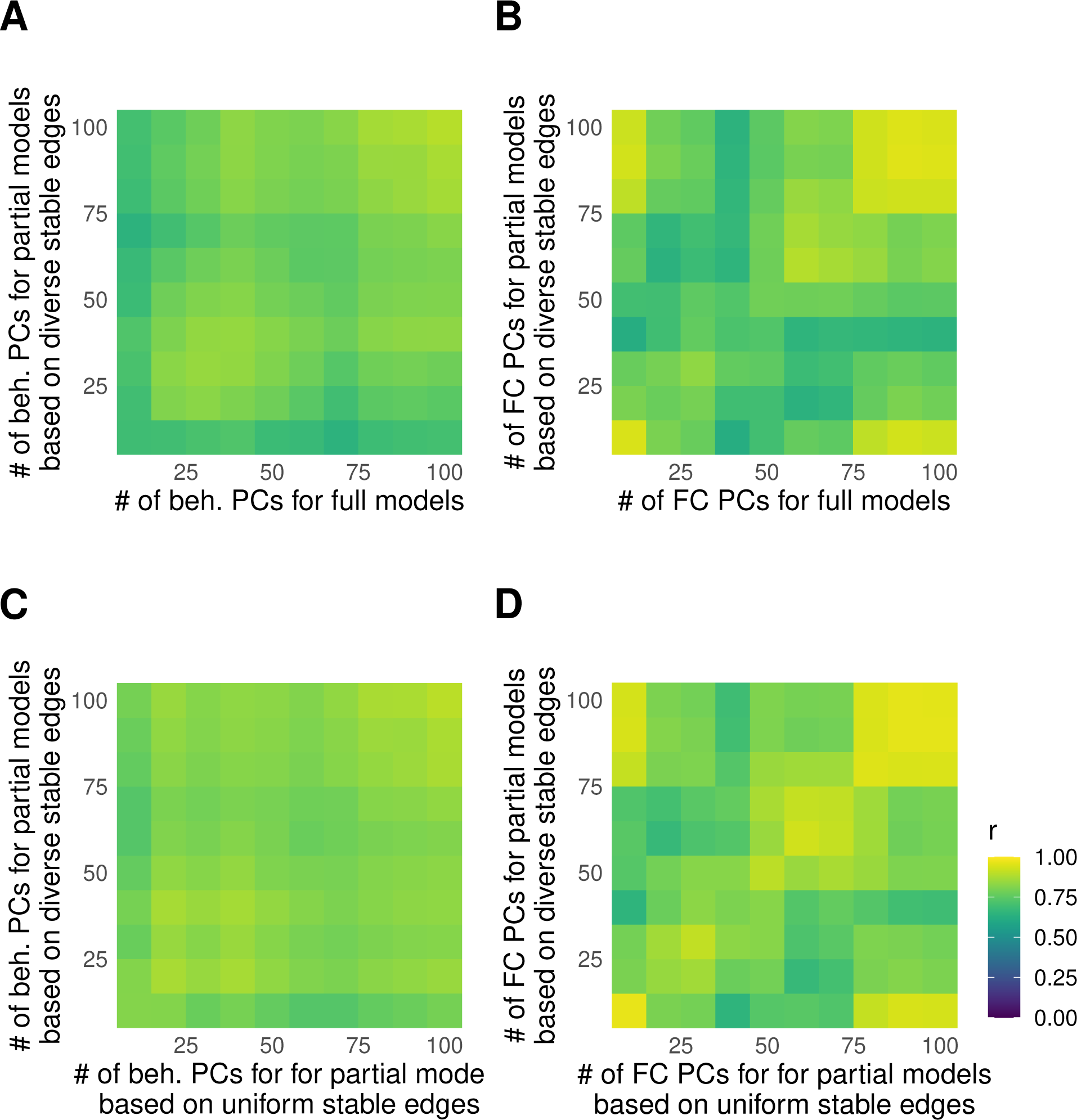
Similarity of weights between CCA models as a function of the number of principal components (PCs) included in the model. Results refer to **Schaefer’s parcellation. A.** Similarity between models based on diverse stable edges and the full model as a function of behavioral PCs. **B.** Similarity between models based on diverse stable edges and full model as a function of functional connectivity PCs. **C.** Similarity between models based on diverse and uniform stable edges as a function of behavioral PCs. **D.** Similarity between models based on diverse and uniform stable edges as a function of functional connectivity PCs.

**Figure S30:**
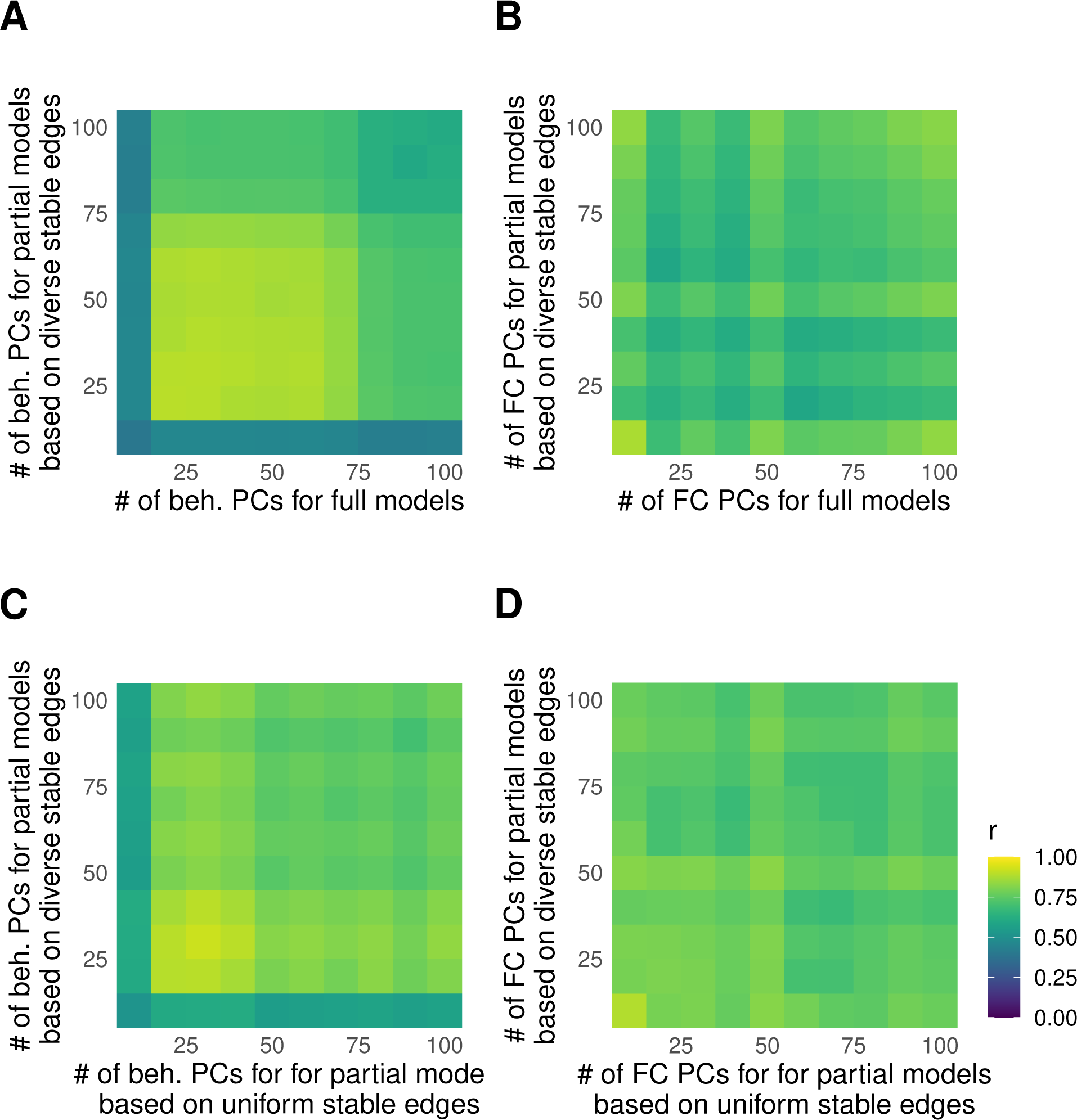
Similarity of weights between CCA models as a function of the number of principal components (PCs) included in the model. Results refer to **Yeo’s parcellation. A.** Similarity between models based on diverse stable edges and the full model as a function of behavioral PCs. **B.** Similarity between models based on diverse stable edges and full model as a function of functional connectivity PCs. **C.** Similarity between models based on diverse and uniform stable edges as a function of behavioral PCs. **D.** Similarity between models based on diverse and uniform stable edges as a function of functional connectivity PCs.

### 10.7.1. Control analysis

**Figure S31:**
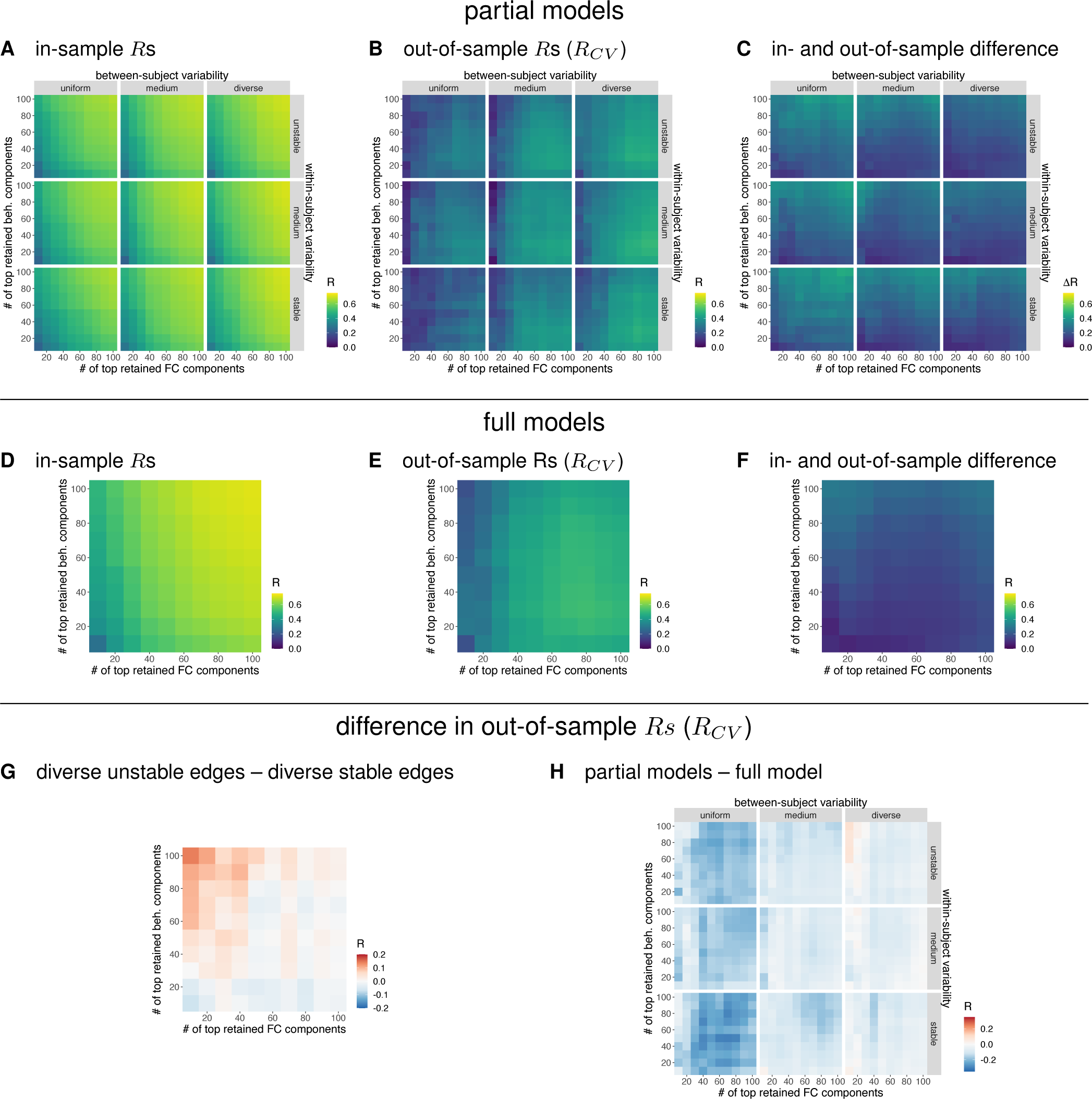
Results of CCA on Glasser’s parcellation. Same as Figure 7, but based on the same number of edges (randomly selected) from each group of edges.

**Figure S32:**
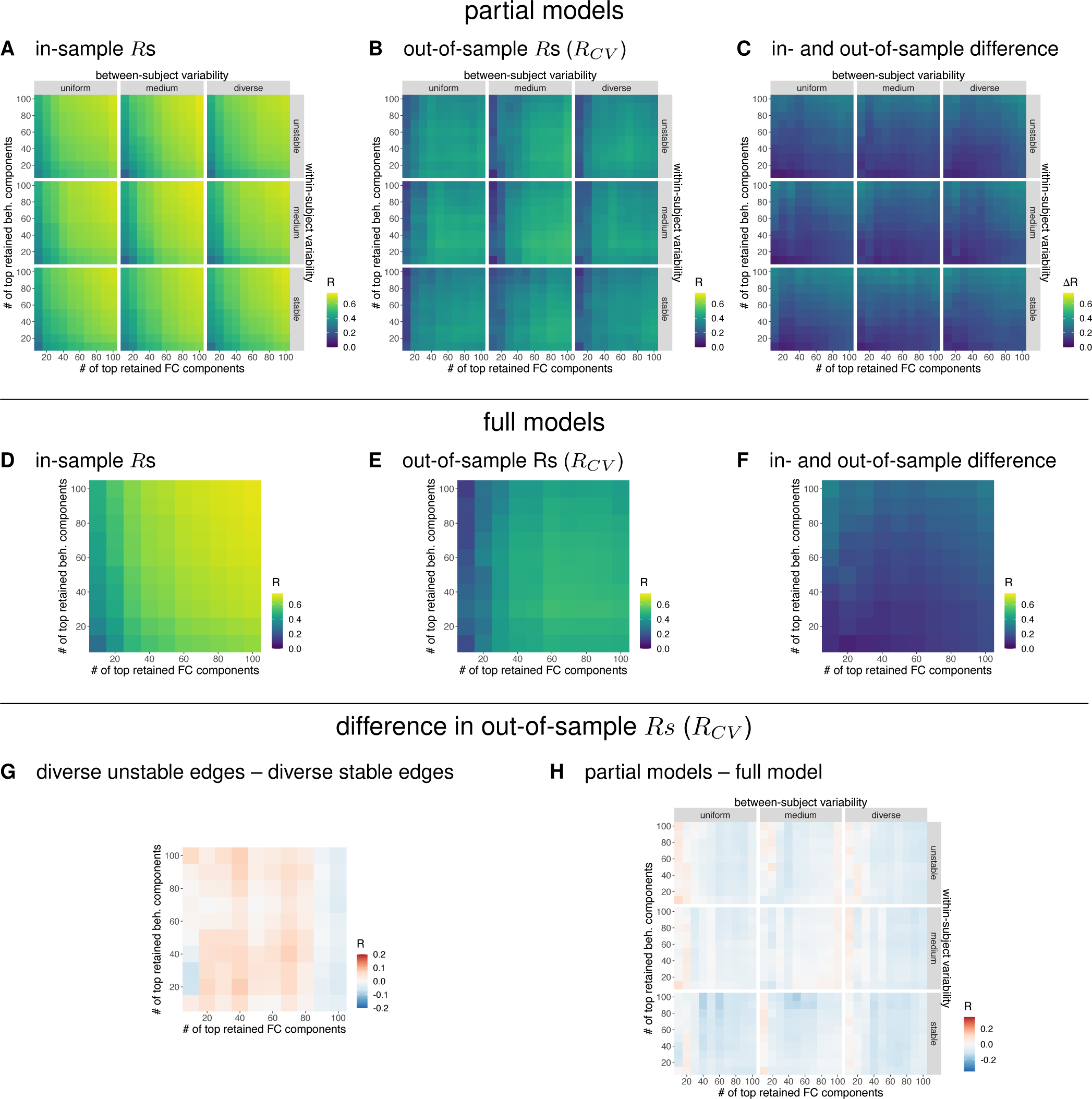
Results of CCA on Schaefer’s parcellation. Same as Figure S26, but based on the same number of edges (randomly selected) from each group of edges.

### 10.8. Simulation

**Figure S33:**
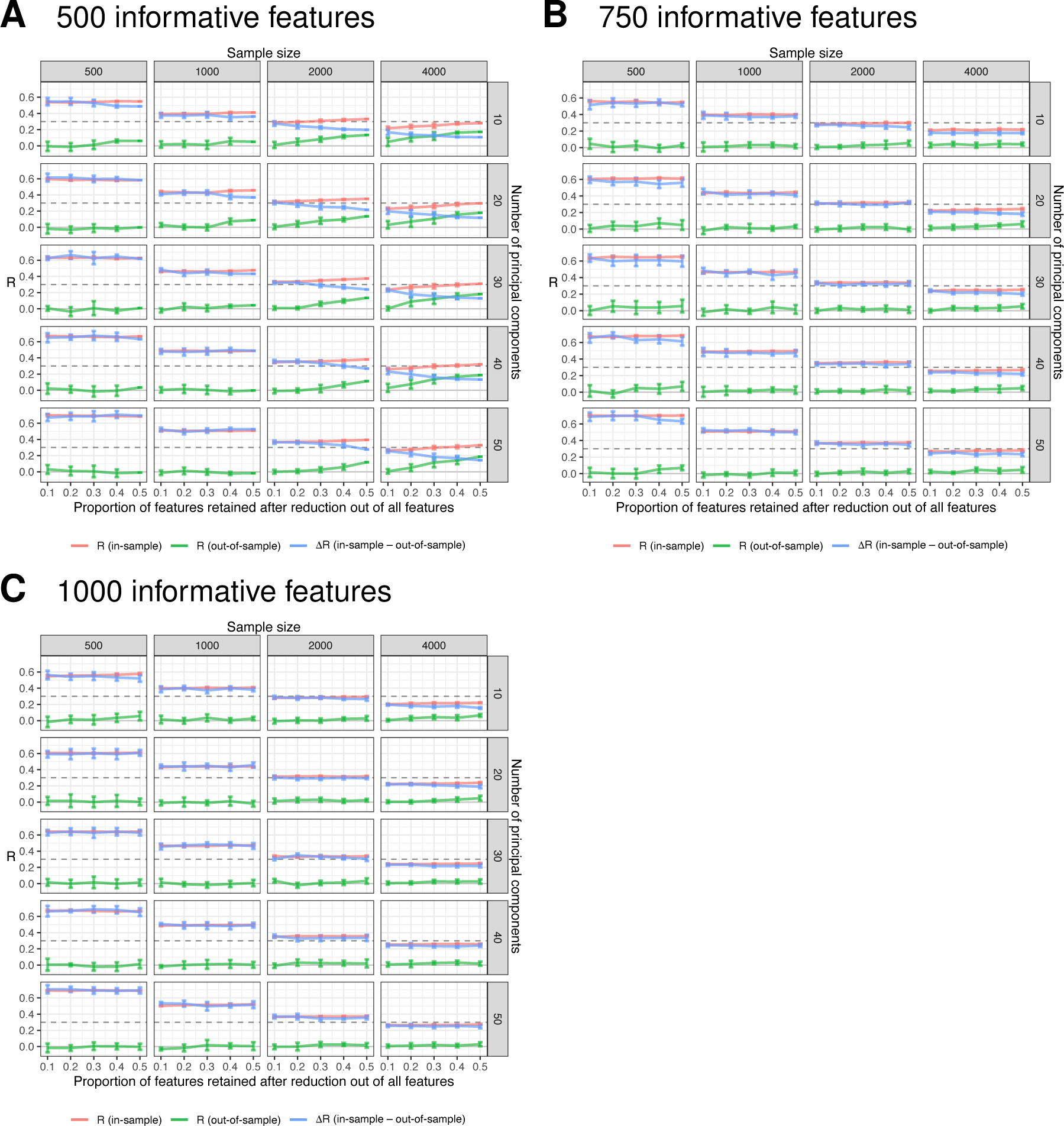
Results of the second simulation of canonical correlation. In this simulation we generated 1000 features with various amounts of informative features (A: 500, B: 750, C: 1000). We combined feature selection with feature reduction (PCA). Note that we also tried to simulate data with 250 informative features, but the simulation could not be completed in time. Dashed line represents true canonical correlation of informative features.

